# Age-related change in task-evoked amygdala—prefrontal circuitry: a multiverse approach with an accelerated longitudinal cohort aged 4-22 years

**DOI:** 10.1101/2021.10.08.463601

**Authors:** Paul Alexander Bloom, Michelle VanTieghem, Laurel Gabard-Durnam, Dylan G. Gee, Jessica Flannery, Christina Caldera, Bonnie Goff, Eva H. Telzer, Kathryn L. Humphreys, Dominic S. Fareri, Mor Shapiro, Sameah Algharazi, Niall Bolger, Mariam Aly, Nim Tottenham

**Author notes:** Please address correspondence to: Paul Alexander Bloom Columbia University Psychology, New York, NY. **Data Availability** Because participants and their parents did not consent to data sharing at the time of participation, we cannot make data from the current study publicly available.

## Abstract

The amygdala and its connections with medial prefrontal cortex (mPFC) play central roles in the development of emotional processes. While several studies have suggested that this circuitry exhibits functional changes across the first two decades of life, findings have been mixed – perhaps resulting from differences in analytic choices across studies. Here we used multiverse analyses to examine the robustness of task-based amygdala–mPFC function findings to analytic choices within the context of an accelerated longitudinal design (4-22 years-old; N=98; 183 scans; 1-3 scans/participant). Participants, recruited from the greater Los Angeles area, completed an event-related emotional face (fear, neutral) task. Parallel analyses varying in preprocessing and modeling choices found that age-related change estimates for amygdala reactivity were more robust than task-evoked amygdala–mPFC functional connectivity to varied analytical choices. Specification curves indicated evidence for age-related decreases in amygdala reactivity to faces, though within-participant changes in amygdala reactivity could not be differentiated from between-participant differences. In contrast, amygdala—mPFC functional connectivity results varied across methods much more, and evidence for age-related change in amygdala–mPFC connectivity was not consistent. Generalized psychophysiological interaction (gPPI) measurements of connectivity were especially sensitive to whether a deconvolution step was applied. Our findings demonstrate the importance of assessing the robustness of findings to analysis choices, although the age-related changes in our current work cannot be overinterpreted given low test-retest reliability. Together, these findings highlight both the challenges in estimating developmental change in longitudinal cohorts and the value of multiverse approaches in developmental neuroimaging for assessing robustness of results. (Preprint: https://www.biorxiv.org/content/10.1101/2021.10.08.463601v1).

**Key Points:**

- Multiverse analyses applied to fMRI data are valuable for determining the robustness of findings to varied analytical choices
- In the current study, age-related change estimates for amygdala reactivity were relatively robust to analytical decisions, though gPPI functional connectivity analyses were much more sensitive, leading some estimates to flip sign
- Both test-retest reliability and robustness to analytical choices are important considerations for developmental research

---

Rodent models and human neuroimaging have provided converging evidence for the importance of the amygdala and medial prefrontal cortex (mPFC) in the development of threat processing (Adolphs, 2008; Forbes et al., 2011), emotion regulation (Pozzi et al., 2020; Silvers et al., 2015; Sullivan & Perry, 2015), and affective learning (Moriceau & Sullivan, 2006; Pattwell et al., 2016). Characterizing growth trajectories of these regions may provide insight into neural constructions underlying emotional development (Meyer & Lee, 2019). To probe amygdala–mPFC circuitry across development, face stimuli are frequently employed because they effectively engage this circuitry while being child-appropriate (Hariri et al., 2002). Though a number of studies have examined age-related changes from childhood to young adulthood in amygdala responses and amygdala–mPFC functional connectivity (FC) associated with emotional face stimuli, findings have varied widely (likely due in part to differences in sample composition and task design; see sTable 1 for details). Several studies have found age-related change in amygdala reactivity, including decreases as a function of age in response to emotional faces (Gee et al., 2013; Guyer et al., 2008; Killgore et al., 2001; Passarotti et al., 2009; Swartz et al., 2014; Telzer et al., 2015) as well as other images (Decety et al., 2012; Silvers et al., 2017b; Vink et al., 2014), increases in amygdala reactivity with age (Joseph et al., 2015; Todd et al., 2011), developmental sex differences (Xu et al., 2021) or peaks during adolescence (Hare et al., 2008; Vijayakumar et al., 2019). Others have observed no age-related changes (Kujawa et al., 2016; Pfeifer et al., 2011; Pine et al., 2001; Wu et al., 2016; Yurgelun-Todd & Killgore, 2006; Zhang et al., 2019).

With task-evoked amygdala–mPFC FC, several studies have found age-related decreases from childhood to young adulthood (Gee et al., 2013; Kujawa et al., 2016; Silvers et al., 2017a; Wu et al., 2016), while others have found increases (Decety et al., 2012; Perlman & Pelphrey, 2011; Vink et al., 2014), developmental sex differences (Xu et al., 2021), or little age-related change (Zhang et al., 2019). While some investigations have found differing age-related change for faces displaying different emotions (Killgore & Yurgelun-Todd, 2007; Swartz et al., 2014; Vijayakumar et al., 2019), even investigations of fearful faces specifically have varied in their developmental findings for both amygdala reactivity and amygdala—mPFC functional connectivity (Forbes et al., 2011; Gee et al., 2013; Killgore et al., 2001; Wu et al., 2016, 2016; Zhang et al., 2019).

While the small sample sizes examined in many studies on amygdala–mPFC development likely contribute to differences in findings (Marek et al., 2020), especially given low reliability of many amygdala—mPFC measures (Elliott et al., 2020; Herting et al., 2017; Sauder et al., 2013), important methodological differences also exist across studies. Differences in age range or sample demographics, stimuli, task (e.g. passive viewing vs. emotion labeling or matching (Lieberman et al., 2007), task design (blocked vs. event-related; Sergerie et al., 2008), or contrast used (faces > shapes vs. faces > baseline) may also contribute to discrepancies (see sTable 1). The brain regions under investigation also differ across studies; for example, prefrontal clusters with which amygdala connectivity was assessed. Interpreting discrepancies across studies without appreciation for these methodological differences would be inappropriate, and in fact, incorrect. Yet, such differences do not account for all discrepancies in findings across studies. Variation in processing pipelines is another source of differences across studies, as varying analytic decisions can produce qualitatively different findings, even between putatively identical analyses of the same dataset (Botvinik-Nezer et al., 2020). Choices including software package (Bowring et al., 2019), spatial smoothing (Jo et al., 2007), treatment of head motion (Achterberg & van der Meulen, 2019), parcellation (Bryce et al., 2021), and functional connectivity approach (Di et al., 2020) can also impact results and qualitatively change findings (Cisler et al., 2014). Additionally, the majority of developmental investigations of amygdala–mPFC function have studied cross-sectional samples. Because cross-sectional studies are susceptible to cohort effects and cannot measure within-participant change, longitudinal work has been recommended for better charting of developmental trajectories (Crone & Elzinga, 2015; Madhyastha et al., 2017).

Here, we used multiverse analyses to examine the robustness of developmental changes to varied analytical decisions. We focused on task-related amygdala–mPFC functional development in an accelerated longitudinal sample ranging from ages 4-22 years. We selected a task that was designed to be appropriate for young ages to characterize developmental change in amygdala–mPFC responses to fear and neutral faces across childhood and adolescence, and we asked whether findings were robust to analytical choices. This accelerated longitudinal design is an extension of the sample reported on by Gee et al. (2013). We preregistered two hypotheses (https://osf.io/8nyj7/) predicting that both amygdala reactivity (1) and amygdala–mPFC connectivity (2) as measured with generalized psychophysiological interaction models (gPPI), would decrease as a function of age during viewing of fearful faces relative to baseline (see Table 1 Aims 1a & 2a).

**Table 1:** Summary of main aims, hypotheses, methods, and findings

| Aim | Preregistered Hypothesis | Analysis Methodology | Key Findings |
| --- | --- | --- | --- |
| 1a. Age-related change in amygdala reactivity to fear faces | Amygdala reactivity to fearful faces will decrease with age, such that younger children will have more positive amygdala reactivity (higher BOLD response to fear faces relative to implicit baseline) than older youth. | Multiverse amygdala ROI (anatomically-defined) analysis using multilevel linear regression at the group level.<br><u>Multiverse decision points:</u><br>Preprocessing software, GLM software, GLM nuisance regressors, amygdala ROI definition, contrast estimate type (t-stat vs. beta estimate), HRF shape, group-level model covariates, exclusion of previously analyzed scans | Across decision points, weak but consistent negative age-related change in amygdala reactivity to fear > baseline and neutral > baseline contrasts<br>No consistent evidence for age-related change in fear > neutral contrast<br>Longitudinal models could identify consistent between-participant differences but not within-participant age-related change |
| 1b. Age-related change in patterns of amygdala responses across task trials | None | Multiverse analysis of slopes of amygdala reactivity across trials, and amygdala reactivity in each half of trials using multilevel linear regression at the group level, single trial models<br><u>Multiverse decision points:</u><br>Global signal subtraction, amygdala ROI definition, group-level model covariates | On average, amygdala reactivity decreased across trials (for both fear and neutral faces)<br>Amygdala reactivity for earlier trials was higher at younger ages<br>Age-related change in amygdala reactivity to fear faces in the first half of trials, but not the second half<br>Similar, but somewhat weaker age-related change for neutral faces |
| 2a. Age-related change in amygdala–mPFC functional connectivity (FC) to fear faces, as measured by generalized psychophysiological interaction (gPPI) | Amygdala–mPFC FC will decrease as a function of age such that as age increases, the valence of FC will shift from positive to negative. | Multiverse gPPI analysis with anatomically defined bilateral amygdala seed and mPFC target ROIs using multilevel linear regression at the group level.<br><u>Multiverse decision points:</u><br>Deconvolution step, mPFC ROI definition, contrast estimate type (t-stat vs. beta estimate), group-level model covariates | No consistent evidence for age-related change in gPPI for any contrast<br>gPPI estimates extremely sensitive to deconvolution step in creation of regressors |
| 2b. Age-related change in amygdala–mPFC functional connectivity to fear faces, as measured by beta-series correlation (BSC) | None for BSC specifically | Multiverse BSC analysis between amygdala and mPFC using multilevel linear regression at the group level.<br><u>Multiverse decision points:</u><br>Global signal subtraction, amygdala ROI definition, mPFC ROI definition, group-level model covariates | No consistent evidence for age-related change in BSC for any condition<br>Amygdala–mPFC BSC was most sensitive to selection of mPFC ROI<br>Global signal subtraction reduced average amygdala–mPFC BSC, but impacts on age-related changes were small<br>BSC estimates were not strongly associated with gPPI estimates |
| 3. Associations of amygdala reactivity, change in amygdala reactivity across trials, or amygdala–mPFC FC with separation anxiety | None | Multiverse multilevel linear regressions with brain measures as predictors for separation anxiety behaviors, controlling for age<br><u>Multiverse decision points:</u><br>Separation anxiety measure, FC measure, mPFC ROI (FC only), amygdala ROI, contrast, deconvolution step (gPPI only) | No evidence that amygdala reactivity, amygdala–mPFC connectivity, or change in amygdala reactivity across trials were associated with separation anxiety behaviors |

Although we did not preregister further hypotheses, we also investigated age-related changes in within-scan differences in amygdala responses across trials and FC using beta series correlations. As previous work identified associations between amygdala–mPFC FC and separation anxiety (Carpenter et al., 2015; Gee et al., 2013), we asked whether any amygdala–mPFC measures were associated longitudinally with separation anxiety behaviors (see Table 1 Aim 3). We used ‘multiverse’ analyses and specification curves to examine the impact of analytical decisions on results. We also investigated test-retest reliability of all brain measurements across longitudinal study visits, given the importance of such reliability for interpreting individual differences or developmental change (Herting et al., 2017). Our multiverse approach allows us to thoroughly explore the robustness of different findings to analytical choices, highlighting the importance of considering both robustness and reliability in developmental research.

## Methods

Before completing analyses, we preregistered methods for the current study through the Open Science Framework at https://osf.io/8nyj7/. Only analyses for age-related changes in amygdala reactivity and amygdala–mPFC gPPI were preregistered in detail (see Table 1 Aims 1a & 2a), and we did not preregister multiverse analyses. Methods detailed below include both information included in the preregistration and additional information and analyses not preregistered. Analysis code & documentation can be found at https://github.com/pab2163/amygdala_mpfc_multiverse.

### Participants

Participants were recruited as part of a larger study examining brain development as a function of early life caregiving experiences. The current sample (N=98; 55 female, 43 male) included typically developing children, adolescents, and young adults covering the ages 4-22 years-old (M = 11.9 years old) who enrolled to participate in a study on emotional development. All participants were reported to be physically and psychiatrically healthy (no medical or psychiatric disorders), as indicated by a telephone screening before participation, and free of MRI contraindications. All except 4 participants fell below clinical cutoffs (see sFigure 2) on the Child Behavior Checklist (CBCL) Total Problems, Internalizing Problems, and Externalizing Problems scales (Achenbach, 1991). The larger study also included youths with a history of institutional and/or foster care outside of the United States, who are not included here. Participants from the greater Los Angeles area were recruited through flyers, state birth records, community events, online advertising, lab website and newsletters, psychologists’ offices, psychology courses at a local university (participants ages 18-22 years old only), and word-of-mouth. Each participant completed up to 3 MRI scans spaced at an average interval of 18 months between visits. Parents provided written consent, children 7+ years old gave written assent, and children under 7 years old gave verbal assent. Study protocols were approved by the local university institutional review board. These data were collected between 2009 and 2015.

An accelerated longitudinal design was used such that participants’ starting ages at scan 1 comprised the entire range of sample ages (4-22 years old), and coverage was approximately balanced across the entire age range (see Figure 1B). The design was structured into 3 study ‘waves’ corresponding with recruitment efforts and visit protocols. Participants were overenrolled at wave 1 to account for anticipated attrition (e.g., braces, relocation, etc) to achieve the desired sample size across the three waves. While most participants were recruited such that their first scan occurred at wave 1, a smaller set of participants were recruited at wave 2, such that some participants completed their first scan at wave 2 (see Figure 1). For these participants, only 2 scans were planned.

**Figure 1.**
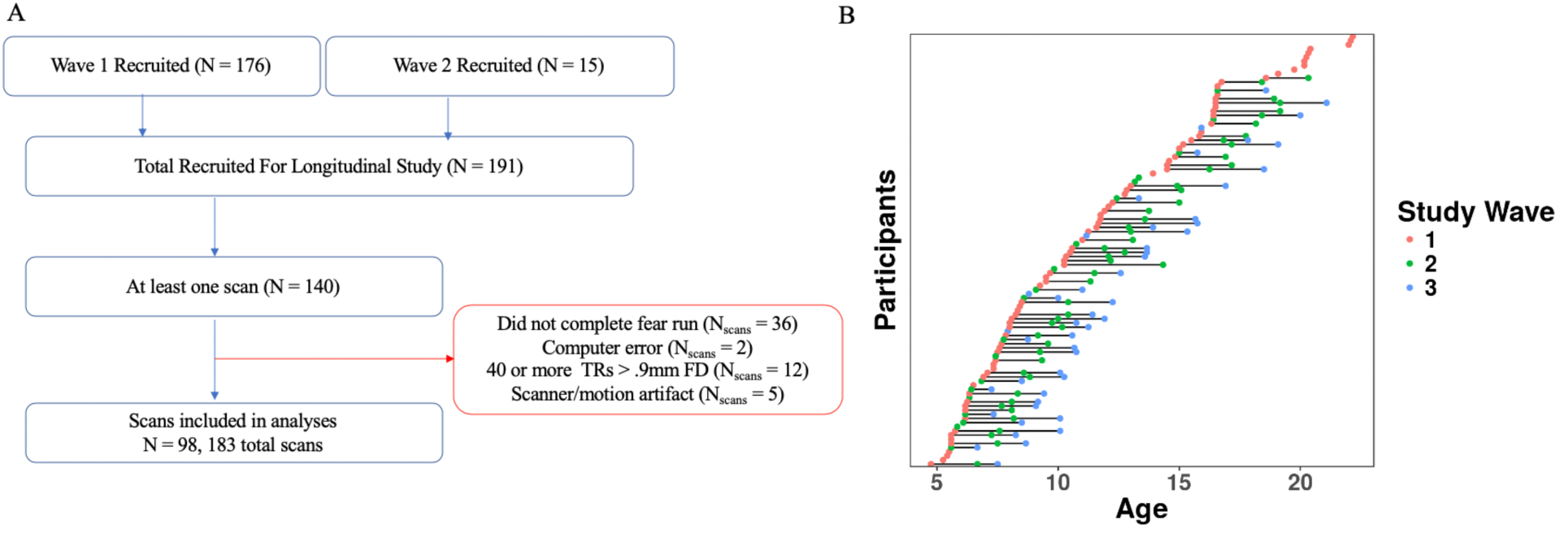
**A.** Schematic showing study inclusion criteria. **B.** Included scans at each study wave, with each dot representing one scan, and horizontal lines connecting participants across study waves.

Of the 191 participants participating in the longitudinal study, 140 completed at least one MRI scan. After exclusions for incomplete task runs (including falling asleep), computer errors resulting in missing stimulus timing files, high head motion, and failed visual QA (scanner/motion artifacts), a final sample of 98 participants (N = 183 total scans) was included for analysis (see Figure 1). Exclusion criteria were preregistered after conducting preliminary preprocessing, but before construction of group-level models and multiverse analysis plans. This sample included 40 participants with 1 scan, 31 with 2 scans, and 27 with 3 scans (one more participant than preregistered due to an initial coding error). Wave 1 data from forty-two of these participants were reported on by Gee et al. (2013).

The median annual household income for participating families was $85,001-$100,000 (for reference, median annual household income in Los Angeles County from 2015-2019 was $68,044; US Census Bureau, 2021). Epidemiological methods were not used to recruit a sample representative of the Los Angeles or United States populations (Heeringa et al., 2004), and Hispanic or Latinx participants were particularly underrepresented. Further sample demographics can be found in the supplementary materials (see sTables 2-3, sFigures 1-2).

### Separation Anxiety

For each participant (except for 10 adults 18-22 years), a parent completed both the Revised Children’s Anxiety and Depression Scale (RCADS-P) and the Screen for Child Anxiety Related Emotional Disorders (SCARED-P) to assess the frequency of symptoms of anxiety and low mood (Birmaher et al., 1999; Chorpita et al., 2000). Following prior work suggesting associations between task-evoked amygdala–mPFC functional connectivity and separation anxiety (Carpenter et al., 2015; Gee et al., 2013), we used the separation anxiety subscales from both the SCARED-P and RCADS-P as measures of anxiety-related behaviors in asking whether such functional connectivity may be linked to anxiety levels during childhood and adolescence. For 11 participants who had missing items on the SCARED-P, indicating parents had skipped or forgotten to answer a question, we imputed responses using 5-Nearest Neighbor imputation using only the other items included in the SCARED-P separation anxiety subscale (Beretta & Santaniello, 2016). As expected, raw separation anxiety scores on both measures decreased as a function of age, while standardized scores (which are normed based on gender and grade level) were consistent across development with few children at or near clinical threshold (see Figure 6).

### Emotion Discrimination Task

Participants completed either two (at wave 3) or three (at waves 1 and 2) runs of a modified ‘go/no-go’ task with emotional faces during fMRI scanning. Runs varied by emotional expression (fear, happy, sad), and within each run participants viewed emotional faces interspersed with neutral faces. To ensure that participants were paying attention, they were asked to press a button whenever they saw a neutral face (no response was required for any other face expression). The order of the runs was counterbalanced across participants; the stimuli within each run were pseudorandomized (Wager & Nichols, 2003) to allow for event-related estimates of the hemodynamic response, and fixed across participants. For the present analysis, only the fear run of the task was used. The other two runs, which used happy and sad faces in place of fear, are not included in the present analysis as these conditions were not present at all waves of data collection. As 50% of trials were ‘go’ trials under this paradigm, we refer to the task as an emotion discrimination task, rather than a true ‘go/no-go’ paradigm since there was no strong prepotent motor response. Stimuli within each run were presented with a jittered ITI (3-10s, Median = 4.93s]) according to a genetic algorithm with a fixation cross on the screen (Wager & Nichols, 2003). Face images were adult White female faces from the Karolinska Directed Emotional Faces database (Calvo & Lundqvist, 2008), and the same face stimuli were used across longitudinal study visits (Vijayakumar et al., 2019). Each run (130 TRs, duration of 4:20) consisted of 48 trials (24 neutral faces, 24 fearful faces), each presented for 350ms. All fMRI analyses of this task used event-related designs.

### MRI Acquisition

Participants under 18-years-old completed a mock scanning session before the MRI scan to acclimate to the scanner environment and practice lying still for data collection. Waves 1 and 2 were collected on a Siemens 3T TIM Trio MRI scanner using a standard radiofrequency head coil. A 2D spin echo image (TR, 4000 ms; TE, 40 ms; matrix size, 256 x 256; 4 mm thick; 0mm gap) was acquired in the oblique plane to guide slice configuration in both structural and functional scans. A whole-brain high-resolution T1-weighted anatomical scan (MPRAGE; 256 x 256 in-plane resolution; 256mm FOV; 192 x 1 mm sagittal slices) was acquired for each participant for registration of functional data. The task was presented through MR-compatible goggles during scanning. T2*-weighted echoplanar images (interleaved slice acquisition) were collected at an oblique angle of ∼30 degrees (130 volumes/run; TR=2000ms; TE=30 ms; flip angle=90°; matrix size=64 x 64; FOV=192 mm; 34 slices; 4 mm slice thickness; skip=0 mm). Wave 3 was collected on a Siemens 3T TIM Trio MRI scanner at a different location using identical acquisition parameters.

### Behavioral Analyses

We used multilevel logistic regression models to estimate age-related changes in several task performance metrics. We fit separate models for the d’ performance metric, overall accuracy (probability of a correct response on any trial), hit rate (on neutral face trials), and false alarm rate (on fear face trials) as the respective outcomes, and included nested random effects for task sessions within participants (models were not nested for d’ as this analysis used only 1 metric per session rather than trial-wise outcomes, but still included random effects for participants). Additionally, to model age-related change in reaction times during correct hit trials, we fit linear, quadratic, cubic, and inverse age (1/age; Luna et al., 2004, 2021) regressions with identical random effects structures. Model equations and results for all behavioral analyses can be found in the supplement (see supplemental methods p.12-14, sFigures 3-4).

### Preregistered fMRI Pipeline

3dskullstrip from the Analysis of Functional NeuroImages (AFNI, v20.1.16) software package (Cox, 1996) was first run on all MPRAGE scans. Next, experimenters checked the quality of the skull stripping. If there were outstanding issues with a particular scan run (areas of brain tissue cut off, or significant areas of skull left in, 30/195 scans), FSL’s brain extraction tool (BET; Jenkinson et al., 2012) was used instead. We used robust brain center estimation, and modified the fractional intensity values between 0.5-0.7 to optimize quality. Slice-time correction was not used. Timeseries of the 6 motion parameters were calculated and subsequent spatial realignment of BOLD volumes was completed using MCFLIRT in FSL (Jenkinson et al., 2002). Scans over a threshold of >40 volumes with > .9mm framewise displacement (framewise displacement calculated as the sum of absolute frame-to-frame differences between head realignment estimates; <u>Power et al., 2012)</u> were excluded from analysis (12 out of an initial 195, or 6.2%). After this exclusion, an average of 96.7% (range = [70.1-100%]) of stimulus-coincident volumes in each scan were below the 0.9mm framewise displacement threshold. The mean age of participants with excluded scans was 7.16 and 8/12 were male. Registration matrices were calculated for registration of functional images to high-resolution structural T1 images using FSL’s FLIRT with boundary-based registration. Registration matrices for standard MNI space were also calculated using both FLIRT (linear registration) and FNIRT (nonlinear registration) with 12 DOF and a warp resolution of 10mm. Data were high-pass filtered at .01Hz and smoothed with an isotropic Gaussian kernel with FWHM of 6mm before running general linear models (GLMs), and 4d volumes were grand mean scaled such that the average intensity value was 10000.

Following preprocessing, we ran scan-level GLMs using FSL’s FEAT (v6.00). We included event-related regressors for fear and neutral faces (convolved with a double-gamma HRF), their temporal derivatives (Pernet, 2014), and 24 head motion nuisance regressors (the 6 head realignment parameters, their temporal derivatives, and their squares (Power et al., 2012). Volumes with FD > .9mm were downweighed to 0 in the GLM. Pre-whitening was used to estimate and remove temporal autocorrelation (Woolrich et al., 2001). For each scan, we calculated fear > baseline, neutral > baseline, and fear > neutral contrasts. We used native-space bilateral amygdala masks generated using Freesurfer (v6.0; Fischl, 2012) by VanTieghem et al. (2021).

### Multiverse Analyses and Specification Curves

In addition to the preregistered pipelines, we conducted multiverse analyses to address all aims in Table 1 and constructed sets of separate specification curves for each aim (see Table 2). In general, multiverse analyses aim to probe the consistency of results across all ‘reasonable’ possible combinations of analysis decisions (i.e. simultaneously taking all possible ‘forking paths’)(Steegen et al., 2016). Because analyzing fMRI data using all reasonable specifications was infeasible (i.e., possibilities are virtually infinite), we took the approach of ‘sampling’ from the many reasonable or commonly-used analysis choices for each multiverse. Despite not being completely comprehensive, this approach still allowed for thorough investigation into the robustness of results. For all multiverse analyses, we constructed specification curves by ranking models by their beta estimates (ascending) for parameters of interest for interpretation and visualization (Cosme & Lopez, 2020; Klapwijk et al., 2019; Orben & Przybylski, 2019; Simonsohn et al., 2015, 2020). Because specification choices were not preregistered, we did not conduct formal null hypothesis testing of specification curves. Instead, as continuous measures of evidence, we report the proportion of specifications resulting in an estimate of the same sign, as well as the proportion of specifications resulting in 95% posterior intervals excluding 0 in the same direction. In addition, to analyze in more detail the impact of specific choices, we submitted point estimates for parameters of interest across all specifications to multiple regression models. From these models, we examined the conditional effects of each analysis decision point on the parameter of interest (see supplemental methods p.30-31, sFigures 11-13, 41-43, & 55-57).

**Table 2:** Summary of forking pipelines used in analyses for each aim^1^

| Aim/Analysis | Decision Point | Choices |
| --- | --- | --- |
| 1a. Age-related change in amygdala reactivity to fear faces > baseline | Preprocessing Software | **FSL FEAT**, C-PAC |
|  | GLM Software | **FSL FEAT**, AFNI 3dDeconvolve |
|  | Hemodynamic Response Function | **Double Gamma**, Single Gamma |
|  | Nuisance Regressors | **24 motion regressors**, 6 motion regressors, 18 motion regressors + WM + CSF |
|  | Low-frequency artifact removal | **High-pass filter (.01Hz)**, Quadratic drift regressor |
|  | First-level GLM Estimates | **Beta Estimates**, T-statistics |
|  | Native vs. Standard MNI Space | **Native Space (Freesurfer)**, Harvard-Oxford Atlas in MNI |
|  | Amygdala ROI | **Bilateral**, Left, Right, High Signal, Low Signal |
|  | Inclusion of 45 previously analyzed scans | **Include**, Exclude |
|  | Outlier treatment | **Exclude +/-3SD from mean**, Exclude +/-3SD from mean + robust regression |
|  | Group-level model covariates | **Mean FD**, Mean FD + run, Mean FD + scanner, Mean FD + run + scanner |
|  | Group-level model quadratic term | Yes, **No** |
|  | Group-level model random slopes | **Yes**, No |
| 1b. Age-related change in patterns of amygdala responses across task trials<br>*FSL preproc & GLM, high-pass filter, 24 motion regressors, 2G HRF, beta estimates, included previously analyzed scans, and robust group-level regression* | Method of quantifying within-scan change | Slopes across trials, trials split into halves, single-trial models |
|  | Global Signal Subtraction | Yes, No |
|  | Amygdala ROI (all MNI space) | Bilateral, Left, Right |
|  | Group-level model covariates | Mean FD, Mean FD + run, Mean FD + scanner, Mean FD + run + scanner |
|  | Group-level model quadratic term | Yes, No |
|  | Group-level model random slopes | Yes, No |
| 2a Age-related change in amygdala–mPFC functional connectivity (FC) to fear faces > baseline, as measured by (gPPI)<br>*FSL preproc & GLM, high-pass filter, 24 motion regressors, 2G HRF, bilateral amygdala ROI in MNI space* | Deconvolution step | Yes, **No** |
|  | mPFC ROI (all MNI space) | **3 different 5mm spheres**, large vmPFC mask |
|  | Outlier treatment | **Exclude +/-3SD from mean**, Exclude +/-3SD from mean + robust regression |
|  | Inclusion of 45 previously analyzed scans | **Include**, Exclude |
|  | Group-level model covariates | **Mean FD**, Mean FD + run, Mean FD + scanner, Mean FD + run + scanner |
|  | Group-level model quadratic term | Yes, **No** |
|  | Group-level model random slopes | **Yes**, No |
| 2b. Age-related change in amygdala–mPFC functional connectivity to fear faces > baseline, as measured by (BSC)<br>*FSL preproc & GLM, high-pass filter, 24 motion regressors, 2G HRF, beta estimates, robust group-level regression, included previously analyzed scans* | Amygdala ROI (all MNI space) | Bilateral, Left, Right |
|  | mPFC ROI (all MNI space) | 3 different 5mm spheres, large vmPFC mask |
|  | Global Signal Subtraction | Yes, No |
|  | Group-level model covariates | Mean FD, Mean FD + run, Mean FD + scanner, Mean FD + run + scanner |
|  | Group-level model quadratic term | Yes, No |
|  | Group-level model random slopes | Yes, No |
| 3. Associations of amygdala reactivity, change in amygdala reactivity across trials, or amygdala–mPFC FC with separation anxiety<br>*See supplemental methods p. 29 for details on included pipelines.* | Brain measure | Amygdala reactivity, amygdala reactivity slopes, amygdala–mPFC gPPI, amygdala–mPFC BSC |
|  | Global Signal Subtraction (amygdala reactivity slopes & BSC only) | Yes, No |
|  | Deconvolution step (gPPI only) | Yes, No |
|  | mPFC ROI (all MNI space, gPPI & BSC only) | 3 different 5mm spheres, large vmPFC mask |
|  | Separation anxiety outcome variable | RCADS, SCARED raw scores, SCARED t-scores |

<sup>1</sup>**Bolded** choices indicate those most closely matching preregistered pipelines.

### Multiverse amygdala reactivity analyses

For amygdala reactivity analyses, we examined the robustness of age-related change estimates to a variety of analytical decisions. In addition to the preregistered FSL-based pipeline, we preprocessed data using C-PAC software (v1.4.1; Craddock et al., 2013). We used C-PAC to take advantage of features supporting running multiple pipeline ‘forks’ in parallel (for example multiple nuisance regression forks using the same registration). No spatial smoothing was used in C-PAC pipelines (see supplemental methods p.14). Following C-PAC and FSL preprocessing, we examined the impact of different sets of commonly-used analysis methods on age-related change in amygdala reactivity. We varied analysis choices of GLM software, hemodynamic response function, nuisance regressors, first-level GLM estimates, amygdala ROI, exclusion criteria (exclude vs. include scans analyzed by Gee et al., 2013), group-level model outlier treatment, and group-level model covariates (see Table 2 & supplemental methods p.14-17). Multiverse analyses of amygdala reactivity included a total of 2808 model specifications (156 ways of defining participant-level amygdala reactivity x 18 group-level model specifications) for each contrast. We analyzed all specifications in parallel. In addition, we examined nonlinear age-related changes using quadratic and inverse age models (see sFigures 14-17) and ran a smaller set of analyses (sFigure 19) to ask whether we could differentiate within-participant change over time from between-participant differences through alternative model parametrization (see supplemental methods p.19).

For all specifications, individual-level amygdala reactivity estimates were submitted to a group-level multilevel regression model for estimation of age-related changes. All models allowed intercepts to vary by participant, and some specifications also allowed for varying slopes (see supplemental methods p.15 for model syntax). All models also included a scan-level covariate for head motion (mean framewise displacement [FD]; Power et al., 2012; Satterthwaite et al., 2012, 2013). Consistent with prior work, head motion was higher on average in younger children, and decreased with age (see sFigure 5), though head motion was not associated with amygdala reactivity estimates for most specifications (see sFigure 26). Age-related change findings examined for the preregistered pipeline also remained consistent under more stringent exclusion thresholds based on mean framewise displacement (see sFigure 27). Across preprocessing specifications, we also examined within-scan similarity of amygdala and whole-brain voxelwise reactivity patterns (see sFigures 20-21) and between-scan correlations of average amygdala reactivity estimates (sFigures 22-24).

### Change in amygdala reactivity across trials

To probe whether amygdala reactivity exhibited within-scan change in an age-dependent manner, we modeled reactivity to each face trial using a Least Squares Separate method (LSS; Abdulrahman & Henson, 2016). After preprocessing, we used FEAT to fit 48 separate GLMs corresponding to each trial in each scan. A given trial was modeled with its own regressor and the remaining 47 trials were modeled with a single regressor. Each GLM also included 24 head motion nuisance regressors and had TRs with framewise displacement > .9mm downweighted to 0. BOLD data were high-pass filtered at .01Hz before the GLM. From each of the 48 GLMs, we extracted the mean amygdala beta estimates corresponding to a contrast for each single trial > baseline.

We constructed separate multiverse analyses using three different methods for measuring change in amygdala reactivity across trials. For method 1 (*slopes*), we measured rank-order correlations between trial number and trial-wise amygdala betas. For method 2 (*trial halves*), we split trials into the first half (trials 1-12) and second half (trials 13-24), and modeled age-related change in each half. For method 3 (*single-trial models*), we constructed larger multilevel models with individual trials as the unit of observation. We conducted several analysis specifications for each method (see Table 2 & supplemental methods p.21-23), and generated corresponding specification curves.

### Multiverse amygdala–mPFC functional connectivity (FC) analyses

We applied multiverse analysis techniques towards examining age-related changes in amygdala–mPFC FC using gPPI and beta-series correlation (BSC) methods. Briefly, gPPI estimates functional connectivity by constructing an interaction term between the timecourse in a seed region of interest and a stimulus (task) regressor. Voxels whose activity is well fit by this interaction term (a psychological-physiological interaction, or PPI) are assumed to be “functionally coupled” with the seed region in a way that depends on the behavioral task (McLaren et al., 2012; O’Reilly et al., 2012). BSC offers a different way of estimating functioning connectivity, by constructing ‘timeseries’ of beta values (i.e., a beta series) in a condition of interest for two regions of interest, and calculating the product-moment correlation between those beta series.

We constructed separate specification curves for age-related change in gPPI and BSC for each contrast. Across gPPI specifications, we varied whether to use a deconvolution step in creating interaction regressors (Di & Biswal, 2017; Gitelman et al., 2003), as well as several other analysis decision points (see Table 2 & supplemental methods p.24-25). The deconvolution step applies to the preprocessed BOLD data from the seed timecourse: these data are first deconvolved to estimate the “underlying neural activity” that produced the BOLD signal (Gitelman et al., 2003), then these deconvolved signals are multiplied with the task regressor (e.g., for fear faces). Finally, this new interaction term is convolved with a hemodynamic response to produce the BOLD functional connectivity regressor of interest. Given recent work indicating that centering the task regressor before creation of the interaction term can mitigate spurious effects (Di et al., 2017), we also compared pipelines in which we centered the task regressor before deconvolution (pipelines including deconvolution in main analyses did not include this step; see sFigure 44).

We preregistered constructing an mPFC ROI containing 120 voxels centered at the peak coordinates reported by Gee at al. (2013) for age-related change in fear > baseline gPPI (Talairach 2,32,8; or MNI 3,35,8). However, after preregistration we discovered that these peak coordinates were not at the center of the ROI reported by Gee at al. (2013), and were quite close to the corpus callosum. The 120-voxel ROI we created that was centered at this peak coordinate would have contained a high proportion of white matter voxels relative to cortical voxels (though this was not true for the mPFC ROI identified by Gee et al. (2013). To address this issue, we instead constructed three spherical ROIs with 5mm radii; the first centered at the above peak coordinates, the second shifted slightly anterior, and the third shifted slightly ventral relative to the second (see Figure 4). Lastly, to examine amygdala functional connectivity with a more broadly-defined mPFC, we also used a ‘large vmPFC’ mask encompassing many of the areas within the ventromedial prefrontal cortex derived from Mackey & Petrides (Mackey & Petrides, 2014).

For BSC analyses, we used beta estimates from the LSS GLMs described above for analyses of within-scan change in amygdala reactivity. Across BSC specifications we varied analyses across several decision points (see Table 2 & supplemental methods p.26), including whether to include a correction for global signal (post-hoc distribution centering(Fox et al., 2009)). We extracted mean beta estimates for amygdala and mPFC ROIs for each trial, then calculated product-moment correlations between the timeseries of betas across trials (neutral and fear separately) for both regions (Di et al., 2020). These correlation coefficients were transformed to z-scores, then submitted to group-level models.

Age-related changes in gPPI and BSC were estimated using multilevel regression models as described for the amygdala reactivity analyses. We focused primarily on linear age-related change, but also examined quadratic and inverse age associations (see sFigures 45-48 & 58-61). We separately examined group mean gPPI and BSC for each contrast (see sFigures 38 & 52), as well as associations between mean framewise displacement and both FC measures across specifications (see sFigures 49 & 62). Additionally, we examined mean estimates and age-related change in ‘task-independent’ FC as measured by beta weight of the ‘physio’ term from the seed amygdala timeseries within the gPPI model (representing baseline amygdala–mPFC functional connectivity controlling for task-induced variance; sFigures 50-51).

### Multiverse analyses of associations between amygdala–mPFC circuitry and separation anxiety behaviors

We used further multiverse analyses to ask whether amygdala reactivity, change in amygdala reactivity over the course of the task, or amygdala–mPFC FC were associated with separation anxiety behaviors. Separate specification curves were created for each brain measure type (amygdala reactivity, amygdala reactivity change across trials, amygdala–mPFC FC). All analyses used multilevel regression models with covariates for age, and specification curves included both RCADS-P and SCARED-P separation anxiety subscales as outcomes (see Table 2 & supplemental methods p.29-30). Because we did not have parent-reported RCADS-P or SCARED-P scores for 10 adult participants, these analyses had an N=173.

### Reliability analyses

To better understand the proportion of variance in each measure explained by the grouping of observations within repeated measurements of the same participants over time, we computed Bayesian intraclass correlation (ICC) estimates through variance decomposition of the posterior predictive distributions of the multilevel regression models previously described. We implemented these through the ‘performance’ R package (Lüdecke et al., 2021; Nakagawa et al., 2017). Negative ICC estimates under this method are possible, and indicate that the posterior predictive distribution has higher variance when not conditioning on random effects than when conditioning on them (likely indicating the posterior predictive variance is large, and random effects explain very little of this variance).

### Model-fitting

All statistical models fit at the group level were run in the R (v 3.6.1) computing environment. In order to most accurately model age-related changes in each of our measures, we attempted to take into account both between-participants information and repeated measurements within participants over time. Unless otherwise indicated, models were estimated using Hamiltonian Markov chain Monte Carlo sampling as implemented in the Stan programming language through the brms package in R (Bürkner, 2019; Gelman et al., 2015). Unless otherwise indicated, all models used package default weakly-informative priors (student-t distributions with mean 0, scale parameter of 10 standardized units, and 3 degrees of freedom for all fixed effects), and were run with 4 chains of 2000 sampling iterations (1000 warmup) each (see supplemental methods p.18-19 and p.30 for syntax).

### Interactive visualizations

Because static plots visualizing the model predictions for all models in each multiverse would require far more page space than available, we created web-based interactive visualization tools for exploring different model specifications and viewing the corresponding raw (participant-level) data and fitted model predictions using R and Shiny (Beeley, 2013). These visualizations can be found at https://pbloom.shinyapps.io/amygdala_mpfc_multiverse/

### Deviations from preregistration

Although we largely completed the preregistered analyses, the current study includes many analyses beyond those proposed in the initial preregistration. Because the additional analyses (i.e., all multiverses) conducted here give us substantial analytical flexibility over that initially indicated by preregistration, we consider all results here to be at least in part exploratory (rather than completely confirmatory), despite the preregistered hypotheses. Additionally, we note that BSC analyses, analyses of change in amygdala reactivity across trials, and analyses of associations between all brain measures and separation anxiety were exploratory, and conducted after we had seen the results of the preregistered reactivity and gPPI analyses. In addition, to avoid possible selection bias introduced by the analytical flexibility inherent in running many parallel analyses, we consider all analysis specifications simultaneously, emphasizing that without further methodological work, we consider all such choices in tandem as providing equal evidential value. While reliability analyses were not preregistered, they too provide key information for interpreting the current analyses.

## Results

### Age-related change in amygdala reactivity

We used multilevel regression models and specification curve analyses to examine age-related changes in amygdala reactivity to faces in an accelerated longitudinal sample ranging from ages 4-22 years (Figure 2). Across specifications, we found relatively consistent evidence for negative age-related change in anatomically-defined (Harvard-Oxford atlas and Freesurfer-defined) amygdala reactivity to fear faces > baseline, such that the vast majority of analysis specifications (99.6%) estimated linear slopes at the group level that were negative in sign, and the majority (60.0%) of 95% posterior intervals about these slopes excluded 0 (Figure 2A; interactive version at https://pbloom.shinyapps.io/amygdala_mpfc_multiverse/). Thus, over half of models indicated that on average, increases in age were associated with decreases in amygdala reactivity to fear faces > baseline. Because the timepoint 1 data in the current study included the 42 scans used by Gee et al. (2013) to age-related changes in amygdala—mPFC circuitry for the fear > baseline contrast, results including these scans may have been more likely to find similar change (particularly for fear > baseline, see sFigures 11-13, & 25). Estimated age-related change was on average weaker, though still largely negative (98.1% negative, 25.3% of posterior intervals excluding 0) when 42 previously analyzed scans (ages 4-17 years) were excluded to provide stricter independence from previously analyzed data (see sFigure 11, Gee et al., 2013). Estimated average age-related change for the fear > baseline contrast was somewhat stronger when using a right amygdala ROI compared to the left amygdala, and when using t-stats extracted from scan-level GLMs rather than beta estimates for group-level models (see sFigure 11).

**Figure 2.**
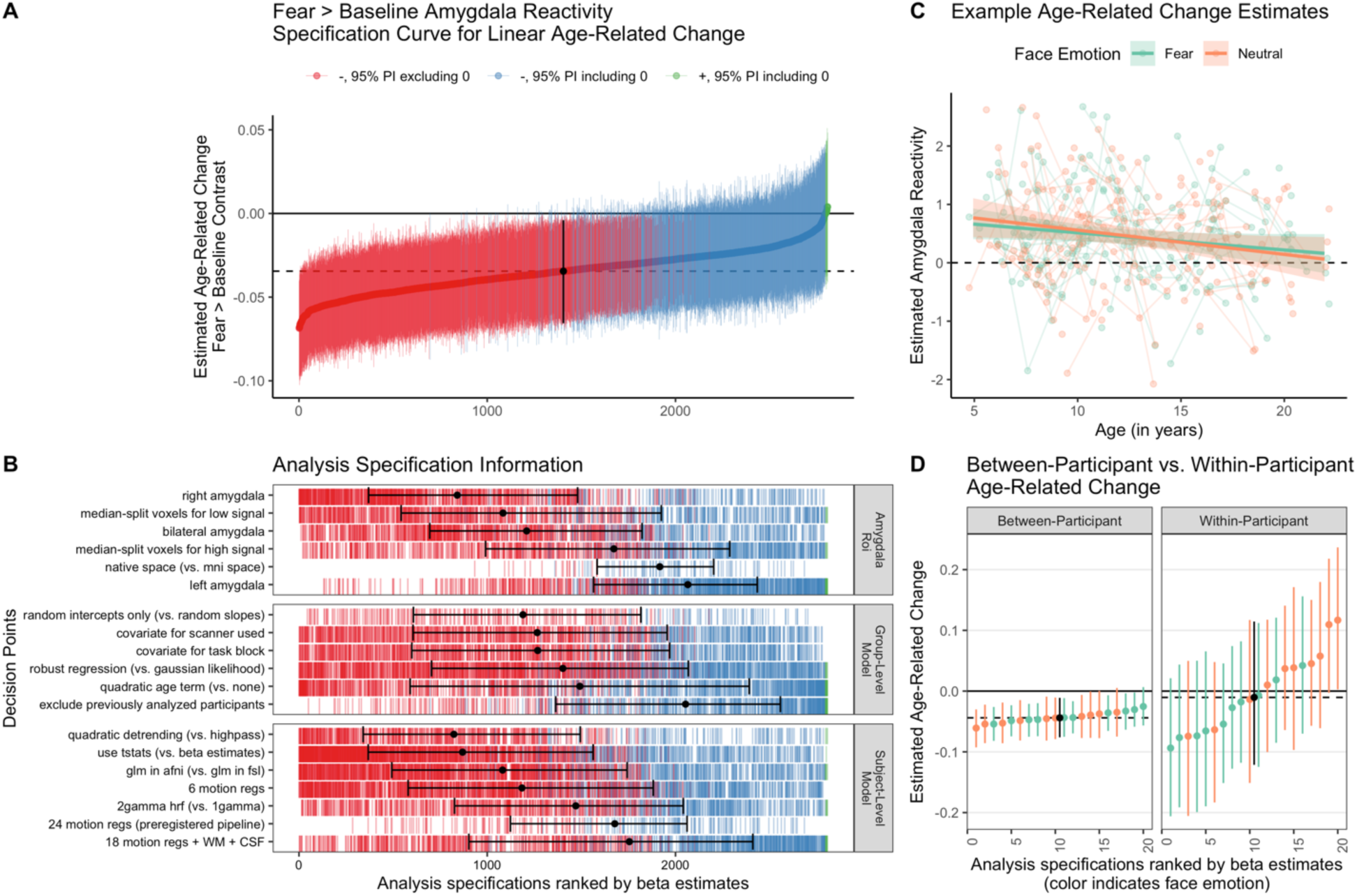
Multiverse analyses of age-related change in amygdala reactivity. **A.** Specification curve of age-related change in fear > baseline amygdala reactivity. Points represent estimated linear age-related change and lines are corresponding 95% posterior intervals (PIs). Models are ordered by age-related change estimates, with the dotted line representing the median estimate across all specifications. Color indicates sign of beta estimates and whether respective posterior intervals include 0 (red = negative excluding 0; blue = negative including 0, green = positive including 0, black = median across all specifications). **B.** Model specification information corresponding to each model in A. Variables on the y-axis represent analysis choices, corresponding color-coded marks indicate that a choice was made, and blank space indicates that the choice was not made in a given analysis. Within each category panel (amygdala ROI, Group-Level Model, and Participant-Level Model), decision points are ordered from top to bottom by the median model rank when the corresponding choice is made (i.e. choices at the top of each panel tend to have more negative age-related change estimates). Black points with error bars represent the median and IQR ranks of specifications making the choice indicated on the corresponding line. **C.** Example participant-level data and model predictions for age-related related change in amygdala reactivity for both the fear > baseline (green) and neutral-baseline (orange) contrasts. Data are shown for a preregistered pipeline using a native space bilateral amygdala mask, 24 motion regressors, t-statistics, high-pass filtering, and participant-level GLMs in FSL. Points represent participant-level estimates, light lines connect estimates from participants with multiple study visits, and dark lines with shaded area represent model predictions and 95% posterior intervals. **D.** Specification curves for a subset of models separately parametrizing within-participant (right) vs. between-participant (left) age-related change for both the fear > baseline (green) and neutral > baseline (orange) contrasts, as well as the median across specifications (black). See https://pbloom.shinyapps.io/amygdala_mpfc_multiverse/ for interactive visualizations.

Parallel multiverse analyses found similarly consistent age-related decreases in neutral faces > baseline amygdala reactivity (see Figure 2C for an example pipeline & sFigure 9 for specification curve), but no consistent evidence for age-related change for the fear > neutral contrast (see sFigure 10). However, there was consistent evidence for higher reactivity for fear faces > neutral on average as well as each emotion compared to baseline (sFigures 6-8), indicating that while the amygdala responses were robust and generally stronger for fear faces compared to neutral, such fear > neutral differences did not change with age. Across contrasts, varying the inclusion of block order or scanner covariates, inclusion of random intercepts, and use of robust regression models had little impact on age-related change estimates (see Figure 2B sFigures 6-8).

While group-level estimates of average age-related change were relatively consistent across specifications, the estimated age terms in these models could be influenced by both within-participant change and between-participant differences (King et al., 2018; Madhyastha et al., 2018). A smaller separate specification curve indicated that when models were parametrized to differentiate within-participant change and between-participant differences, average within-participant change was not consistent across specifications and could not be estimated with precision (Figure 2D). In contrast, estimates of between-participant differences largely indicated negative age-related change in concurrence with our initial model parametrization. At the same time, within-participant versus between-participant terms were not reliably different from one another, indicating that models could not distinguish them despite higher precision for estimating between-participant differences (see sFigure 19). We did not find consistent evidence for quadratic age-related changes in amygdala reactivity (see sFigures 14-17). Inverse age models (i.e. amygdala reactivity modeled as a function of 1/age) indicated results similar to those of linear and quadratic models with most specifications for the fear > baseline and neutral > baseline (though less consistent) contrasts indicating age-related decreases (see sFigure 18).

### Age-related differences in within-scan amygdala reactivity change

To ask whether age-related changes in amygdala reactivity could be due to developmental changes in patterns of amygdala reactivity across face trials (within a run), we examined whether within-scan change in amygdala reactivity varied with age (see Table 1 Aim 1b). Analyses included 42 specifications (3 amygdala regions of interest [ROIs] x 2 global signal correction options x 7 group-level models). Across both fear and neutral trials, linear slopes of amygdala reactivity were negative on average, indicating higher amygdala reactivity at the beginning of the run (Figure 3A, sFigure 30). Across specifications, for both fear (100% of estimates had the same sign, 95.2% of posterior intervals excluding 0 in the same direction) and neutral trials (100% of estimates in the same direction, 38.1% of posterior intervals excluding 0), there was evidence that these within-scan slopes were steeper (i.e., more negative) at younger ages, though evidence was relatively weaker for neutral trials (Figure 3D-E). Specifications with a global signal subtraction step also tended to find stronger age-related change.

**Figure 3.**
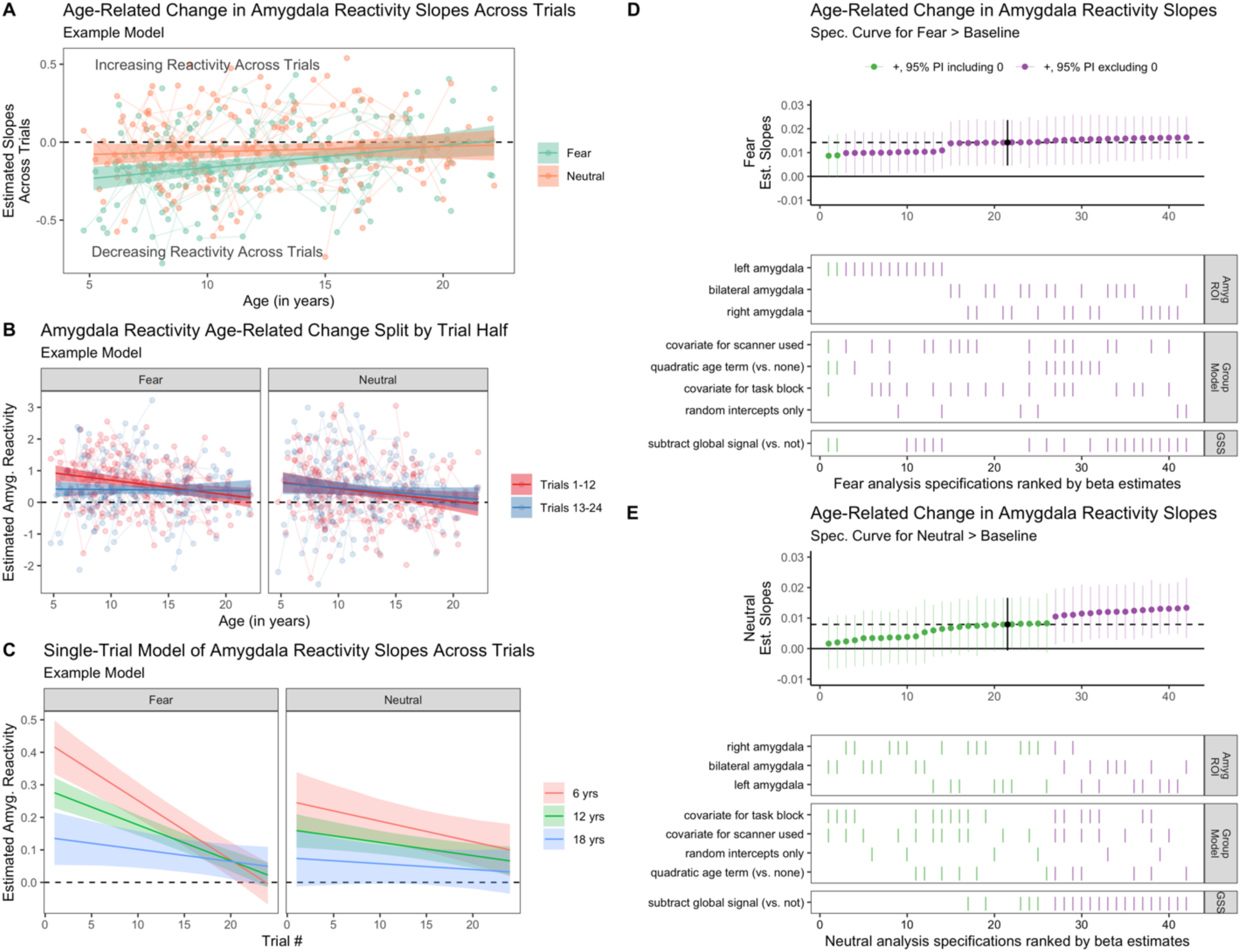
Age-related change in amygdala reactivity across trials. **A.** An example model of estimated age-related change in slopes of beta estimates across both fear (green) and neutral (orange) trials. Negative slopes represent higher amygdala activity in earlier trials relative to later trials. **B.** Example models of estimated age-related change in amygdala reactivity for the fear > baseline (left) and neutral > baseline (right) contrasts for both the first (red) and second (blue) halves of trials. In both A and B, points represent participant-level estimates, light lines connect estimates from participants with multiple study visits, and dark lines with shaded area represent model predictions and 95% posterior intervals. **C.** Example single-trial model predictions of estimated amygdala reactivity for fear (left) and neutral (right) faces as a function of age and trial number. Age was modeled as a continuous variable, and average predictions for participants of age 6 (red), 12 (green) and 18 (blue) years are shown for visualization purposes. All estimates in A-C shown are from an example analysis pipeline using bilateral amygdala estimates and without global signal correction. **D.** Specification curve for age-related change in slopes across fear trials (i.e., many parallel analyses for the fear trials in subplot B). **E.** Specification curve for age-related change in slopes across neutral trials (i.e., neutral trials in plot B). GSS = global signal correction using post-hoc mean centering. For both D and E, color indicates sign of beta estimates and whether respective posterior intervals include 0 (green = positive including 0, purple = positive excluding 0, black = median across all specifications), and horizontal dotted lines represent median estimates across all analysis decisions. Variables on the y-axis represent analysis choices, corresponding color-coded marks indicate that a choice was made, and blank space indicates that the choice was not made in a given analysis.

Similarly, when splitting trials into the first half (trials 1-12) versus second half (trials 13-24), there was consistent evidence (100% of estimates had the same sign, 69.2% with posterior interval excluding 0) for an interaction between age and trial half, such that average reactivity to fear faces > baseline in the first half of trials decreased as a function of age more so than did average reactivity during the second half of trials (see Figure 3B, sFigure 32). This interaction was in the same direction for neutral trials across most specifications (88.5% of estimates), but was typically not as strong (3.8% of posterior intervals excluding 0). Single-trial models indicated similar age-related change in within-scan amygdala dynamics (see Figure 3C, sFigures 33-34). Mean group-level amygdala reactivity was higher for the first half of trials for fear faces > baseline across several specifications, though there were not consistent differences between trial halves for mean amygdala reactivity to neutral faces (sFigure 31).

### Age-related change in task-evoked amygdala–mPFC functional connectivity

We used multilevel regression modelling and specification curve analyses to examine age-related change in task-evoked amygdala–mPFC functional connectivity within the accelerated longitudinal cohort (see Table 1 Aims 2a-b). For the fear > baseline contrast, a specification curve with 288 total specifications (4 definitions of participant-level gPPI estimates x 4 mPFC ROIs x 18 group-level models) of amygdala–mPFC gPPI did not find consistent evidence of age-related change: while 59.0% of models found point estimates in the positive direction, only 23% of posterior intervals excluded 0 (Figure 4C-D, interactive version at https://pbloom.shinyapps.io/amygdala_mpfc_multiverse/). Specification curve analyses found that the sign of the estimated age-related change depended almost entirely on deconvolution, such that most specifications including a deconvolution step resulted in negative age-related change estimates never distinguishable from 0 (78.5% of point estimates negative, 0% of posterior intervals excluding 0), and most specifications not including a deconvolution step resulted in positive age-related change estimates (96.5% of point estimates positive, 47.9% of posterior intervals excluding 0). A visualization of the effects of the deconvolution step on amygdala FC with each of four mPFC ROIs is presented in Figure 4B. While mPFC ROI definition and other analysis decision points also influenced estimates of age-related change in gPPI (Figure 4D), follow-up regression models indicated that the effect of including the deconvolution step was several times larger for the fear > baseline contrast (see sFigures 41-43).

**Figure 4.**
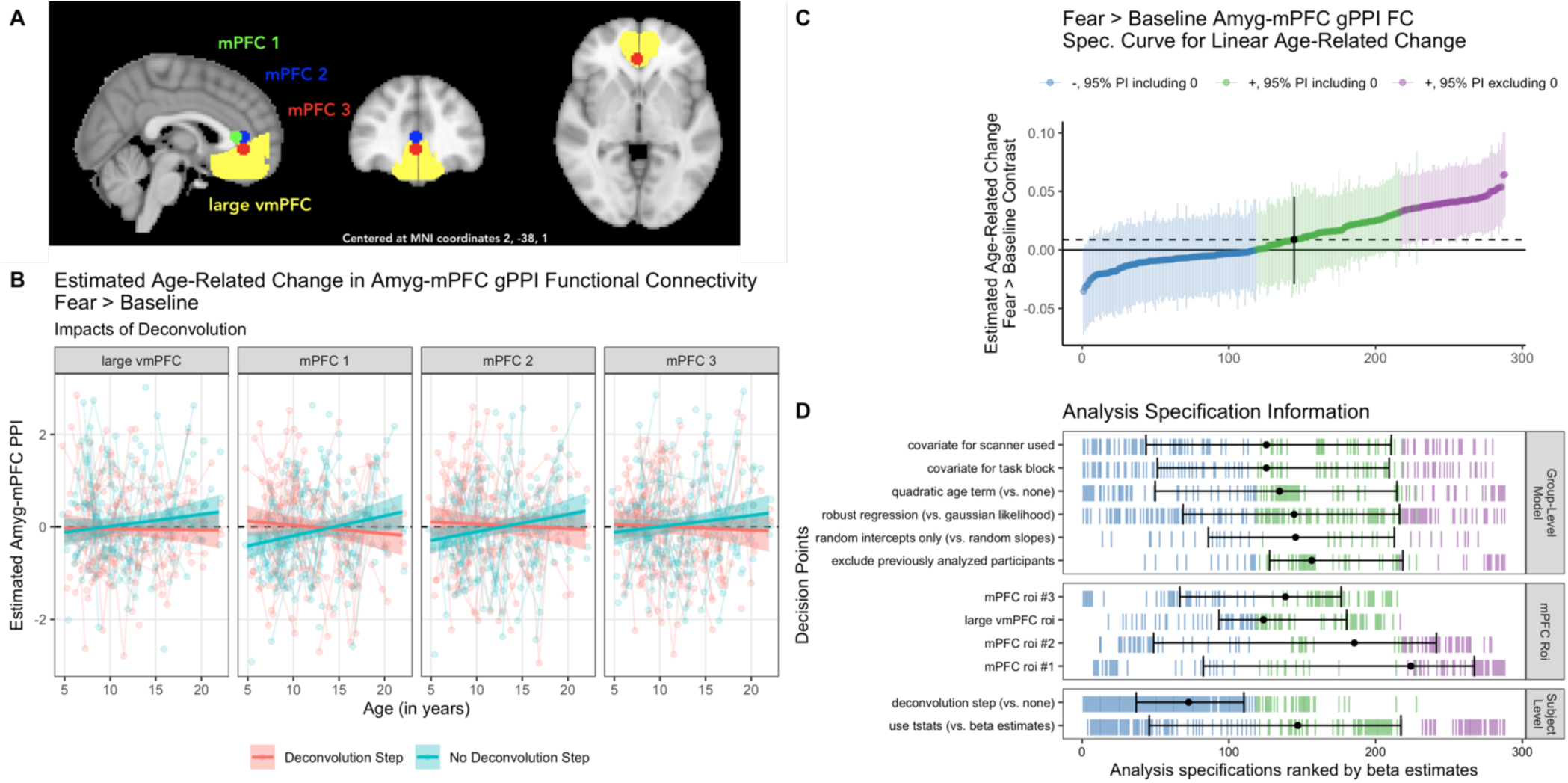
Multiverse analyses of age-related change in amygdala–mPFC connectivity using gPPI methods. **A.** MNI space mPFC ROIs used in connectivity analyses. **B.** Example participant-level data and model predictions for age-related related change in amygdala–mPFC gPPI for analysis pipelines with a deconvolution step (red), or without (blue) for each of the four regions shown in A. Although deconvolution changed the sign of age-related change estimates, the estimates are not ’statistically significant’ for each pipeline alone, except for mPFC ROIs 1 & 2 without deconvolution. **C.** Specification curve of age-related change in fear > baseline amygdala–mPFC gPPI. Points represent estimated linear age-related change and lines are corresponding 95% posterior intervals. Models are ordered by age-related change estimates, and the dotted line represents the median estimate across all specifications. Color indicates sign of beta estimates and whether respective posterior intervals include 0 (blue = negative including 0, green = positive including 0, purple = positive excluding 0, black = median across all specifications). Black points with error bars represent the median and IQR ranks of specifications making the choice indicated on the corresponding line. **D.** Model specification information corresponding to each model in C. Variables on the y-axis represent analysis choices, corresponding color-coded marks indicate that a choice was made, and blank space indicates that the choice was not made in a given analysis. Within each category (Group-Level Model, mPFC ROI, and Participant-Level Model) respectively, decision points are ordered from top to bottom by the median model rank when the corresponding choice is made (i.e., choices at the top of each panel tend to have more negative age-related change estimates). See https://pbloom.shinyapps.io/amygdala_mpfc_multiverse/ for interactive visualizations.

Through equivalent multiverse analyses we also found no evidence of consistent linear age-related change in amygdala–mPFC gPPI for the neutral > baseline and fear > neutral contrasts (see sFigures 39-40), or nonlinear change for any contrast (see sFigures 45-48). In addition, we did not see consistent evidence for group average amygdala–mPFC gPPI for any contrast, though such results often differed as a function of whether a deconvolution step was included (see sFigure 38). Though we included gPPI analysis specifications excluding the 42 scans at timepoint 1 studied by Gee et al. (Gee et al., 2013), exclusion of these scans had little impact on age-related change results (see Figure 4D).

In addition to gPPI analyses, we used beta series correlation (BSC) analyses to examine age-related changes in task-evoked amygdala–mPFC connectivity (see Table 1 Aim 2b). As with gPPI, multiverse analyses of amygdala–mPFC BSC (168 total specifications; 3 amygdala ROI definitions x 4 mPFC ROI definitions x 2 global signal options x 7 group-level models) for fear trials (vs baseline) did not yield strong evidence of age-related change across pipelines (84.5% of point estimates in the same direction, 24.4% of posterior intervals excluding 0; Figure 5A, interactive version at https://pbloom.shinyapps.io/amygdala_mpfc_multiverse). Unlike gPPI analyses, however, choice of mPFC ROI (as well as amygdala ROI, though this was not examined for gPPI) most impacted age-related change in BSC estimates, rather than preprocessing or modeling analytical choices (Figure 5B, sFigures 55-57). Accordingly, while global signal subtraction resulted in weaker amygdala–mPFC BSC on average (see sFigure 52), inclusion of this step did not consistently affect age-related change estimates (Figure 4C). We did not find consistent evidence for age-related change in amygdala–mPFC BSC for neutral trials (vs baseline), or for fear > neutral trials (sFigures 53-54). We did not find consistent evidence for nonlinear age-related change for any contrast (sFigures 58-61).

**Figure 5.**
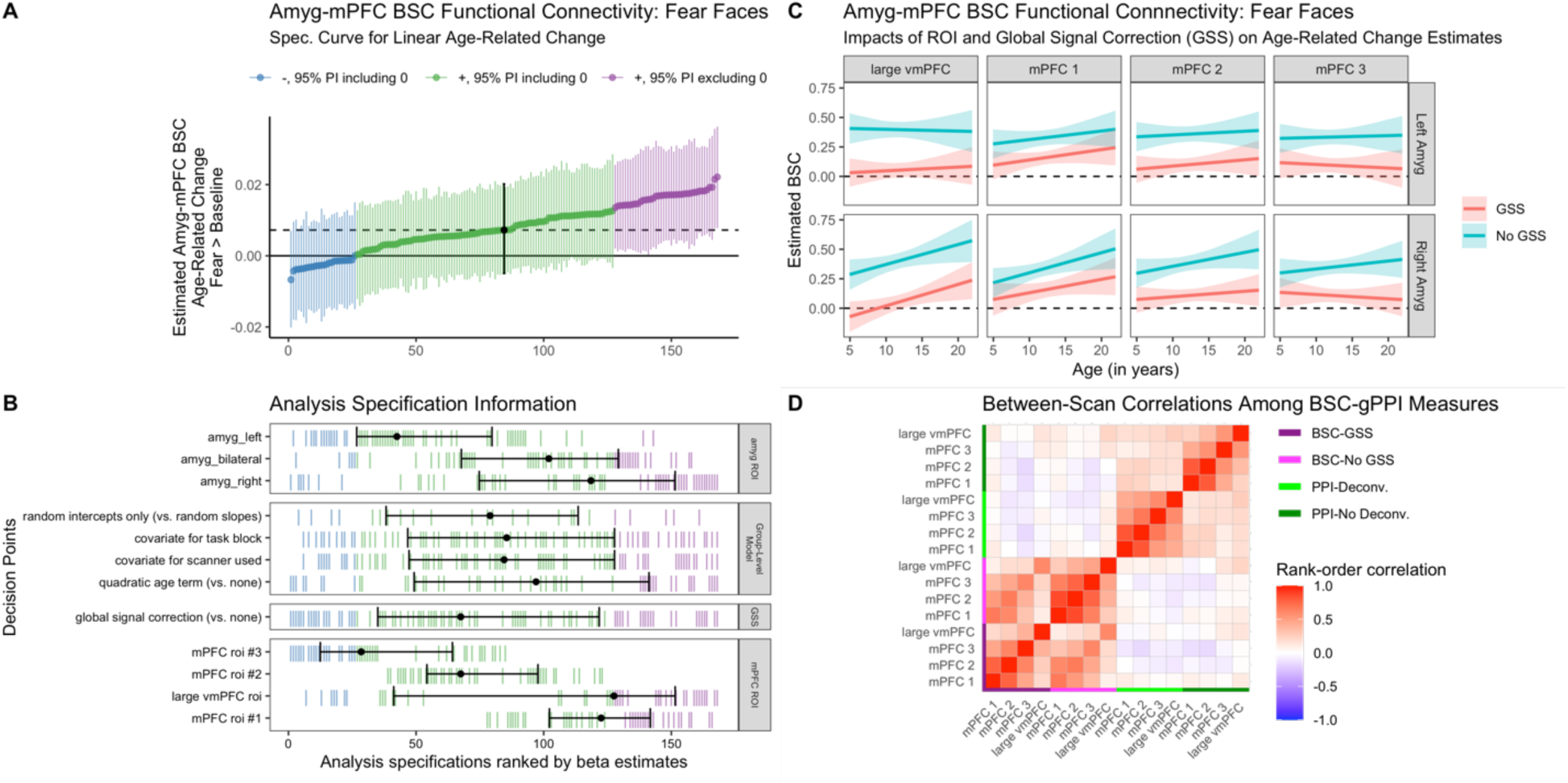
Multiverse analyses of age-related change in amygdala–mPFC connectivity using beta-series correlation (BSC) methods. **A.** Specification curve of age-related change in amygdala–mPFC BSC for fear trials. Points represent estimated linear age-related change and lines are corresponding 95% posterior intervals. Models are ordered by age-related change estimates, and the dotted line represents the median estimate across all specifications. Color indicates sign of beta estimates and whether respective posterior intervals include 0 (blue = negative including 0, green = positive including 0, purple = positive excluding 0, black = median across all specifications). **B.** Model specification information corresponding to each model in A. Variables on the y-axis represent analysis choices, corresponding color-coded marks indicate that a choice was made, and blank space indicates that the choice was not made in a given analysis. Within each category (amygdala ROI, group-level model, global signal subtraction, and mPFC ROI) respectively, decision points are ordered from top to bottom by the median model rank when the corresponding choice is made (i.e., choices at the top of each panel tend to have more negative age-related change estimates). Black points with error bars represent the median and IQR ranks of specifications making the choice indicated on the corresponding line. GSS = global signal correction using post-hoc mean centering. **C.** Example model predictions for age-related change in amygdala–mPFC BSC for fear trials for analysis pipelines with a global signal subtraction (GSS, post-hoc mean centering) step (red), or without (blue) for each of the four mPFC regions (see Figure 4A) with the left and right amygdala. Pipelines shown have random slopes, no covariates for task block or scanner, and no quadratic age term. **D.** Between-scan rank-order correlations between amygdala–mPFC connectivity measures. All gPPI measures are for the fear > baseline contrast, and BSC measures are for fear trials. See https://pbloom.shinyapps.io/amygdala_mpfc_multiverse/ for interactive visualizations.

Additionally, we constructed a correlation matrix using rank-order correlations of scan-level BSC and gPPI estimates for the fear (vs baseline) condition. Across scans, there was little evidence of correspondence between BSC and gPPI metrics for amygdala–mPFC connectivity (Figure 5D, sFigures 63-66). Further, FC estimates tended to be positively correlated within a method type (BSC, gPPI) across mPFC ROIs, though less strongly for gPPI estimates with versus without a deconvolution step.

In addition to gPPI and BSC methods for functional connectivity, we also explored between-scan associations between amygdala reactivity and mPFC reactivity (sFigures 28-29). Multilevel models indicated that amygdala reactivity for fear faces > baseline was positively associated with mPFC reactivity for fear faces > baseline for all mPFC ROIs, though we did not find consistent evidence for age-related changes in associations between amygdala and mPFC reactivity to fear faces > baseline (see sFigure 29).

### Amygdala–mPFC Measures & Separation Anxiety

We conducted multiverse analyses of associations between several amygdala–mPFC measures (amygdala reactivity, amygdala–mPFC FC, within-scan changes in amygdala reactivity) and separation anxiety behaviors (see Table 1 Aim 3). Separation anxiety behaviors on average decreased with age, as indicated by the RCADS-P and SCARED-P raw scores (Figure 6A-C). Neither specification curves for amygdala reactivity (18 total specifications, 56% of point estimates in the same direction as median estimate, 0% of posterior intervals excluding 0), amygdala–mPFC gPPI FC (90 total specifications, 72% of point estimates in the same direction as median estimate, 1% of posterior intervals excluding 0), amygdala–mPFC BSC FC (18 total specifications, 83% of point estimates in the same direction as median estimate, 0% of posterior intervals excluding 0), nor slope of amygdala responses across trials (12 total specifications, 75% of point estimates in the same direction as median estimate, 17% of posterior intervals excluding 0), found consistent evidence for associations between brain measures and separation anxiety. Similar specification curves found little consistent evidence for associations between brain measures and generalized anxiety, social anxiety, or total anxiety behaviors (see sFigure 67). All specifications controlled for age (see supplemental methods p.30).

**Figure 6.**
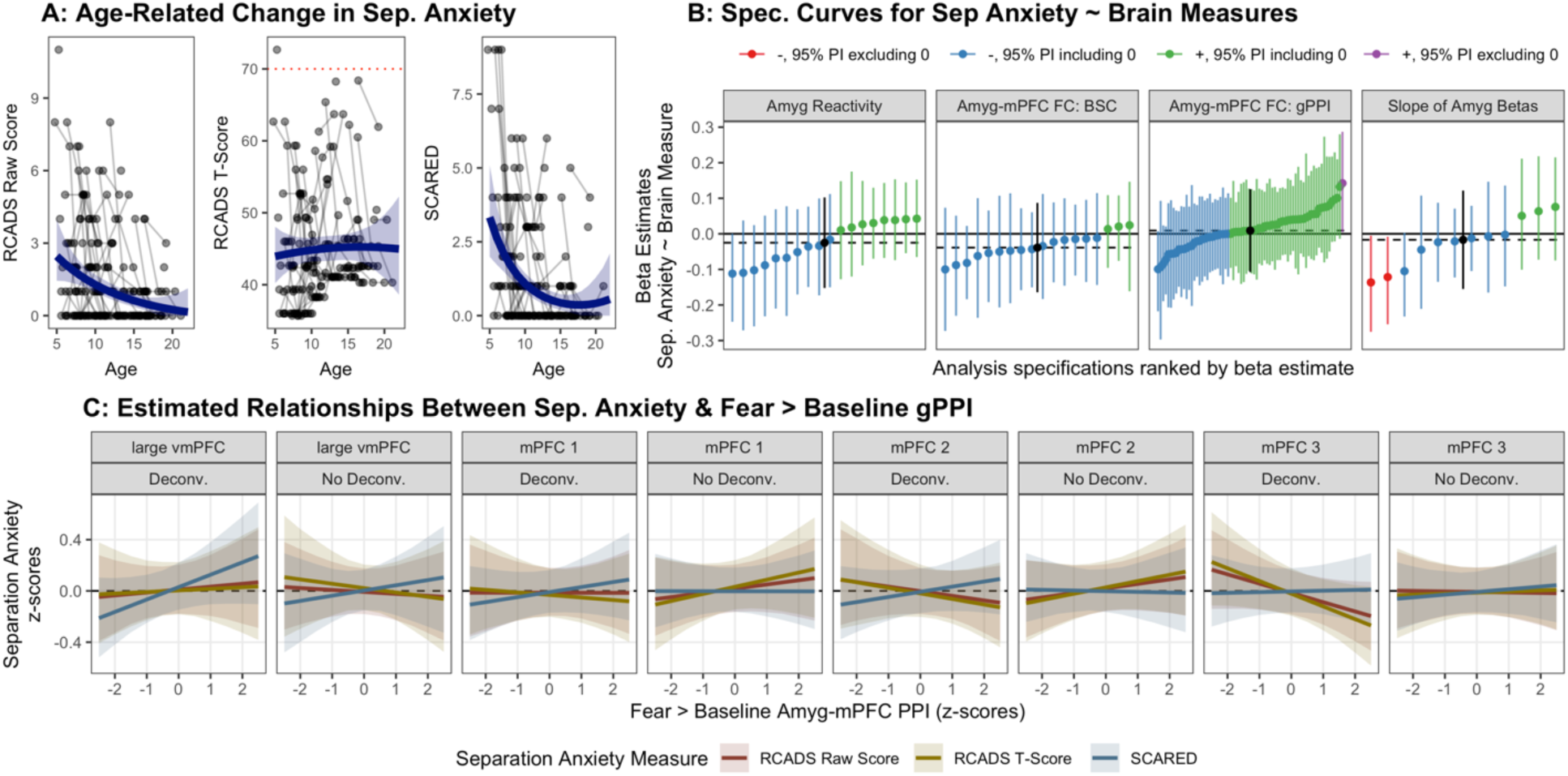
Multiverse analyses of associations between amygdala–mPFC circuitry and separation anxiety. **A.** Age-related change in SCARED and RCADS raw and t-scores for parent-reported separation anxiety subscales. The red dotted line in the middle panel represents the clinical threshold for the standardized RCADS measure (because this T-score measure is standardized based on age and gender, no age-related change is expected). **B.** Separate specification curves for associations of amygdala reactivity (left), amygdala–mPFC connectivity (both gPPI and BSC; center two panels), and amygdala reactivity slopes across trials (right) with the three separation anxiety outcomes shown in A. Points represent estimated associations between brain measures and separation anxiety (controlling for mean framewise displacement and age) and lines are corresponding 95% posterior intervals. Models are ordered by beta estimates, and the dotted line represents the median estimate across all specifications. Color indicates sign of beta estimates and whether respective posterior intervals include 0 (red = negative excluding 0, blue = negative including 0, green = positive including 0). Scores on each separation anxiety outcome were z-scored for comparison. **C.** Example model predictions for associations between fear > baseline amygdala–mPFC gPPI and each separation anxiety measure. Predictions and 95% posterior intervals are plotted for each separation anxiety measure separately for each mPFC region, and for gPPI pipelines with and without a deconvolution step. Pipelines shown use robust regression, have random slopes, no covariates for task block or scanner, and no quadratic age term.

To more specifically follow up on previous work reporting associations between separation anxiety behaviors and amygdala–mPFC gPPI for fear > baseline specifically (Gee et al., 2013), we plotted model predictions for such models from the above multiverse analysis for each of the four mPFC ROIs, across all three separation anxiety outcome measures, and both with and without a deconvolution step (Figure 6E). We did not find consistent evidence for associations with separation anxiety, and results showed high sensitivity to the deconvolution step, mPFC ROI, and outcome measure used.

### Reliability

To examine test-retest reliability estimates of amygdala—mPFC measures across longitudinal visits, we computed Bayesian ICC estimates using a variance decomposition method (Lüdecke et al., 2021). Because such models can accommodate missing data, all observations (98 participants, 183 total scans) were used, including participants with only 1 visit. All amygdala reactivity (Figure 7A) and amygdala—mPFC functional connectivity (Figure 7C) measures, as well as slopes of amygdala reactivity estimates across trials (Figure 7B), demonstrated poor reliability (ICC < 0.4; Cicchetti & Sparrow, 1981; Elliott et al., 2020). Separation anxiety measures demonstrated somewhat higher, though still largely poor reliability (point estimates ∼0.4, 95% CIs included values below 0.4; Figure 7D). Head motion in the scanner (mean framewise displacement) showed the highest reliability (ICC = 0.52, 95% CI [0.29, 0.68]),

**Figure 7.**
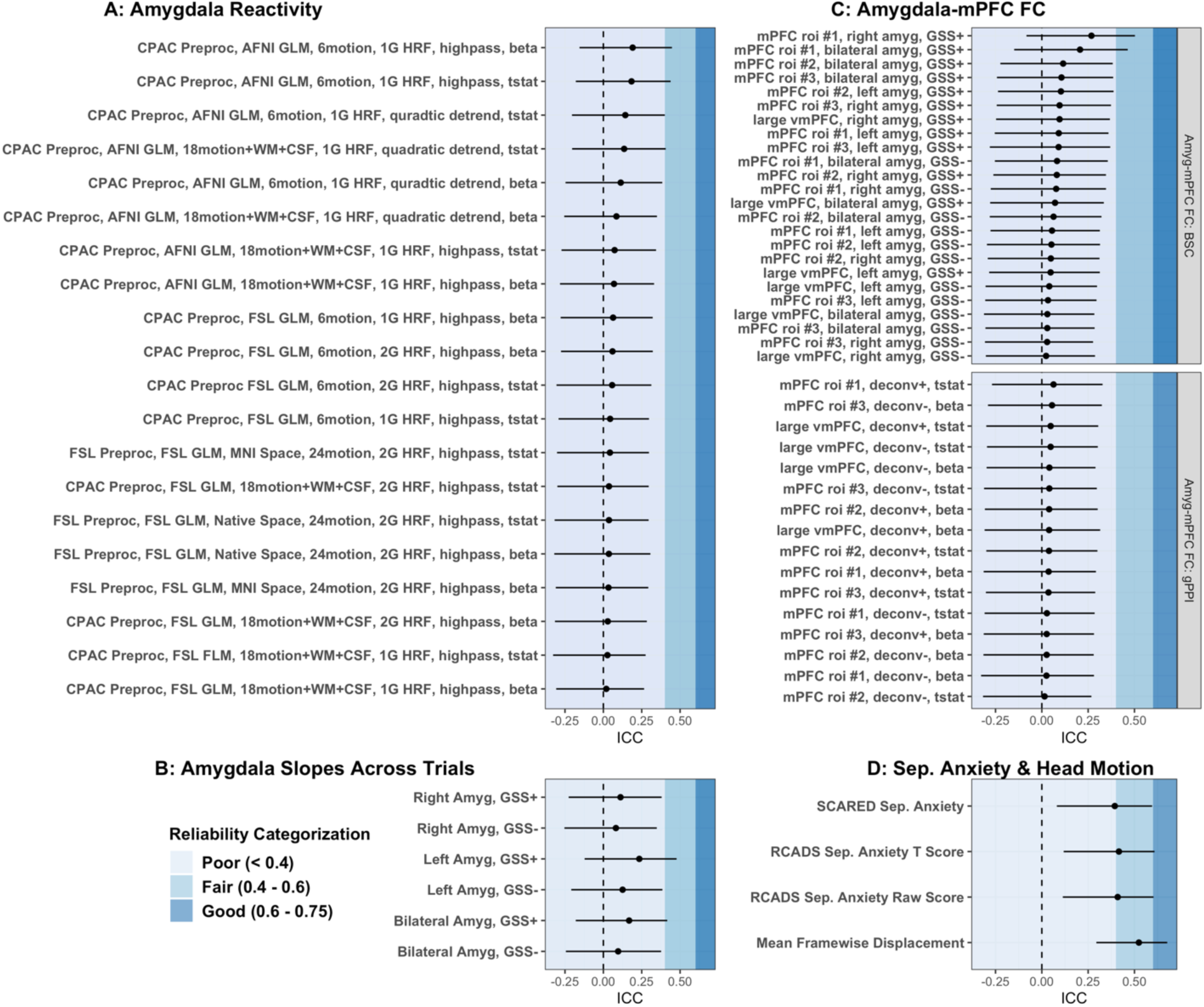
Longitudinal test-retest Bayesian ICC estimates. ICC values are shown for amygdala reactivity (A), slopes of amygdala reactivity betas across trials (B), amygdala—mPFC functional connectivity using both gPPI and BSC methods (C), and separation anxiety and in-scanner head motion measurements (D). Shaded background colors depict whether ICC estimates are categorized as poor (< 0.4), fair (0.4 - 0.6), or good (0.6 – 0.75) reliability. No ICC estimates met the threshold for excellent reliability (>0.75). Bayesian ICC estimates were calculated through a variance decomposition based on posterior predictive distributions. Negative values indicate higher posterior predictive variances not conditioned on random effect terms than conditioned on random effects terms.

### Discussion

Measures that are both robust to researcher decisions and reliable across measurement instances are critical for studies of the human brain (Botvinik-Nezer et al., 2020; Bowring et al., 2019; Elliott et al., 2020; T. Xu et al., 2022). The accelerated longitudinal design and multiverse analysis approach used in the current study allowed a rare opportunity to examine both reliability and robustness of amygdala—mPFC measures using a rapid event-related face task from early childhood through young adulthood. Overall, estimates for age-related change in amygdala reactivity were relatively robust to a variety of analytical decision points, while age-related change estimates for amygdala—mPFC connectivity were more sensitive to researcher choices. gPPI analyses were particularly sensitive to whether a deconvolution step was applied. Yet, in concurrence with previous work (Elliott et al., 2020; Haller et al., 2022; Herting et al., 2017; Infantolino et al., 2018; Kennedy et al., 2021; Nord et al., 2017; Sauder et al., 2013), amygdala—mPFC measures displayed consistently poor test-retest reliability across many analytical specifications. While low reliability estimates in the present study may be due in part to the long (∼18 months) test-retest interval (Elliott et al., 2020) and potential true developmental change (Herting et al., 2017), low reliability nevertheless imposes a major caveat towards interpretation of the current developmental findings.

The present findings are valuable from a methodological standpoint in evaluating the robustness of analytical tools used. A measurement can have high test-retest reliability yet low robustness (high sensitivity) to analytical decisions, or vice versa (Li et al., 2021). Because neither robustness nor reliability guarantee the other, current findings on the impacts of analytic choices will likely be informative in guiding future studies. Thus, we discuss each of the main analyses below, with particular emphasis on how findings are impacted by analytic choices.

### Amygdala reactivity

While there were differences across model specifications, the majority of pipelines supported our hypothesis that amygdala reactivity to fearful faces decreases with age from early childhood through early adulthood (see Table 1 Aim 1a). Across specifications, we found relatively robust evidence for age-related decreases in amygdala reactivity to both fearful and neutral faces (Figure 2A). Yet, findings also varied considerably across specifications. For example, only 60% of pipelines produced results that would be individually labeled as ‘significant’ (under *α* = .05), indicating that multiple investigations of this dataset could likely lead to qualitatively different conclusions. While over half of analyses found evidence consistent with studies indicating greater amygdala reactivity to fear faces > baseline in younger children (Forbes et al., 2011; Gee et al., 2013; Guyer et al., 2008; Swartz et al., 2014), the other 40% of specifications would have been consistent with investigations that found little age-related change (NB: there were also differences in samples, age ranges, task parameters, and behavioral demands across these studies; Kujawa et al., 2016; Wu et al., 2016; Zhang et al., 2019). We also found that different specifications resulted in somewhat different nonlinear trajectories (see sFigures 14-18). Not only did inverse age and quadratic age models find different trajectories (as would be expected), but quadratic trajectories themselves also displayed considerable analytic variability, with some specifications finding “convex” and others finding “concave” fits (see sFigure 17). Although estimating nonlinear age-related change was not a primary goal of the present study, future work should use model comparisons for better differentiating nonlinear patterns (Curran et al., 2010; Luna et al., 2021).

Models also found evidence for between-participant differences, but could neither identify within-participant change (Figure 2D) nor differentiate between-participant from within-participant estimates. As such, interpretation of the age-related change reported here is subject to many of the same limitations that apply to cross-sectional designs (Glenn, 2003), where age-related changes may not necessarily indicate ‘true’ developmental growth. High uncertainty in estimating average within-participant change could be driven by several factors, including true heterogeneity in individual trajectories, low measurement reliability, scanner differences across longitudinal timepoints, or unmodeled variables impacting amygdala reactivity. Additionally, the within-participants terms represent a smaller age range (a maximum of 4 years for any given participant), relative to the broader age range assessed by the between-participants terms (18 years), which may have placed additional limits on identifying reliable within-participant change.

Age-related change in amygdala responses to fear faces over baseline seemed largely the result of earlier trials in the task (see sFigures 32-34). While differences in task design and contrast across studies have been highlighted as potential sources of discrepant findings on the development of amygdala function (Killgore & Yurgelun-Todd, 2007; Lieberman et al., 2007; Swartz et al., 2014), this result indicates that attention to trial structure and task duration may also be necessary in comparing studies. Because the paradigm used in the current study involved a task requiring participants to press for one face (‘neutral’) and not press for ‘fear’ faces, findings specific to fear faces over baseline under the current paradigm may also be driven by behavioral task demands.

### Amygdala–mPFC Functional Connectivity

We did not find evidence for our second hypothesis, as neither gPPI nor BSC analyses indicated consistent evidence of age-related change in amygdala–mPFC functional connectivity (see Table 1 Aims 2a-2b, Figures 4-5). Thus, the age-related changes in task-evoked amygdala–mPFC connectivity identified in prior work (Gee et al., 2013; Kujawa et al., 2016; Wu et al., 2016) was not identified here, consistent with (Zhang et al., 2019). Crucially, however, our specification curves did not find strong evidence *against* such age-related change, as we did not observe precise and consistent ‘null’ estimates across specifications. Additionally, quadratic and inverse age models did not find consistent evidence for nonlinear age-related change (see sFigures 45-48 & 58-61).

gPPI results were sensitive to whether a deconvolution step had been included in the preprocessing pipeline, such that we mostly found age-related decreases in amygdala–mPFC connectivity with a deconvolution step included, and age-related increases without it (although most pipelines would not have been ‘statistically significant’ on their own, see Figure 4B). While deconvolution has been argued to be a necessary step for event-related PPI analyses (Gitelman et al., 2003), recent work has shifted guidelines on its use, and it may not be recommended for block designs (Di et al., 2020; Di & Biswal, 2017). Because the true ‘neuronal’ signal underlying the BOLD timeseries within a given ROI cannot be directly measured, deconvolution algorithms are difficult to validate. Further, deconvolution may cause PPI results to be driven by baseline connectivity if task regressors are not centered (Di et al., 2017), although such centering did not have a major influence on age-related change results in the present analyses (see sFigure 44). Within the current study, small tweaks to AFNI’s 3dTfitter algorithm for deconvolution resulted in vastly different regressors (see sFigure 36), suggesting the potential for high analytic variability even between gPPI analyses ostensibly using deconvolution. While the present study does not provide evidence that can inform whether or not deconvolution is recommended, further work is needed to optimize and validate applications of gPPI methods and selection of appropriate task designs. gPPI may be better equipped for block-designs and particularly ill-posed for rapid event-related tasks due to both difficulties in resolving which times within the BOLD timeseries reflect functional connectivity evoked by rapid (350ms) events and low statistical power in estimating such task-evoked connectivity <u>(see sFigures 35-37; O’Reilly et al., 2012)</u>. Concurrent with previous work, beta series correlation analyses may have higher statistical power for identifying task-related connectivity signal than gPPI within event-related designs more generally (Cisler et al., 2014).

Age-related change estimates for amygdala—mPFC BSC showed somewhat higher robustness to analytic decisions compared to gPPI. For BSC analyses, choice of mPFC ROI contributed most to variability in age-related change estimates (see Figure 5B, sFigures 55-57). While a global signal correction (post-hoc distribution centering) greatly decreased *average* amygdala—mPFC BSC connectivity (see Figure 5D, sFigure 52) for both fear and neutral faces, this analytical step did not impact age-related change estimates as heavily (sFigures 55-57). The fact that global signal correction so dramatically decreased average estimated amygdala—mPFC BSC may indicate that, like with resting-state fMRI analyses, positive functional connectivity values are due in part to motion and physiology-related confounds (Gratton et al., 2020; Power et al., 2019). Supporting this, BSC estimates were correlated with mean framewise displacement across scans for the fear > baseline and neutral > baseline contrasts only when a global signal correction was not applied (see sFigure 62). In addition, while test-retest reliability for all BSC measures was poor, BSC estimates from pipelines including a global signal correction step mostly demonstrated somewhat higher ICC (Figure 6). While these results are consistent with prior work indicating that correcting for the global signal can mitigate artifacts (Ciric et al., 2017; Satterthwaite et al., 2012), other work indicates that such corrections also remove meaningful biological signals (Belloy et al., 2018; Glasser et al., 2018; Yousefi et al., 2018).

### Amygdala–mPFC circuitry and separation anxiety

We did not find associations between any task-related amygdala–mPFC measures (reactivity or functional connectivity) and separation anxiety behaviors (see Table 1 Aim 3; Figure 6). This finding stands in contrast to associations between amygdala–mPFC connectivity and anxiety identified in previous developmental work (Gee et al., 2013; Jalbrzikowski et al., 2017; Kujawa et al., 2016; Qin et al., 2014). However, given that analyses of brain-behavior associations may require imaging cohorts much larger than the current sample (especially considering the low reliability of the measures used; Grady et al., 2020; Marek et al., 2020), the absence of relationships here may not be strong evidence against the existence of potential associations between amygdala–mPFC circuitry and developing anxiety-related behaviors.

### Advantages and pitfalls of the multiverse approach

Our findings contribute to a body of work demonstrating that preprocessing and modeling choices can meaningfully influence results (Botvinik-Nezer et al., 2020). Indeed, most studies involving many analytical decision points could benefit from multiverse analyses. Such specification curves can help to examine the stability of findings in both exploratory and confirmatory research (Flournoy et al., 2020). Particularly when methodological ‘gold standards’ have not been determined, specification curves may be informative for examining the impacts of potential analysis decisions (Bridgeford et al., 2020; Dafflon et al., 2020). Further, wider use of specification curves might help to resolve discrepancies between study findings stemming from different analysis pipelines.

While specification curve analyses may benefit much future research, we also note that multiverses are only as comprehensive as the included specifications (Steegen et al., 2016), and such analyses alone do not solve problems related to unmodeled confounds, design flaws, inadequate statistical power, circular analyses, or non-representative sampling. Further, unless all specifications are decided *a priori*, analyses are vulnerable to problems of analytic flexibility (Gelman & Loken, 2014), and inclusion of less justified specifications can bias results (Del Giudice & Gangestad, 2021). Because specification curves can include hundreds or thousands of individual analyses, rigorous evaluation of individual models can be difficult. To this end, we created interactive visualizations for visual exploration of individual analysis specifications.

Computational resources are a relevant concern when conducting multiverse analyses as well. In the current study, preprocessing (registration in particular) was the most computationally intensive step, taking an estimated 4 hours of compute time per scan per pipeline using 4 cores on a Linux-based institutional research computing cluster. However, specification curve analyses themselves were relatively less intensive, with all group-level models of amygdala reactivity completing in a total of 48 hours using 4 cores on a laboratory Linux-based server. Specification curves using maximum likelihood models (lme4 in R; <u>Bates & Bolker, 2011)</u> were even more efficient, with thousands of models running within minutes using a 2019 MacBook Pro (2.8 GHz Intel Core i7).

### Limitations

The present study is subject to several limitations that may be addressed in future investigations. Perhaps most crucially, our conclusions (along with those of many developmental fMRI studies) are limited by the poor test-retest reliability of the fMRI data. Because amygdala—mPFC measures showed low reliability across study visits, the statistical power of our analyses of age-related changes is likely low (Elliott et al., 2020; Zuo et al., 2019). Low-powered studies can yield increased rates of both false positive and false negative results (as well as errors of the sign and magnitude of estimates; Button et al., 2013; Gelman & Carlin, 2014); therefore we caution against interpretation of our developmental findings (and brain-behavior associations) beyond the cohort studied in the present investigation. In particular, the low statistical power of our rapid event-related task design may be a major contributor to the low test-retest reliability and variance in outcomes across analysis specifications. That being said, achieving high-powered studies presents a challenge for studying populations that cannot tolerate lengthy fMRI sessions. Both findings that were more robust to analytical decisions (amygdala reactivity) and findings that were less so (amygdala— mPFC connectivity, associations with separation anxiety) may be most valuable in meta-analytic contexts where greater aggregate statistical power can be achieved. In particular, future work on amygdala—mPFC development will benefit from optimization of measures both on robustness to analytic variability (Li et al., 2021) and reliability (Kragel et al., 2021).

Present findings are also limited by the number of participants studied (Bossier et al., 2020; Marek et al., 2020), the number of longitudinal study sessions per participant (King et al., 2018), and the duration of the task (Nee, 2019). Work with larger sample sizes, more study sessions per participant, and more task data collected per session will be necessary for charting functional amygdala–mPFC development and examining heterogeneity across individuals (although collecting task-based fMRI will continue to be challenging for studies including younger children). The generalizability of the current findings may also be limited by the fact that this cohort was skewed towards high incomes and not racially or ethnically representative of the Los Angeles or United States population.

Findings are also somewhat limited by the fact that the present study is not wholly confirmatory, despite preregistration. Because our multiverse analysis approaches expanded significantly beyond the methods we preregistered, most of the present analyses, while hypothesis-driven, must be considered exploratory (Flournoy et al., 2020). The fact that some specifications used data included in previous similar analyses of the same cohort(Gee et al., 2013) also limits the confirmatory power of the present study (Kriegeskorte et al., 2009). This may be especially true because longitudinal models could not identify within-person change as distinct from between-participant differences (see Figure 2D), indicating that our age-related change estimates may be influenced by cross-sectional information similar to that investigated by Gee et al. (Gee et al., 2013).

Though the current study aimed to estimate longitudinal age-related changes in amygdala–mPFC functional circuitry evoked by fear and neutral faces, the current findings may not be specific to these stimuli (Hariri et al., 2002). Because our task did not include non-face foils or probe specific emotion-related processes, results may be driven by attention, learning, or visual processing, rather than affective or face processing. In particular, because participants were instructed to press a button for neutral faces and withhold a button press for fear faces, observed amygdala—mPFC responses may in part reflect response inhibition <u>(for fear faces; Menon et al., 2001)</u> and target detection processes <u>(for neutral faces; Jonkman et al., 2</u>003). Findings for the fear faces > baseline and neutral > baseline contrasts also may not be valence-specific in the absence of a different emotional face as part of the contrast. Further, because all faces were adult White women, the current results may not generalize to faces more broadly (Richeson et al., 2008; Telzer et al., 2012). Additionally, because face stimuli were the same across study visits, exposure effects across sessions may confound longitudinal findings (although exposure effects may be possible any time a task is repeated, even if stimuli are unique), particularly age-related decreases in amygdala responses (Telzer et al., 2018). While within-session amygdala habituation effects have been shown across several paradigms (Geissberger et al., 2020; Hare et al., 2008; Hein et al., 2018), between-session habituation effects are unlikely beyond 2-3 weeks (Geissberger et al., 2020; Johnstone et al., 2005; Plichta et al., 2014; Spohrs et al., 2018).

Finally, our findings on age-related change in amygdala and mPFC function may be biased or confounded by age-related differences in head motion (Ciric et al., 2017), anatomical image quality and alignment (Gilmore et al., 2020; Rorden et al., 2012), signal dropout, and physiological artifacts (Boubela et al., 2015; Fair et al., 2020; Gratton et al., 2020). While our multiverse analyses included preprocessing and group-level modeling specifications designed to minimize some of such potential issues, future work is still needed to optimize discrimination of developmental changes of interest from such potential confounds.

Despite these limitations, the present study concords with prior investigations in demonstrating the value of multiverse approaches to quantify sensitivity to researcher decisions. The results highlight key analytic considerations for future studies of age-related changes in amygdala—mPFC function, as well as for studies of human brain development more broadly.

## CRediT Author Statement

**Paul Alexander Bloom:** Conceptualization, Methodology, Formal Analysis, Writing - Original Draft, Visualization **Michelle VanTieghem:** Methodology, Writing – Review & Editing **Laurel Gabard-Durnam:** Investigation, Methodology, Writing – Review & Editing, **Dylan Gee:** Investigation, Methodology, Writing – Review & Editing **Jessica Flannery:** Investigation, Writing – Review & Editing **Christina Caldera:** Investigation, Writing – Review & Editing **Bonnie Goff:** Investigation, Writing – Review & Editing **Eva Telzer:** Investigation, Writing – Review & Editing **Kathryn L. Humphreys:** Investigation, Writing – Review & Editing **Dominic Fareri:** Investigation, Writing – Review & Editing **Mor Shapiro:** Investigation, Writing – Review & Editing **Sameah Algharazi:** Validation, Writing – Review & Editing **Niall Bolger:** Methodology, Formal Analysis, Writing – Review & Editing **Mariam Aly:** Methodology, Formal Analysis, Supervision, Writing – Review & Editing **Nim Tottenham:** Conceptualization, Methodology, Investigation, Resources, Data Curation, Writing - Original Draft, Supervision, Funding Acquisition

## Code Availability

Code for preprocessing, analysis, and data visualizations for this manuscript is available at https://github.com/pab2163/amygdala_mpfc_multiverse. While unfortunately this code cannot be run as written without data, we have attempted to document analysis steps clearly. In addition, we provide publicly available simulated data structured similarly to the study data on amygdala reactivity, such that interested readers can view multiverse analysis walkthroughs (https://pab2163.github.io/amygdala_mpfc_multiverse) and experiment with analysis code. Additional materials, including MNI space masks and preregistration documentation, are available at https://osf.io/hvdmx/

## Notes

**Funding** This work was supported by funding from the National Institutes of Mental Health (2R01MH091864) and the Dana Foundation for Nim Tottenham, and a National Science Foundation Graduate Research Fellowship (DGE 1644869) for Paul Alexander Bloom. The authors declare no competing interests.

https://pab2163.github.io/amygdala_mpfc_multiverse/

### Competing Interest Statement

The authors have declared no competing interest.

### Summary of Updates

Figure 7 added, manuscript revised, supplement updated.

