## Supplement for "Age-related change in task-evoked amygdala—prefrontal circuitry: a multiverse approach with an accelerated longitudinal cohort aged 4-22 years"

Table of Contents

**PREVIOUS STUDIES OF AGE-RELATED CHANGE IN AMYGDALA—MPFC FUNCTION** 5
**sTable 1:** Previous studies of age-related change in amygdala—mPFC function 8

**SUPPLEMENTAL METHODS** 9

**Participant demographics** 9
**sTable 2:** Race, gender, and ethnicity distributions for study participants 10
**sTable 3:** Parent free responses for their children's racial identities beyond available categories 10
**sFigure 1:** Distribution of annual household income for participating families. 11
**sFigure 2:** Parent-reported CBCL scores 11

**Analyses of task behavior** 11

**Reactivity Analyses** 13
*C-PAC preprocessing pipeline* 13
*Amygdala reactivity: multiverse details (Table 2, Aim 1)* 14
*Longitudinal Amygdala Model Syntax* 17
*Parametrization of within-participant change versus between-participant differences* 18
*Within-person similarity of reactivity voxel-wise statistical maps across multiverse forks* 19
*Between-scan associations between amygdala reactivity estimates across specifications* 20
*Dependence of amygdala reactivity findings on previous work* 20

**Within-scan changes in amygdala reactivity across trials** 21
*Changes in amygdala reactivity across trials: multiverse details (Table 2, Aim 2)* 21
*Longitudinal model syntax for models of within-scan change in amygdala reactivity* 22

**Amygdala—mPFC functional connectivity analyses** 23
*Amygdala—mPFC gPPI analyses: multiverse details (Table 2, Aim 3)* 23
*Amygdala—mPFC BSC analyses: multiverse details (Table 2, Aim 3)* 25
*Longitudinal models for functional connectivity* 26
*Rank-order associations between gPPI & BSC amygdala—mPFC FC* 26
*Effects of up-sampling/lasso regularization parameters during deconvolution on gPPI regressors* 26
*Within-person similarity of voxel-wise statistical maps for amygdala FC from gPPI pipelines* 27
*Within-person similarity between gPPI & BSC amygdala FC* 27

**Associations between amygdala—mPFC circuitry & separation anxiety:** 28
*Amygdala—mPFC circuitry & separation anxiety: multiverse details (Table 2, Aim 5)* 28
*Longitudinal model syntax for amygdala—mPFC circuitry & separation anxiety* 29

**Estimating impacts of specific forked decision points on age-related change estimates** 29

**BEHAVIOR: SUPPLEMENTAL RESULTS** 31

***Behavioral Task Performance*** 31
**sFigure 3:** d' as a function of age. 31
**sFigure 4:** Task performance metrics as a function of age 32

***Head Motion*** **33**
**sFigure 5:** Mean framewise displacement as a function of age (years) 33

### AMYGDALA REACTIVITY: SUPPLEMENTAL RESULTS 33

***Group Mean Amygdala Reactivity*** **34**
**sFigure 6:** Specification curve for mean fear > baseline amygdala reactivity 35
**sFigure 7:** Specification curve for mean neutral > baseline amygdala reactivity 36
**sFigure 8:** Specification curve for mean fear > neutral amygdala reactivity 37

***Multiverse analyses of age-related changes in amygdala reactivity*** **38**
**sFigure 9:** Specification curve for age-related change in neutral > baseline amygdala reactivity 39
**sFigure 10:** Specification curve for age-related change in fear > neutral amygdala reactivity 39

***Impacts of pipeline choices on age-related change estimates for amygdala reactivity*** **40**
**sFigure 11:** Fork impacts on age-related change for fear > baseline amygdala reactivity 40
**sFigure 12:** Fork impacts on age-related change for neutral > baseline amygdala reactivity 41
**sFigure 13:** Fork impacts on age-related change for fear > neutral amygdala reactivity 42

***Nonlinear Changes in Amygdala Reactivity*** **42**
**sFigure 14:** Spec. curve for quadratic age-related change in fear > baseline amygdala reactivity 44
**sFigure 15:** Spec. curve for quadratic age-related change in neutral > baseline amygdala reactivity 45
**sFigure 16:** Spec. curve for quadratic age-related change in fear > neutral amygdala reactivity 46
**sFigure 17:** Predictions of age-related change across linear and quadratic model specifications. 47
**sFigure 18**: Inverse age models for amygdala reactivity 48

***Between-participant age associations versus within-participant age-related change*** **48**
**sFigure 19:** Differences between within-participant and between-participant terms for age-related associations with amygdala reactivity 49

***Within-person similarity of voxel-wise amygdala reactivity statistical maps across forks*** **49**
**sFigure 20.** Voxelwise image similarity for amygdala reactivity contrasts between the preregistered FSL pipeline and C-PAC pipelines. 51
**sFigure 21:** Voxelwise similarity for amygdala reactivity contrasts between all C-PAC pipelines. 52

***Between-scan correlations of amygdala reactivity estimates across specifications*** **52**
**sFigure 22:** Between scan correlations for amygdala reactivity pipelines for fear > baseline. 53
**sFigure 23:** Between-scan correlations for amygdala reactivity pipelines for neutral > baseline. 54
**sFigure 24:** Between scan correlations for amygdala reactivity pipelines for fear > neutral. 55

***Dependence of amygdala reactivity age-related change findings on previous work*** **55**
**sFigure 25:** Permutation tests for age-related change excluding previously studied scans 57

***Head Motion & Amygdala Reactivity*** **57**
**sFigure 26:** Correlations between head motion and amygdala reactivity for each contrast 58
**sFigure 27:** Estimated age-related change as a function of mean FD exclusion threshold 59

### *MPFC REACTIVITY: SUPPLEMENTAL RESULTS* 59
**sFigure 28:** Group average mPFC reactivity and age-related change for fear faces > baseline 60
**sFigure 29:** Associations between amygdala & mPFC reactivity 61

### WITHIN-SCAN CHANGES IN AMYGDALA REACTIVITY: SUPPLEMENTAL RESULTS 61

***Group average within-scan changes in amygdala reactivity*** **61**
**sFigure 30:** Group average slopes in amygdala reactivity across trials 62
**sFigure 31:** Group average differences in amygdala reactivity across first > second half of trials 63

***Age-related differences in within-scan amygdala reactivity change*** **63**
**sFigure 32:** Spec. curve for differences across task half in amygdala reactivity age-related change 64
**sFigure 33:** Multiverse single-trial model predictions as a function of trial and age 65
**sFigure 34:** Spec. curve of age*trial amygdala reactivity interactions from single-trial models 66

### GPPI FUNCTIONAL CONNECTIVITY RESULTS 66

***Impacts of a deconvolution step on gPPI regressors and estimates*** **66**
**sFigure 35:** Example gPPI and amygdala seed regressors for one scan 68
**sFigure 36:** Correlations between gPPI regressors, and between gPPI regressors and the seed 69
**sFigure 37:** Voxelwise image similarities of gPPI estimates with vs. without deconvolution 70

***Group mean amydala-mPFC gPPI estimates*** **70**
**sFigure 38:** Group average amygdala—mPFC gPPI estimates for each contrast and ROI 71

***Multiverse analyses of age-related change in amygdala—mPFC gPPI*** **71**
**sFigure 39:** Spec. curve for age related change in amygdala—mPFC gPPI for neutral > baseline 72
**sFigure 40:** Spec. curve for age related change in amygdala—mPFC gPPI for fear > neutral 73

***Impacts of analysis choices on age-related change estimates for amygdala—mPFC gPPI*** **73**
**sFigure 41.** Fork impacts on age-related change for fear > baseline amygdala—mPFC gPPI 74
**sFigure 42:** Fork impacts on age-related change for neutral > baseline amygdala—mPFC gPPI 75
**sFigure 43:** Fork impacts on age-related change for fear > neutral amygdala—mPFC gPPI 75

***Impacts of regressor centering on age-related change in amygdala—mPFC gPPI*** **75**
**sFigure 44:** Impacts of centering the task regressor in gPPI models with deconvolution 76

***Non-linear age-related changes in amygdala—mPFC gPPI*** **76**
**sFigure 45:** Spec. curve for quadratic age-related changes in fear > baseline amygdala—mPFC gPPI 78
**sFigure 46:** Spec. curve for quadratic age-related changes in neutral > baseline amygdala—mPFC gPPI 78
**sFigure 47:** Spec. curve for quadratic age-related changes in fear > neutral amygdala—mPFC gPPI 79
**sFigure 48:** Inverse age models for amygdala—mPFC gPPI 80

***Correlations between head motion and amygdala—mPFC gPPI estimates*** **80**
**sFigure 49:** Correlations between mean FD & gPPI estimates across scans 81

***Task-independent amygdala—mPFC connectivity estimates from gPPI models*** **81**
**sFigure 50:** Group average task-independent amygdala—mPFC connectivity 82
**sFigure 51:** Age-related change in task-independent amygdala—mPFC connectivity 82

### BSC FUNCTIONAL CONNECTIVITY RESULTS 83

***Group mean amygdala—mPFC BSC*** **83**
**sFigure 52:** Group mean amygdala—mPFC BSC across contrasts, mPFC ROIs, and pipelines 83

***Multiverse analyses of age-related change in amygdala—mPFC BSC*** **83**
**sFigure 53:** Spec. curve for age-related change in neutral > baseline BSC 84
**sFigure 54:** Spec. curve for age-related change in fear > neutral BSC 85

***Impacts of analysis choices on age-related change estimates for amygdala—mPFC BSC*** **85**
**sFigure 55:** Fork impacts on age-related change for fear > baseline amygdala—mPFC BSC 86
**sFigure 56:** Fork impacts on age-related change for neutral > baseline amygdala—mPFC BSC 87
**sFigure 57.** Fork impacts on age-related change for fear > neutral amygdala—mPFC BSC 87

***Nonlinear age-related changes in amygdala—mPFC BSC*** **87**
**sFigure 58:** Spec. curve for quadratic age-related changes in fear > baseline amygdala—mPFC BSC 88
**sFigure 59:** Spec. curve for quadratic age-related changes in neutral>baseline amygdala—mPFC BSC 89
**sFigure 60:** Spec. curve for quadratic age-related changes in fear > neutral amygdala—mPFC BSC 90
**sFigure 61:** Inverse age models for amygdala—mPFC BSC 91

***Correlations between head motion & amygdala—mPFC BSC estimates*** **91**
**sFigure 62:** Correlations between mean FD & BSC estimates across scans 92

***Within-scan similarity between gPPI & BSC amygdala FC*** **92**
**sFigure 63.** Similarity of amygdala FC with the rest of the brain between gPPI & BSC 93

***Between-scan correlations between gPPI & BSC estimates*** **93**
**sFigure 64:** Between-scan correlations between fear > baseline gPPI & BSC estimates 95
**sFigure 65:** Between-scan correlations between neutral > baseline gPPI & BSC estimates 96
**sFigure 66:** Between-scan correlations between fear > neutral gPPI & BSC estimates 96

**Associations with generalized anxiety and social anxiety behaviors** **96**
**sFigure 67:** Associations between amygdala—mPFC measures and anxiety-related behaviors 97

### Previous Studies of Age-Related Change in Amygdala—mPFC Function

| Authors | Year | N | Design | Ages | Task | Faces | MRI Design | Contrast | Reactivity | Amyg—mPFC Connectivity | Connectivity Method |
|---|---|---|---|---|---|---|---|---|---|---|---|
| **Baird et al.** | 1999 | 12 | Cross-sectional | 12 to 17 | Emotion labeling with fear faces | Ekman & Friesen, 1976 | Block | fear > nonsense grayscale figures | No age-related changes | NA | NA |
| **Killgore et al.** | 2001 | 19 | Cross-sectional | 9 to 17 | Emotion labeling with fear faces | Ekman & Friesen, 1976 | Block | fear > baseline | Age-related decrease in left amyg, but not right amyg, only in females | NA | NA |
| **Thomas et al.** | 2001 | 18 | Cross-sectional | Youth (mean = 11, sd = 2.4), and male adults (mean = 24, sd = 6.6) | Passive viewing with fear & neutral faces | Ekman & Friesen, 1976 | Block | fear > neutral | Adults show higher fear > neutral reactivity in left amyg than children | NA | NA |
| **Pine et al.** | 2001 | 20 | Cross-sectional | Youth (age 12-16) and adults (age 25-38) | Masking paradigm with happy and fear faces | Ekman & Friesen, 1976 | Block | masked fear > fixation, masked happy > fixation | No group differences in amygdala reactivity for any contrast | NA | NA |
| **Monk et al.** | 2003 | 34 | Cross-sectional | Youth (9-17) and adults (25-36) | 1) passive viewing with fear, angry, happy, neutral 2) emotional rating of faces 3) subjective rating of nose width | NimStim (2009); Ekman & Friesen, 1976; Gur, 2001 | Event-related | A) rating of fear for fear faces > nose width for fear faces, B) nose width for fear faces > nose width for neutral faces, C) passive viewing for fear faces > passive viewing for neutral faces | No age-related amygdala differences for contrasts A/B. For C, adolescents show higher reactivity than adults in right amyg, but no age-related change within adolescent group | NA | NA |
| **McClure et al.** | 2004 | 34 | Cross-sectional | Youth (9-17) and adults (25-36) | Threat rating for fear, angry, happy, neutral faces | NimStim (2009); Ekman & Friesen, 1976; Gur, 2001 | Event-related | Angry faces > all other faces, each emotion separately > baseline | Only in females, adults showed greater right amygdala reactivity to angry > neutral and angry > fear faces | NA | NA |
| **Yurgelun-Todd & Killgore** | 2006 | 16 | Cross-sectional | 8 to 15 | Passive viewing of fearful and happy faces | Ekman & Friesen, 1976 | Block | fear > baseline, happy > baseline | No age-related changes in amygdala reactivity for either contrast | NA | NA |
| **Killgore & Yurgelun-Todd** | 2007 | 22 | Cross-sectional | Adolescents (age 9-17) and adults (mean= 23.7, sd=2.1) | Masking paradigm with sad and happy faces | Erwin et al., 1992 | Block | masked sad > baseline, masked happy > baseline, masked happy > masked sad | Adolescents show greater right amygdala activation for masked sad > baseline than adults. No differences for other contrasts | NA | NA |

| Authors | Year | N | Design | Ages | Task | Faces | MRI Design | Contrast | Reactivity | Amyg—mPFC Connectivity | Connectivity Method |
|---|---|---|---|---|---|---|---|---|---|---|---|
| **Guyer et al.** | 2008 | 61 | Cross-sectional | Adolescents (9-17) and adults (21-40) | Passive viewing with fear, angry, happy, neutral faces | NimStim (2009); Ekman & Friesen, 1976; Gur, 2000 | Event-related | fear > neutral, fear > fixation, neutral > fixation | Adolescents show greater amygdala activation than adults for fear > neutral and fear > fixation. No difference for the neutral > fixation contrast. No age-related change within adolescent group. | No differences in seed-based amyg—mPFC functional connectivity between adolescents and adults | Seed-based correlation |
| **Hare et al.** | 2008 | 60 | Cross-sectional | Children (7-12), Adolescents (13-18), Adults (19-32) | Go/no-go with fear, happy, calm faces. All combinations of emotions used as targets | NimStim (2009) | Event-related | fear > baseline, fear > calm | Adolescents show greatest amygdala activation to fear > baseline compared to children and adults. No differences in amygdala activation to fear > baseline between children and adults | NA | NA |
| **Passarotti et al.** | 2009 | 20 | Cross-sectional | Adolescents (mean age = 14), adults (mean age 30) | Age judgement, affect judgement with angry and happy faces | Gur, 2002 | Event-related | angry + happy > fixation | Adolescents show greater right amyg activation than adults in the incidental condition > fixation under a liberal contiguity threshold, but not a strict one. No group differences in amygdala during directed condition | NA | NA |
| **Somerville et al.** | 2010 | 62 | Cross-sectional | 6 to 29 | Go/no-go with fear, happy, calm faces. All combinations of emotions used as targets | NimStim (2009) | Event-related | happy > baseline, calm > baseline | No age-related changes in amygdala from whole-brain analysis | NA | NA |
| **Pfeifer et al** | 2011 | 38 (76 scans) | Longitudinal (2 waves) | 10 to 13 | Passive viewing with fear, angry, happy, sad, neutral faces | NimStim (2009) | Event-related | all faces individually > fixation, all faces together > fixation, all emotional faces individually > neutral faces | No age-related change in right or left amygdala activity in all faces averaged together > fixation. Increase in activity in the right amygdala in response to sad > neutral faces | NA | NA |
| **Forbes et al.** | 2011 | 76 | Cross-sectional | 11 to 13 | Matching with fear, angry, neutral faces, and shapes | NimStim (2009) | Block | fear > shapes, angry > shapes, neutral > shapes | No differences for fear > shapes. Pre/early-pubertal adolescents show greater amyg reactivity to neutral > shapes than mid/late pubertal adolescents | NA | NA |
| **Todd et al.** | 2011 | 45 | Cross-sectional | 31 children (age 3.5-8.5), 14 young adults (age 18-33) | Passive viewing of angry and happy mother/stranger faces, phase-scrambled images | Angry & happy face images of mothers of participants | Block | all faces > scrambled faces, angry > scrambled faces, happy > scrambled faces | Age-related increase in bilateral amyg activation to faces > scrambled faces, and angry faces > scrambled faces | NA | NA |
| **Perlman & Pelphrey** | 2011 | 20 | Cross-sectional | 5 to 11 | Go/no-go with fear faces interspersed. Also, block structure with winning, losing, and recovery of points | NimStim (2009) | Event-related | Effective connectivity calculated during block 3 of the task | NA | Effective connectivity during block 3 between left amygdala and inferior frontal gyrus/ACC increased with age | Granger causality |
| **Telzer et al.** | 2012 | 32 | Cross-sectional | 4 to 16.5 | Matching with angry, happy, | NimStim (2009) | Block | African-American faces > baseline, European- | Right amygdala response to African-American faces > baseline increases with age, no | NA | NA |

| Authors | Year | N | Design | Ages | Task | Faces | MRI Design | Contrast | Reactivity | Amyg—mPFC Connectivity | Connectivity Method |
|---|---|---|---|---|---|---|---|---|---|---|---|
| | | | | | neutral faces, and shapes | | | American faces > baseline | age-related change in right amygdala response to European-American faces | | |
| **Gee et al.** | 2013 | 45 | Cross-sectional | 4 to 22 | Go/no-go with fear, happy, and neutral faces. Withhold press for emotional faces | NimStim (2009) | Event-related | fear > baseline | Age-related decreases in right amygdala | Positive early in development between right amyg—mPFC, turning negative in adolescence | gPPI with AFNI deconvolution |
| **Swartz et al.** | 2014 | 39 | Cross-sectional | 9 to 19 | Gender identification with fear, happy, sad, neutral faces | NimStim (2009) | Event-related | all emotions individually > baseline | Age-related decreases in left amygdala to fear > baseline. Age-related decreases in left amygdala to each other emotion individually > baseline | No age-related change in right or left amygdala with mPFC for all faces > baseline contrast | gPPI with SPM8, deconvolution not mentioned |
| **Dreyfuss et al.** | 2014 | 80 | Cross-sectional | 6 to 27 | Go/no-go with fear, happy, calm faces. All combinations of emotions used as targets | NimStim (2009) | Event-related | fear > calm | No age-related changes in amygdala from whole-brain analysis | NA | NA |
| **Telzer et al.** | 2015 | 52 | Cross-sectional | 4 to 18 | Matching with angry, happy, neutral, faces, and shapes | NimStim (2009) | Block | same sex > shapes, opposite sex > shapes | Bilateral amygdala responses to opposite-sex faces > shapes decrease with age, bilateral amygdala response to same-sex faces > shapes increase with age | NA | NA |
| **Joseph et al.** | 2015 | 42 | Cross-sectional | 5 to 18 | Passive viewing of faces (73% happy, the rest neutral), objects, and textures | Custom unfamiliar high-school yearbook faces | Block | face > texture | Age-related increase in right and left amyg reactivity to faces > textures contrast | NA | NA |
| **Wu et al.** | 2016 | 61 | Cross-sectional | 7 to 25 | Matching with fear, angry, happy, neutral faces, and shapes. | Hariri et al., 2002 | Block | fear > shapes, angry > shapes, happy > shapes | No age-related change in amygdala found for any contrast, or all together | For each emotion > shapes, positive early in development between left amyg-ACC, turning to negative around age 15. Same with right amyg | PPI done in AFNI |
| **Kujawa et al.** | 2016 | 61 | Cross-sectional | 7 to 25 | Matching with fear, angry, happy, neutral faces, and shapes. | Hariri et al., 2002 | Block | fear > shapes, angry > shapes, happy > shapes | No age-related change effects in either hemisphere for either emotion > shapes | Positive early in development between both hemispheres/ACC, turning negative in around age 15 | PPI done in SPM with deconvolution |
| **Heller et al.** | 2016 | 155 | Cross-sectional | 5 to 32 | Go/no-go with fear, happy, calm faces. All combinations of emotions used as targets | NimStim (2009) | Event-related | happy > baseline, calm > baseline | NA | No age-related changes reported in whole-brain analysis | Beta series correlation analysis done with amygdala seed to whole brain |
| **Vijayakumar et al.** | 2019 | 82 | Longitudinal (3 waves) | 9 to 18 | Passive viewing with fear, angry, happy, sad, neutral faces | NimStim (2009) | Event-related | all emotions together > baseline, all emotions individually > baseline | None for fear > baseline. Age-related increase for sad > baseline, and sad > neutral | NA | NA |
| **Zhang et al.** | 2019 | 759 | Cross-Sectional | 8 to 23 | Emotion labeling with fear, angry, happy, sad faces | Gur, 2002 | Rapid Event-related | all emotions individually > baseline | Depending on ROI definition and model, either no age-related change or age-related increases for fear > baseline, happy > baseline, sad > baseline | Depending on ROI definition and model, either no age-related change in amyg-PFC connectivity, or age-related increases for fear, happy, angry | gPPI & BSC |

| Authors | Year | N | Design | Ages | Task | Faces | MRI Design | Contrast | Reactivity | Amyg—mPFC Connectivity | Connectivity Method |
|---|---|---|---|---|---|---|---|---|---|---|---|
| **Xu et al.** | 2021 | 321 | Cross-Sectional | 243 (7-12), 78 young adults (19-25) | Emotion matching with fear & angry faces | Wang & Luo (2005) | Block | negative (fear/angry) faces > shapes | Age*sex interaction -- age-related decreases in females in BLA and CMA, increases (though not always significant) in males | age*sex interaction -- age-related increases in males, decreases in females | gPPI with deconvolution using SPM |

**sTable 1:** Previous studies of age-related change in amygdala—mPFC function
We summarize cohort, task design, contrast, connectivity method (when available), and amygdala—mPFC result information from prior work with fMRI analyses of age-related differences in amygdala—mPFC responses to faces (Baird et al., 1999; Dreyfuss et al., 2014; Ekman & Friesen, 1976; Erwin et al., 1992; Forbes et al., 2011; Gee et al., 2013; Gur et al., 2002; Guyer et al., 2008; Hare et al., 2008; Hariri et al., 2002; Heller et al., 2016; Joseph et al., 2015; Killgore et al., 2001; Killgore & Yurgelun-Todd, 2007; Kujawa et al., 2016; McClure et al., 2004; Monk et al., 2003; Passarotti et al., 2009; Perlman & Pelphrey, 2011; Pfeifer et al., 2011; Pine et al., 2001; Somerville et al., 2010; Swartz et al., 2014; Telzer et al., 2012, 2015; Thomas et al., 2001; Todd et al., 2011; Tottenham et al., 2009; Vijayakumar et al., 2019; Wang & Luo, 2005; Wu et al., 2016; Xu et al., 2021; Yurgelun-Todd & Killgore, 2006; Zhang et al., 2019).

### Supplemental Methods

Participant demographics

Parents reported their child's gender, race, and whether their child was *"Hispanic or Latino"* via questionnaire items (except for 10 participants ages 18-22 who self-reported). Parents indicated separately whether their child's race fell in differeint categories (*African-American/Black, American Indian/Alaska Native, Asian-American, European-American/Caucasian, Native Hawaiian or Other Pacific Islander;* see sTable 2). Some parents indicated multiple race categories (therefore total proportions of racial groups sum to greater than 1). Parents also had the option to indicate any other racial categories their children belonged to through an open-response item (see sTable 3). Parents also reported annual household income within discrete bins (see sFigure 1). Income data was missing for 13 families.

| | | *N* | *Proportion* |
|---|---|---|---|
| Gender | Female | 55 | 0.56 |
| | Male | 43 | 0.44 |
| Race | European-American/Caucasian | 56 | 0.57 |
| | African-American/Black | 28 | 0.29 |
| | Asian-American | 24 | 0.24 |
| | American Indian/Alaska Native | 3 | 0.03 |
| | Native Hawaiian or Other Pacific Islander | 1 | 0.01 |
| Hispanic or Latino ethnicity | Hispanic or Latino | 12 | 0.12 |

| | | |
|---|---|---|
| Not Hispanic or Latino | 84 | 0.86 |
| Missing Data | 2 | 0.02 |

**sTable 2:** Race, gender, and ethnicity distributions for study participants

| *Response* | *N* |
|---|---|
| Hispanic | 3 |
| Arab | 2 |
| Filipino | 1 |
| Iranian American | 1 |
| Mexican | 1 |
| Spanish | 1 |

**sTable 3:** Parent free responses for their children’s racial identities beyond available categories

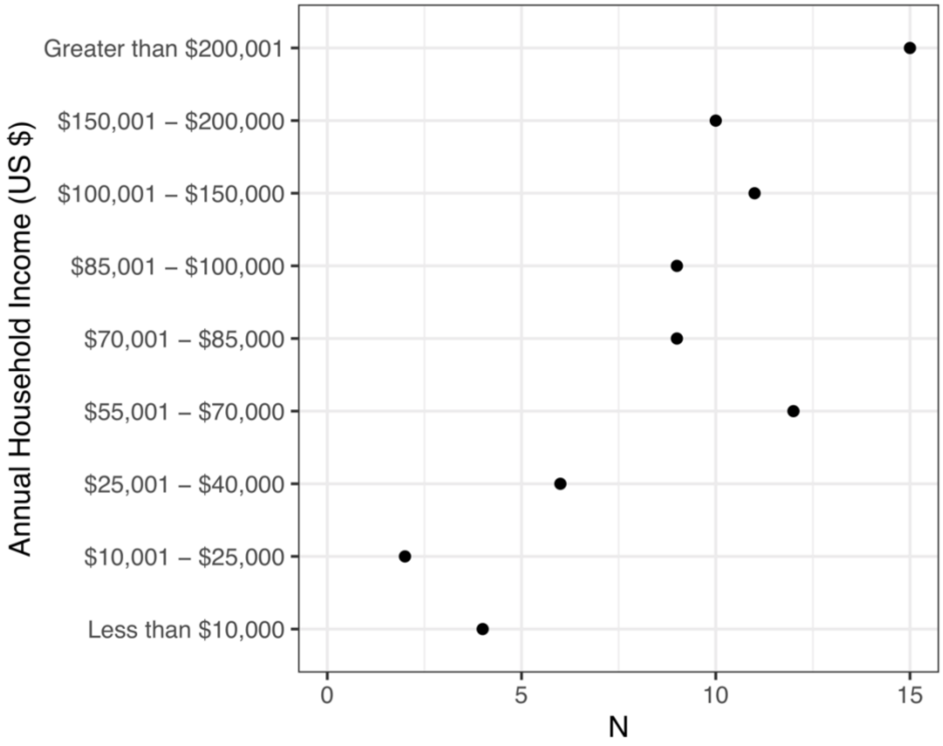

**sFigure 1:** Distribution of annual household income for participating families.

*CBCL Scores:*

Parents reported on child emotional and behavioral symptoms via the Child Behavior Checklist (CBCL; Achenbach, 1991) for age 1.5-5 and 4-18 years. Age and gender-normed scores for participants' first study timepoint indicated that 4 participants met clinical threshold for internalizing problems, while none met clinical threshold for externalizing or total problems.

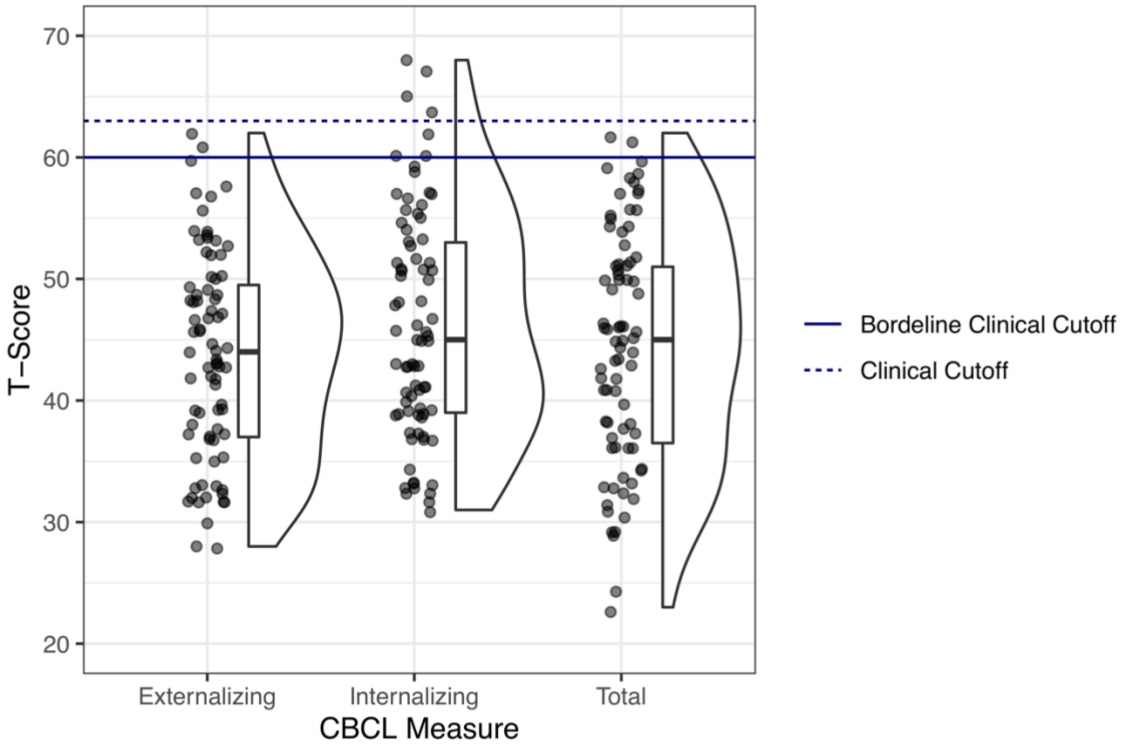


**sFigure 2:** Parent-reported CBCL scores
Boxplots show medians and IQRs for CBCL T-scores for each scale, and points are individual participants. No participants met clinical cutoff for externalizing or total problems at timepoint 1, while 4 participants met the clinical cutoff for internalizing problems.

Analyses of task behavior

To characterize age-related changes in task performance, we modeled behavior in several ways. First, we calculated the d' performance metric for each scan using a correction for extreme proportions (Hautus, 1995) where button presses for neutral faces were considered ‘hits’, withheld presses for neutral faces ‘misses’, withheld presses for fear faces ‘correct rejections’, and presses for fear faces ‘false alarms’. We constructed multilevel models to look at linear only, linear + quadratic, and linear + quadratic + cubic age-related changes in d’ with the following R syntax:

```
# Linear:
brm(dprime ~ age + (age | participant))

# Linear + Quadratic:
brm(dprime ~ poly(age, 2, raw = TRUE) + (age | participant))

# Linear + Quadratic + Cubic:
brm(dprime ~ poly(age, 3, raw = TRUE) + (age | participant))

# Inverse age (1/age)
brm(dprime ~ age_inverse + (age_inverse | participant))
```

We fit all models using the brms package (Bürkner, 2017), and all models included varying linear slopes for age and intercepts across participants. In addition, to characterize more specific aspects of task behavior, we fit separate single-trial multilevel logistic regression models with overall accuracy (probability of a correct response on any trial), hit rate (on neutral face trials), and false alarm rate (on fear face trials) as the respective outcomes and included nested varying effects for task sessions within participants. For the accuracy model, ‘hits’ and ‘correct rejections’ were coded as 1 and ‘misses’ and ‘false alarms’ coded as 0.

```
# Accuracy:
brm(accuracy ~ age + (age |participant/session), family = bernoulli(link = 'logit'))

# False Alarms:
brm(false_alarms  ~ age + (age |participant/session), family = bernoulli(link = 'logit') )
```

```
# Hits:
brm(hits  ~ age + (age |participant/session), family = bernoulli(link = 'logit'))
```

To examine age-related changes in reaction times (for hits only), we used similar single-trial multilevel regression models with linear only, linear + quadratic, and linear + quadratic + cubic terms and nested varying effects for task sessions within participants.

```
# Linear:
brm(reaction_time ~ age + (age |participant / session))

# Linear + Quadratic:
brm(reaction_time ~ poly(age, 2, raw = TRUE) + (age |participant / session))

# Linear + Quadratic + Cubic:
brm(reaction_time ~ poly(age, 3, raw = TRUE)  + (age |participant / session))

# Inverse Age (age_inverse = 1/age)
brm(reaction_time ~ age_inverse + (age_inverse |participant / session))
```

### Reactivity Analyses

*C-PAC preprocessing pipeline*

In addition to the preregistered FSL preprocessing pipeline, we preprocessed BOLD images using C-PAC software ( v1.4.1; Craddock et al., 2013). For these pipelines, we mostly used software defaults. In these pipelines, ANTS (Tustison et al., 2013) was used to skull-strip MPRAGE images, and slice-time correction was applied (slices were acquired in interleaved order). BOLD spatial realignment and motion parameters were calculated with MCFLIRT as with the preregistered pipeline. Registration matrices were then calculated for functional images to be registered to high-resolution structural T1 images using FSL's FLIRT with boundary-based registration. Registration matrices for T1 images to standard MNI space were calculated using ANTS, and functional images were warped to MNI space before running GLMs.

*Amygdala reactivity: multiverse details (Table 2, Aim 1)*

As detailed in in the main manuscript, we constructed forking pipelines for analyses of age-related change in amygdala reactivity. Below we provide details into each decision point.

a. **GLM Software:** Within-participant first-level GLMs were conducted either using FEAT or AFNI 3dDeconvolve. Prewhitening as specified in software defaults was used in all GLMs to adjust for temporal autocorrelation.

b. **Hemodynamic Response Function:** Within the GLM, regressors for fear and neutral faces were convolved with either a canonical double-gamma or single-gamma hemodynamic response function.

c. **Nuisance Regressors:** Nuisance regressors were added to the first-level GLMs in all analysis pipelines. Pipelines included either 6 head motion regressors, 24 motion regressors (as preregistered; 3 for rotation, 3 for translation, their temporal derivatives, and the square roots of all the above; see Friston et al., 1996), or 18 motion regressors plus additional regressors for mean white matter and cerebrospinal fluid signal (C-PAC preprocessing only, 6 head motion regressors, their squares, and their backwards derivatives). In addition, to remove low-frequency artifacts, we either applied a high-pass filter (cutoff = .01Hz) to BOLD data before the GLM, or included a quadratic drift term in the model (AFNI GLMs only). In all pipelines, TRs with >.9mm framewise displacement (FD) were down-weighted to 0 in the GLM, effectively removing their influence on the model.

d. **First-Level GLM Estimates:** From each first-level GLM, we estimated contrasts for fear faces > baseline, neutral faces > baseline, and fear > neutral faces for each voxel using an event-related design. Although the fear > baseline contrast was of primary interest to the current study in following up work by Gee et al. (2013), we repeated analyses for the other contrasts as well. For each contrast, we either submitted beta estimates for each contrast (i.e. FSL COPEs), or t-statistics for group-level models. While the beta estimates represent the raw magnitude of estimated contrast effects for each scan, the t-statistics represent standardized effect sizes; i.e. these magnitudes scaled by the estimation uncertainty.

e. **Amygdala ROIs:** We conducted reactivity analyses with the amygdala defined in both native space and in standard MNI space. For native space analyses, participant-specific native space masks were defined using Freesurfer (v6.0; Fischl, 2012) for the bilateral amygdala, as well as left and right amygdala separately, and manually inspected for quality assurance by an experimenter. For analyses in standard space, we used an amygdala mask from the Harvard-Oxford Atlas (probability threshold = .5). In addition, to check whether effects were driven by signal dropout, we performed a median split of all voxels in the right and left amygdala, respectively, based on the grand mean BOLD intensity across all scans to create 'high signal' and 'low signal' amygdala sub-regions. All amygdala masks are available on OSF (https://osf.io/hvdmx/). For all amygdala ROI definitions, we calculated the mean reactivity across included voxels for each scan for bilateral, left, and right regions respectively for group-level analyses.

f. **Exclusion of previously analyzed scans:** 45 of the scans in this dataset were previously analyzed by Gee et al. (2013). Because these scans were used in a whole-brain mass univariate analysis as a discovery sample to identify regions changing with age in both reactivity and connectivity with the amygdala for the fear > baseline contrast, analyses in the current study including these scans will could be partially dependent (i.e. circular analysis) on the previous selection process (Kriegeskorte et al., 2009, 2010). Therefore, we included pipelines in the multiverse that excluded the 42 scans (3 scans originally included were excluded in the current study for high motion) previously analyzed. In addition, to disentangle whether any differences in results between such pipelines are due to nonindependence versus reduced sample size (i.e. reduced estimation precision), we also conducted permutation testing in which we iteratively removed 42 scans *not* originally analyzed in the Gee et al. study before group-level modeling.

g. **Outliers:** As preregistered, scans where amygdala reactivity estimates were > 3 standard deviations from the mean were excluded from analysis. In addition, to account for the possibility of remaining outliers, additional models were fit using a Student's t-distribution (rather than a Gaussian) for the likelihood function. A gamma(4,1) prior was used for the parameter for degrees of freedom (v). Such models, because they model the outcome variable with a heavy-tailed t-distributions (i.e. t-distributions with few degrees of freedom), have been demonstrated to be robust to outliers (Gelman & Hill, 2006; Kurz, 2019).

h. **Group-level models:** ROI estimates from each scan were submitted to multilevel linear regression models to estimate group-level effects. Age was grand mean-centered and modeled as a continuous variable, and all models included a covariate for mean framewise displacement (Power et al., 2012). In multiverse analyses, we included models with all combinations of additional covariates for task run (coded as a binary variable indicating first run versus second/third run) and scanner (coded as a binary variable for scanner 1 versus scanner 2). In addition, we included models with an additional quadratic term for age to explore potential non-linear age-related changes. We included models both with and without participant-specific random slope terms, but all models included participant-specific intercepts.

i. **Within-participant change vs. between-participant differences:** To ask whether any observed age-related changes in amygdala reactivity were due to true developmental growth (i.e. longitudinal change within the same participant across timepoints) versus between-participant differences, we conducted a separate smaller multiverse analysis. In this analysis, we included separate model parameters for ‘within-participant’ (i.e. mean-centered within participants) and ‘between-participant’ (i.e. grand mean-centered) age effects, with random effects for within-participant age. These analyses included preprocessing pipelines using both FSL and C-PAC, GLMs run in both FSL and AFNI, and both native and MNI space amygdala t-stat estimates.

*Longitudinal Amygdala Model Syntax*

As described above, we modeled longitudinal age-related changes in amygdala reactivity using several different specifications with the brms package. R syntax for the 9 specifications for longitudinal models is shown below. Each model (plus equivalent models with normal likelihood functions instead) was applied to all 156 preprocessing pipelines for a total of 2808 models.

```
# 1: Linear, no exclusions
modLinear = brm(reactivity ~ age_centered + motion + (age_centered|participant), data = ., cores = 2, chains = 4,
family = 'student', prior = prior(gamma(4, 1), class = nu))

# 2: Linear, exclude previously studied participants
modLinearExclude = brm(reactivity ~ age_centered + motion + (age_centered|participant), data = dplyr::filter(.,
is.na(prev_studied)), cores = 2, chains = 4, family = 'student', prior = prior(gamma(4, 1), class = nu))

# 3: Quadratic, exclude previously studied participants
modQuadraticExclude = brm(reactivity ~ poly(age_centered,2, raw = TRUE) + motion + (age_centered|participant),
data = dplyr::filter(., is.na(prev_studied)), cores = 2, chains = 4, family = 'student', prior = prior(gamma(4, 1), class =
nu))

# 4: Quadratic, no exclusions
modQuadratic = brm(reactivity ~ poly(age_centered,2, raw = TRUE) + motion + (age_centered|participant), data = .,
cores = 2, chains = 4, family = 'student', prior = prior(gamma(4, 1), class = nu))
```

```
# 5: Linear + scanner covariate, no exclusions
modLinearScanner = brm(reactivity ~ age_centered + motion + scanner + (age_centered|participant), data = ., cores
= 2, chains = 4, family = 'student', prior = prior(gamma(4, 1), class = nu))

# 6: Linear + block covariate, no exclusions
modLinearBlock = brm(reactivity ~ age_centered + motion + blockBin + (age_centered|participant), data = ., cores =
2, chains = 4, family = 'student', prior = prior(gamma(4, 1), class = nu))

# 7: Quadratic + block + scanner covariates, no exclusions
modQuadraticBlockScanner = brm(reactivity ~ poly(age_centered,2, raw = TRUE) + motion + blockBin + scanner +
(age_centered|participant), data = ., cores = 2, chains = 4, family = 'student', prior = prior(gamma(4, 1), class = nu))

# 8: Linear + block + scanner covariates, no exclusions
modLinearBlockScanner = brm(reactivity ~ age_centered + motion + blockBin + scanner +
(age_centered|participant), data = ., cores = 2, chains = 4, family = 'student', prior = prior(gamma(4, 1), class = nu)))

# 9: Linear without participant-varying slopes, no exclusions
modLinearNoRandomSlopes = brm(reactivity ~ age_centered + motion + (1|participant), data = ., cores = 2, chains =
4, family = 'student', prior = prior(gamma(4, 1), class = nu)))
```

*Parametrization of within-participant change versus between-participant differences*

In efforts to better discriminate truly longitudinal within-participant changes from between-participant differences, we created alternate model parametrizations for a subset of specifications (bilateral amygdala only, t-statistic amygdala reactivity estimates) for both the fear > baseline and neutral > baseline contrast. For each model, we included two separate terms for age. The first term represented between-participant differences and was grand mean-centered (such that this term was equal to 0 at age 11.9 years; the mean age across all participants and scans); the second, representing within-participant change, was centered within participants (such that this term was equal to 0 at the mean age of each participant across visits). We allowed only the second within-participant change term to vary across participants, as follows:

```
# Within vs. between model (reactivity)
brm(reactivity ~ age_grand_mean_centered + age_centered_within_participant +
            motion + (age_centered_within_participant|participant))
```

After fitting these model formulations to the subset of preprocessing specifications, we examined the posterior distributions of resulting grand mean-centered (between-participant) and within-participant centered (within-participant) parameters in a smaller specification curve (Figure 2D). In addition, we also examined approximate leave-one-out cross-validated $R^2$ using the loo_R2() function from the brms package (Bürkner, 2017). This metric uses the posterior likelihood to provide an adjusted $R^2$ metric that estimates predictive performance.

*Within-person similarity of reactivity voxel-wise statistical maps across multiverse forks*

We asked whether, for each given scan, whether different pipelines yielded similar voxel-wise patterns of estimates in a within-scan analysis. To explore how changes in preprocessing impacted scan-level statistical results for the fear > baseline, neutral > baseline, and fear > neutral contrasts, we computed image similarities between t-statistic maps for each scan across all pipelines. Similarities were calculated using product-moment correlations of 3d images (using AFNI 3ddot; https://afni.nimh.nih.gov/pub/dist/doc/program_help/3ddot.html) for the same scan after transformation to MNI space at 2mm (isotropic) resolution, and across all voxels in the brain. We computed similarities across all pairwise comparisons of pipelines preprocessed using C-PAC, as well the similarity of all C-PAC pipelines to the preregistered FSL pipeline. In addition to whole-brain similarity, we also computed similarity statistics in the same way for all voxels within the bilateral amygdala mask from the Harvard Oxford atlas.

*Between-scan associations between amygdala reactivity estimates across specifications*

We also asked whether different pipelines yielded similar relationships between amygdala betas across scans in a between-scan analysis. This analysis examined whether between-scan relationships among amygdala reactivity estimates were preserved across preprocessing specifications for each contrast. To accomplish this, we computed rank-order correlations between amygdala-reactivity estimates (t-tstats, bilateral amygdala only) between preprocessing specifications for each contrast. Because this analysis was focused on examining whether scan-level differences were preserved across specifications, correlations were conducted across all scans from all participants without taking into account the nesting of repeated observations within participants across sessions.

*Dependence of amygdala reactivity findings on previous work*

42 scans from the first timepoint were previously analyzed as a 'discovery set' by Gee et al (2013). Thus, we were concerned that analyses of amygdala reactivity including all data (including these scans) might be biased to find stronger effects of negative age-related change due to the pre-selection of the amygdala for showing an effect within part of the current sample (see Kriegeskorte et al., 2009). Indeed, age-related change in amygdala reactivity for the fear > baseline contrast was stronger and more negative in specifications using all data, compared to when excluding these 42 scans (see sFigure 25). However, such stronger effects without exclusion may also have been due to the larger sample size. To address this, we conducted permutation tests across several pipelines to assess differences in age-related change estimates after exclusion of these 42 scans versus 42 other randomly-drawn scans. To conserve computational resources, we fit these models using maximum likelihood with the lme4 R package (Bates et al., 2019), rather than fully Bayesian inference. Approximate 95% confidence intervals were constructed from these models by computing the interval ±2 standard errors from the maximum likelihood estimate.

#### Within-scan changes in amygdala reactivity across trials

*Changes in amygdala reactivity across trials: multiverse details (Table 2, Aim 2)*

a. **Quantification of change across trials:** We conducted separate multiverse analyses using several different methods for measuring change in reactivity across trials.

    1. *Slopes:* For each scan we performed a rank-order correlation between trial number and amygdala betas corresponding to each trial number (separately for fear and neutral trials). These correlation coefficients representing slopes for linear changes in reactivity across trials were then submitted to group-level models.

    2. *Trial halves:* We split trials into the first half (trials 1-12) and second half (trials 13-24) for fear and neutral faces respectively, and modeled age-related change in amygdala reactivity separately for each half of trials at the group level. These models included half x age interaction terms to specifically estimate whether age-related change in amygdala reactivity differed in the first versus second half of trials.

    3. *Single-trial models:* We constructed single-trial models with age x trial number interaction terms and crossed random effects for age (random intercept and age effects for each participant) and trial number (random intercepts and linear slopes for trial for each scan).

b. **Global signal subtraction:** It is possible that changes in amygdala reactivity across trials may reflect temporal trends in the whole-brain 'global signal', rather than differing amygdala reactivity specifically. To correct for this, we included pipelines with a global signal correction using post-hoc distribution centering of reactivity for each trial. Post-hoc distribution centering consisted of subtracting the mean beta estimate across all voxels from each voxel such that the distribution of beta estimates for each respective trial was centered at 0 (mean-centering).

c. **Amygdala ROI:** Only Harvard-Oxford anatomically-defined amygdala ROIs in MNI space were used in these analyses. Analyses of bilateral, left, and right amygdala ROIs were each included.

d. **Group-level models:** Age was grand mean-centered and modeled as a continuous variable, and all models included a covariate for mean framewise displacement. In multiverse analyses, we included models with all combinations of additional covariates for task run (coded as a binary variable indicating first run versus second/third run) and scanner (coded as a binary variable for scanner 1 versus scanner 2). In addition, we included models with an additional quadratic term for age. All models were formulated to be robust to outliers as described above.

*Longitudinal model syntax for models of within-scan change in amygdala reactivity*

Example brms model formulas are shown below for each of the different types of models for within-scan changes in amygdala reactivity.

```
# Slope model (method 1)
slope_model_ = brm(slope ~ age_centered + motion + (age_centered | participant),
        data = ., cores = 2, chains =4, family = 'student',
        prior = prior(gamma(4, 1), class = nu))


# Trial half model (method 2)
half_model = brm(reactivity ~ age_centered*half + motion +
            (age_centered|participant) + (1 | scan),
        data = ., cores = 2, chains =4, family = 'student',
        prior = prior(gamma(4, 1), class = nu))


# Single trial model (linear trial term, method 3)
```

```
single_trial_linear = brm(reactivity ~ age_centered*trial +
       motion + (age_centered | participant ) + (trial | scan),
       data = ., cores = 2, chains =4, family = 'student',
       prior = prior(gamma(4, 1), class = nu))


# Single trial model (discrete trial term, method 3)
single_trial_discrete = brm(reactivity ~ age_centered*trial_discrete +
       motion + (age_centered | participant ) + (trial | scan),
       data = ., cores = 2, chains =4, family = 'student',
       prior = prior(gamma(4, 1), class = nu))
```

### Amygdala—mPFC functional connectivity analyses

*Amygdala—mPFC gPPI analyses: multiverse details (Table 2, Aim 3)*

a. **Preprocessing:** We ran all gPPI analyses using preregistered preprocessing pipelines in FSL. No gPPI analyses were run on data preprocessed using C-PAC.

b. **Amygdala gPPI seed:** All gPPI analyses used the anatomically-defined Harvard-Oxford bilateral amygdala mask as a seed region. We extracted mean timecourses from the amygdala from the preprocessed BOLD data to use as the seed regressor.

c. **Deconvolution step**: In all pipelines interaction terms between the amygdala seed timeseries and both fear and neutral face regressors were constructed following generalized form (gPPI; McLaren et al., 2012). Before multiplication of the seed and stimulus timeseries, however, some pipelines included an additional step such that the seed timeseries was deconvolved to recreate the seed 'neuronal' timeseries using AFNI 3dTfitter (https://afni.nimh.nih.gov/pub/dist/doc/program_help/3dTfitter.html). To most closely match pipelines run by Gee et al. (2013), we did not up-sample the seed timeseries before using 3dTfitter, and applied no regularization to the regression solver. This deconvolution step (often included in AFNI and SPM PPI analyses, but not FSL) has been suggested to allow the PPI regressors to better approximate task-modulated connectivity at the level of the neuronal response, rather than after being filtered by the hemodynamic response function (Di & Biswal, 2017; Gitelman et al., 2003). Following deconvolution, the resulting timeseries was multiplied with the fear and neutral face regressors, then re-convolved with the HRF for entry into the GLM. For pipelines not including a deconvolution step, seed timeseries were multiplied by stimulus regressors that had already been convolved with the HRF.

d. **First-Level gPPI GLM:** First-level GLMs for gPPI analyses were constructed in FSL almost identically to those used in preregistered amygdala reactivity analyses, with the addition of the amygdala seed timeseries regressor and gPPI terms for both fear and neutral faces. gPPI GLMs included 24 head motion parameters and had TRs with framewise displacement >.9mm down-weighted to 0. As with amygdala reactivity analyses, we extracted both t-statistics and contrast beta estimates (COPEs) for the fear > baseline, neutral > baseline, and fear > neutral contrasts, although the fear > baseline contrast was of primary interest for the current study.

e. **mPFC ROI definition:** We preregistered constructing an mPFC ROI containing 120 voxels centered at the peak coordinates reported by Gee at al. (2013) for age-related change in fear > baseline gPPI (Tailarach 2,32,8; or MNI 3,35,8). However, after discovery that this ROI heavily overlapped the corpus collosum, we instead constructed three spherical ROIs with 5mm radii, the first centered at the above peak coordinates, the second shifted slightly anterior, and the third shifted slightly ventral relative to the second (see Figure 4). Lastly, we also used a large mask encompassing the 'whole vmPFC', taken from Mackey & Petrides (2014). All masks used are available on OSF (https://osf.io/hvdmx/). For each scan, we calculated mean gPPI beta estimates and t-statistics for each of these four ROIs for submission to group-level models.

f. **Group-level models, outliers, and previously analyzed scans:** Multiverse gPPI analyses included identical decision points to amygdala reactivity analyses with respect to group-level modeling, dealing with outliers, and use of previously analyzed scans.

*Amygdala—mPFC BSC analyses: multiverse details (Table 2, Aim 3)*

a. **Preprocessing:** We conducted all BSC analyses using preregistered preprocessing pipelines in FSL.
b. **GLMs:** We used beta estimates from LSS GLMs fit separately to each trial (as described on p. 12 of the main text) for BSC analyses.
c. **Amygdala ROI definitions:** Only Harvard-Oxford amygdala ROIs in MNI space were used in these analyses. Analyses of bilateral, left, and right amygdala ROIs were each included.
d. **mPFC ROI definitions:** For BSC analyses, we used the same mPFC ROIs as with gPPI analyses (see Table 2, Aim 3).
e. **Global signal subtraction:** We included BSC pipelines both with and without the global signal subtraction step previously described (post-hoc distribution centering).
f. **Beta-series correlations:** For each respective amygdala—mPFC ROI pair (12 pairs in total for 3 amygdala x 4 mPFC), we extracted mean beta estimates for each ROI for each trial, then calculated product-moment correlations between the timeseries across trials (neutral and fear separately) for both regions (Di et al., 2020). These correlation coefficients were then submitted to group-level models.
g. **Group-level models:** Multiverse BSC analyses included identical decision points to gPPI analyses with respect to group-level modeling, with the exceptions that all BSC models were formulated to be robust to outliers (using Student’s t likelihood functions), and we did not exclude previously analyzed scans from any BSC analysis pipelines.

*Longitudinal models for functional connectivity*

Longitudinal models for gPPI used the same 9 specifications as previously described (with R syntax) above for amygdala reactivity. BSC models were similar, except that all BSC models used t-distributed likelihood functions and no BSC models excluded previously studied participants because BSC analyses had not previously been conducted with these data. Thus, there were a total of 7 longitudinal model specifications for BSC analyses.

*Rank-order associations between gPPI & BSC amygdala—mPFC FC*

In addition to examining age-related changes in amygdala—mPFC FC using both gPPI and BSC methods, we asked whether between-scan differences in scan-level FC estimates were similar across methods. To accomplish this, we computed rank-order correlations between amygdala—mPFC FC estimates using both gPPI (with deconvolution and without) and BSC (with global signal subtraction and without) methods for all four mPFC ROIs. Because this analysis aimed to investigate the scan-level similarity of FC estimates across method, we computed correlations across all scans for all participants.

*Effects of up-sampling/lasso regularization parameters during deconvolution on gPPI regressors*

The AFNI documentation for the 3dTfitter program comes with the warning, “Deconvolution is a tricky business, so be careful out there! ... Experiment with different parameters to make sure the results in your type of problems make sense” (https://afni.nimh.nih.gov/pub/dist/doc/program_help/3dTfitter.html). While our initial choices of deconvolution parameters were guided by previous work done with the same dataset, we explored the impact of different parameters on the resulting gPPI regressors of interest. We systematically varied whether to up-sample the seed amygdala timeseries to a sampling resolution of 10Hz (effective TR = 0.1s), and whether to apply no regularization versus an L1 penalty to the deconvolution solution parameters. We then compared within-scan similarity of gPPI regressors by computing product-moment correlations between all generated gPPI regressors for a given scan. We also computed equivalent correlations between gPPI regressors and the seed timeseries.

*Within-person similarity of voxel-wise statistical maps for amygdala FC from gPPI pipelines*

We asked whether, for each given scan, whether different pipelines yielded similar voxel-wise patterns of estimates in a within-scan analysis. To explore the degree to which scan-level statistical maps for gPPI contrasts were impacted by whether a deconvolution step was included in the pipeline, we computed similarity statistics for fear > baseline, neutral > baseline, and fear > neutral gPPI contrast t-statistic maps between the pipelines with versus without deconvolution. Similarities were calculated using product-moment correlations of 3d images (using AFNI 3ddot; https://afni.nimh.nih.gov/pub/dist/doc/program_help/3ddot.html) after transformation to MNI space at 2mm (isotropic) resolution, and across all voxels in the brain. In addition to whole-brain similarity, we also computed similarity statistics in the same way for all voxels within all four mPFC ROIs previously described.

*Within-person similarity between gPPI & BSC amygdala FC*

Given previous evidence that gPPI and BSC connectivity methods detect similar differences in connectivity between task contrasts (Di et al., 2020), we asked whether this was true for the fear > neutral faces contrast, specifically for amygdala FC. To accomplish this, we computed mean FC with the Harvard-Oxford bilateral amygdala for each ROI in the Harvard-Oxford cortical and subcortical atlases (https://neurovault.org/collections/262/). For gPPI, mean FC was computed as the mean t-statistic over all voxels in each ROI for the fear > neutral contrast. For BSC, mean FC was computed as the product-moment correlation between the average timeseries of the bilateral amygdala and each ROI. We thus compiled a vector representing amygdala FC with 62 ROIs (we removed all subcortical ROIs representing white matter, ventricles, or entire hemispheres of cerebral cortex) across the brain for gPPI (both with and without deconvolution) and BSC pipelines (both with and without global signal subtraction (GSS)). For each scan, we computed product-moment correlations between each of these vectors as a measure of the similarity of amygdala FC with the rest of the brain across pipelines. We also computed within-person similarity for BSC amygdala connectivity for pipelines with versus without global signal subtraction.

### Associations between amygdala—mPFC circuitry & separation anxiety:

*Amygdala—mPFC circuitry & separation anxiety: multiverse details (Table 2, Aim 5)*

a. **Amygdala reactivity measures:** We used t-statistic estimates from the bilateral amygdala (both native and MNI space) for fear > baseline, neutral > baseline, and fear > neutral contrasts as the predictor of interest.
b. **Amygdala reactivity slope measures:** For analyses of change in amygdala reactivity over trials, we used the slope calculated across fear and neutral trials, respectively, as previously described. We used a bilateral amygdala mask (in MNI space) to define amygdala reactivity slopes, and included pipelines both with and without global signal correction.
c. **Amygdala—mPFC FC measures:** We used t-statistic estimates for gPPI between bilateral amygdala (in MNI space) and all four mPFC ROIs previously described, as well as BSC estimates for FC between the same regions. We included gPPI estimates both with and without a deconvolution step, and BSC estimates with and without global signal

correction. We submitted estimates to group-level models for fear, neutral, and fear > neutral contrasts.

d. **Separation anxiety outcomes:** Analyses were run for each of three separation anxiety outcomes: scores from the RCADS separation anxiety subscale, and both raw and standardized t-scores from the SCARED separation anxiety subscale.

e. **Group-level models:** In all models, separation anxiety outcomes were modeled using multilevel linear regressions with crossed random effects for age and brain measure. All separation anxiety models were formulated to be robust to outliers (using Student's t likelihood functions), and we did not exclude previously analyzed scans from any analysis pipelines.

*Longitudinal model syntax for amygdala—mPFC circuitry & separation anxiety*

Longitudinal models for associations between amygdala—mPFC circuitry and separation anxiety included participant-varying slopes for the respective brain measure included as a predictor in each model, as well as for age.

```
# Separation anxiety model
brm(separation_anxiety ~ brain_measure + age_centered + motion + (brain_measure + age_centered|participant,
chains = 4, cores = 4,
          family = 'student', prior = prior(gamma(4, 1), class = nu))
```

**Estimating impacts of specific forked decision points on age-related change estimates**

To explore impacts of different analytical decision points (or the impacts of 'taking different forks' in the path from beginning to end of the analysis) on age-related change estimates, we submitted point estimates for linear age-related change (posterior medians) from each model to separate Bayesian linear regression models. Models were fit using the rstanarm package (Gabry et al., 2019), and included each decision point (one-hot encoded if there were more than 2 options) as a binary predictor of the point estimates for age-related change. Following modeling, we plotted posterior distributions and 95% posterior intervals for each parameter, representing effects of each decision point conditional on all others. Example syntax of one model for amygdala reactivity decision points is below.

```
decision_point_model = stan_glm(data = sca_frame, estimate ~ tstat + quadratic +
                    random_slopes + ctrl_scanner + ctrl_block + exclude_prev +
                    amyg_right + amyg_left + amyg_high_sig + amyg_low_sig +
                    native_space + motion_reg6 + motion_reg18 +
                    hrf_2gamma + highpass + robust,
                    cores = 4, chains = 4)
```

### Supplemental Results

#### Behavior: Supplemental Results

*Behavioral Task Performance*

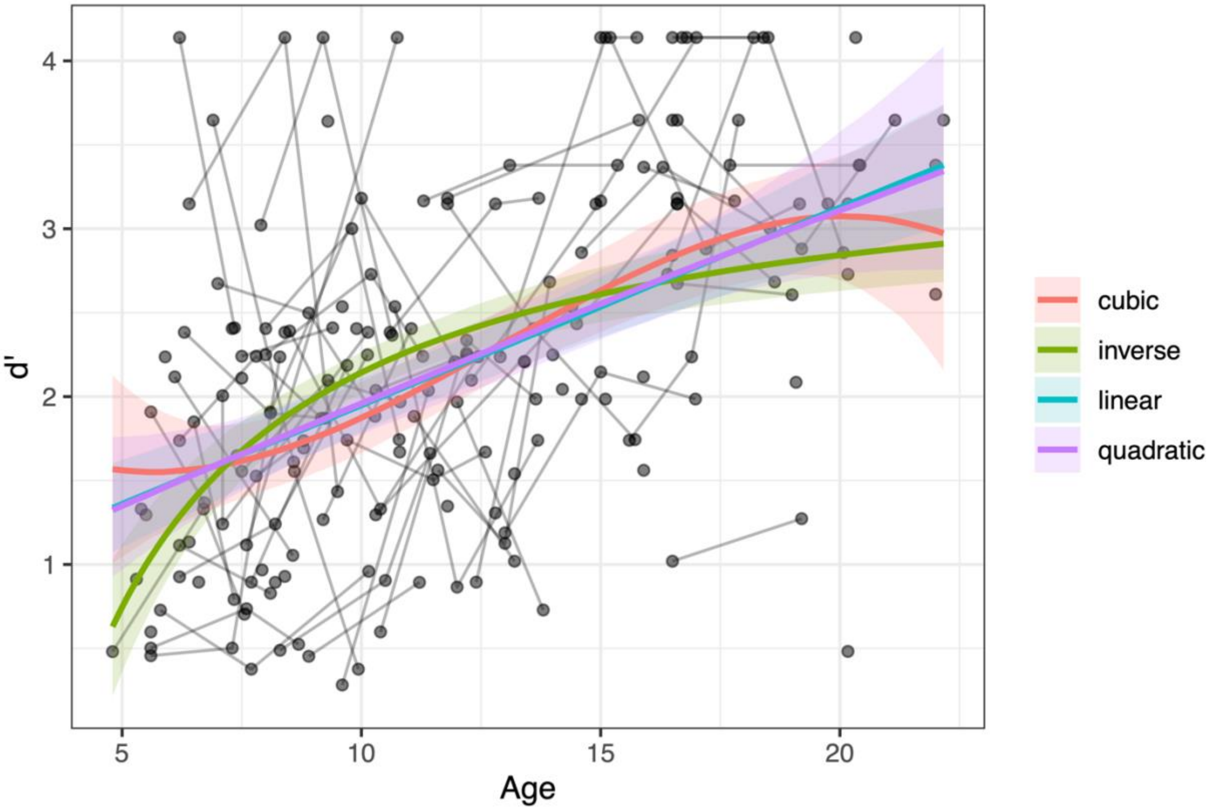


**sFigure 3:** d' as a function of age.
d' for task performance as a function of age, modeled using linear, quadratic, cubic, and inverse age longitudinal models. Points display performance for one participant at one timepoint, black lines connect estimates from participants with multiple study visits, and colored lines with shaded area represent fitted model predictions and 95% posterior intervals.

We modeled task performance with d' (using a correction for extreme proportions; Hautus, 1995) as a function of linear, quadratic, cubic, and inverse age trends (sFigure 3). Models indicated age-related improvements in d' without notable quadratic or cubic change. The ICC for d' scores was estimated to be 0.31 (95% PI [0.04, 0.51]). We also modeled accuracy (probability of a correct response on any trial, with hits and correct rejections coded as 1, misses and false alarms coded as 0), hits, and false alarms using single-trial multilevel logistic regression models, and found similar age-related increases in task performance (sFigure 4). Even at the youngest ages, task performance was well above chance levels. We also modeled reaction time as a function of age using linear, quadratic, and cubic models. Along with general age-related decreases in reaction times to neutral faces ('hits'), quadratic and cubic models indicated that the most age-related change in average reaction times occurred at the younger end of the age range, between approximately 4-11 (sFigure 4, bottom right panel).

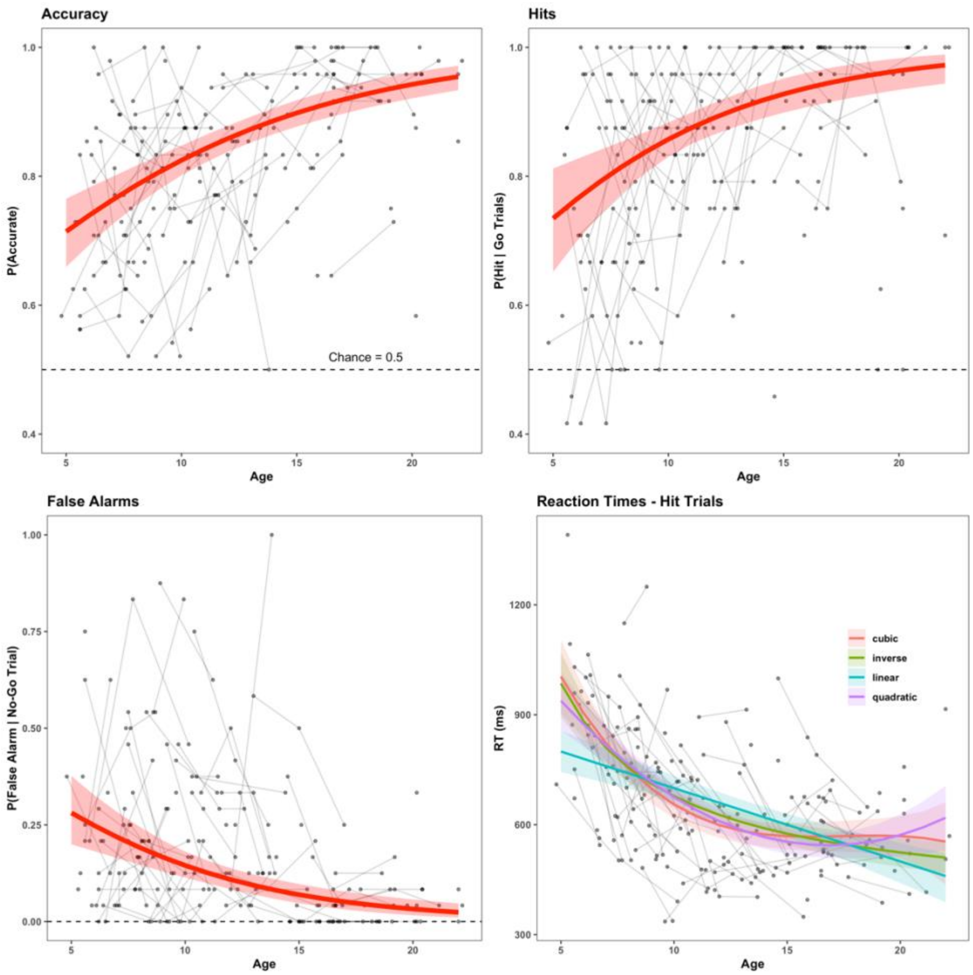


**sFigure 4:** Task performance metrics as a function of age
Age-related change in accuracy (top left), hits (top right), false alarms (bottom left), and reaction times during go trials (bottom right). Points display summarized performance for one participant at one timepoint (e.g. the proportion accurate across trials during that run), black lines connect

performance summaries from participants with multiple study visits, and colored lines with shaded area represent fitted trial-by-trial model predictions and 95% posterior intervals.

*Head Motion*

Head motion, as measured by mean framewise displacement estimates derived from MCFLIRT (Jenkinson et al., 2002) decreased with age ($\hat{\beta}$=-0.04, 95% PI [-0.05, -0.02]) among all participants with available task fMRI data (sFigure 5). Among only included participants with <= 40 TRs with FD < 0.9mm, mean framewise displacement also decreased with age, although somewhat less strongly ($\hat{\beta}$=-0.02, 95% PI [-0.02, -0.01]).

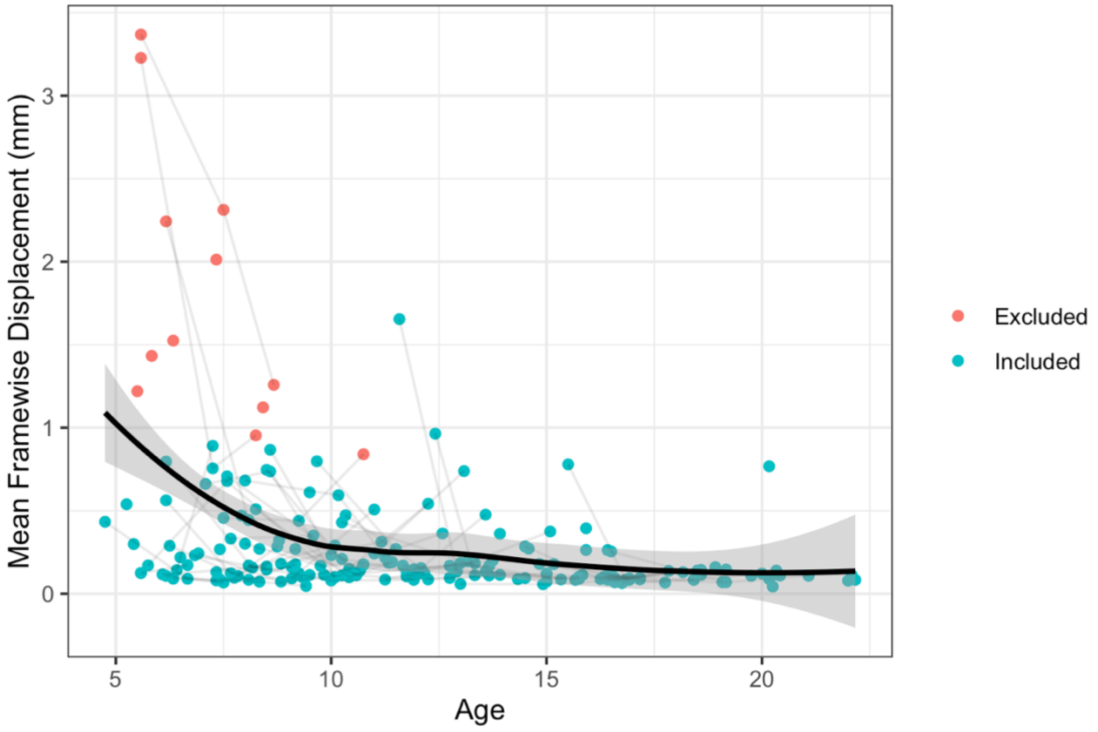


**sFigure 5:** Mean framewise displacement as a function of age (years)

### Amygdala Reactivity: Supplemental Results

*Group Mean Amygdala Reactivity*

We separate specification curves to estimate group averages in amygdala reactivity for the fear > baseline, neutral > baseline, and fear > neutral contrasts. To preserve computational resources, all models for mean amygdala reactivity were fit using the lme4 R package with intercepts allowed to vary by participant. All specifications for the fear > baseline contrast (sFigure 6) and most specifications for the neutral > baseline contrast (sFigure 7) resulted in positive amygdala reactivity with a confidence interval distinct from zero, indicating robust average amygdala reactivity to faces of both emotions. In addition, most specifications found higher amygdala reactivity for fear compared to neutral faces (fear > neutral contrast, sFigure 8). Higher average amygdala reactivity for fear > neutral faces may have been due to either the face emotions or the fact that participants were instructed to press for neutral faces and withhold button press for fear faces.

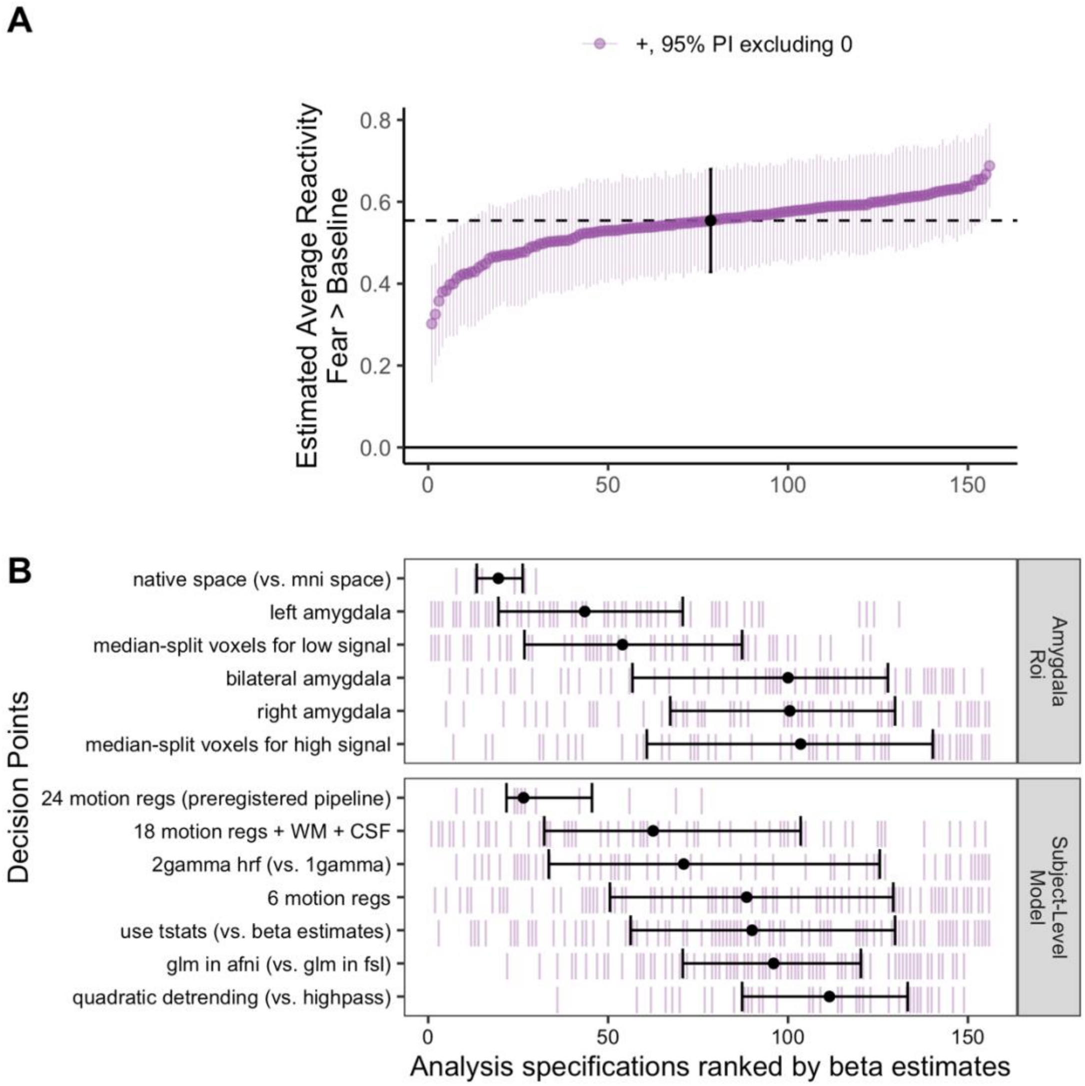


**sFigure 6:** Specification curve for mean fear > baseline amygdala reactivity
**A:** Points represent estimated mean amygdala reactivity for each specification, and lines represent corresponding 95% posterior intervals. **B:** Variables on the y-axis represent analysis choices, corresponding lines indicate that a choice was made, and blank space indicates that the choice was not made in a given analysis.

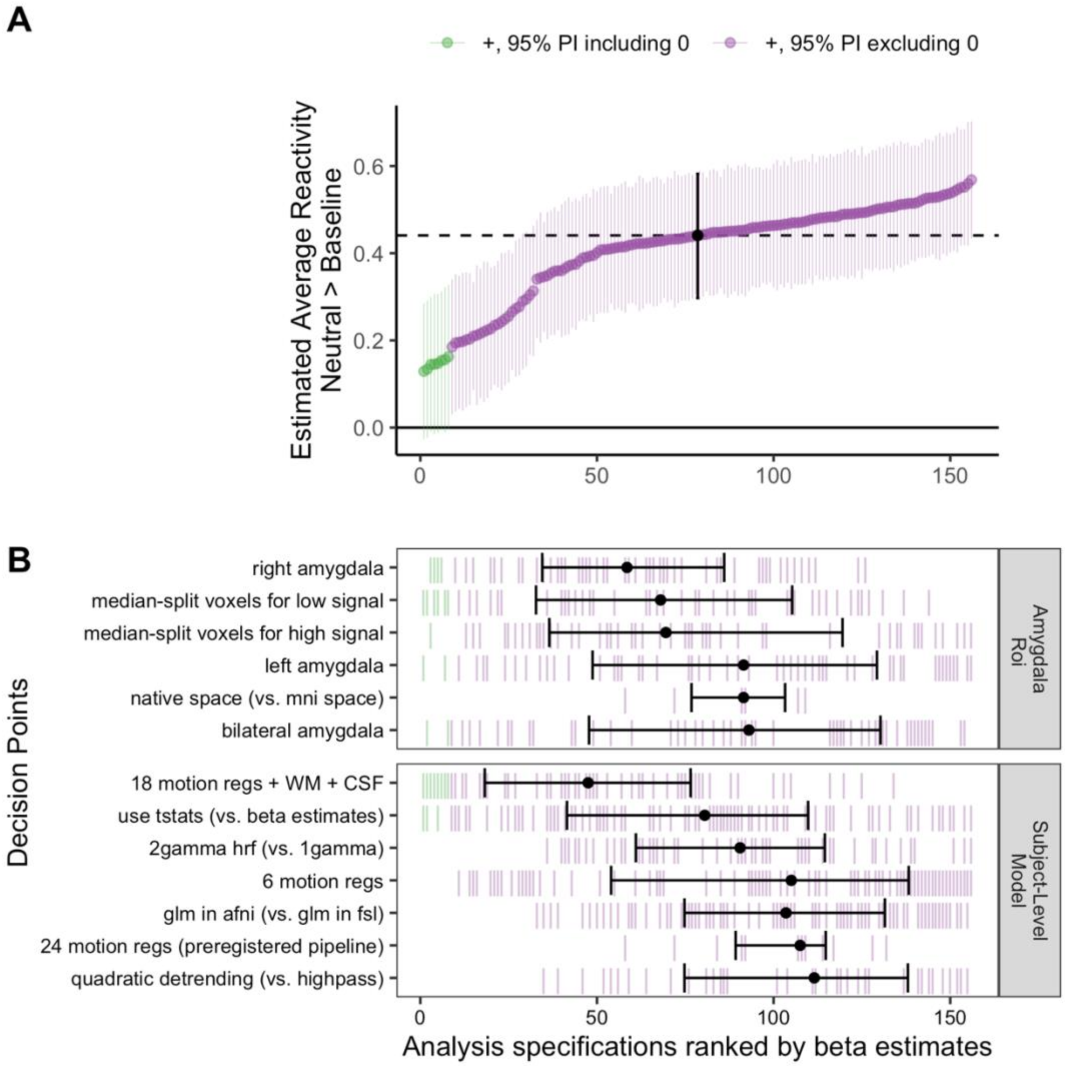


**sFigure 7:** Specification curve for mean neutral > baseline amygdala reactivity
**A:** Points represent estimated mean amygdala reactivity for each specification, and lines represent corresponding 95% posterior intervals. **B:** Variables on the y-axis represent analysis choices, corresponding lines indicate that a choice was made, and blank space indicates that the choice was not made in a given analysis.

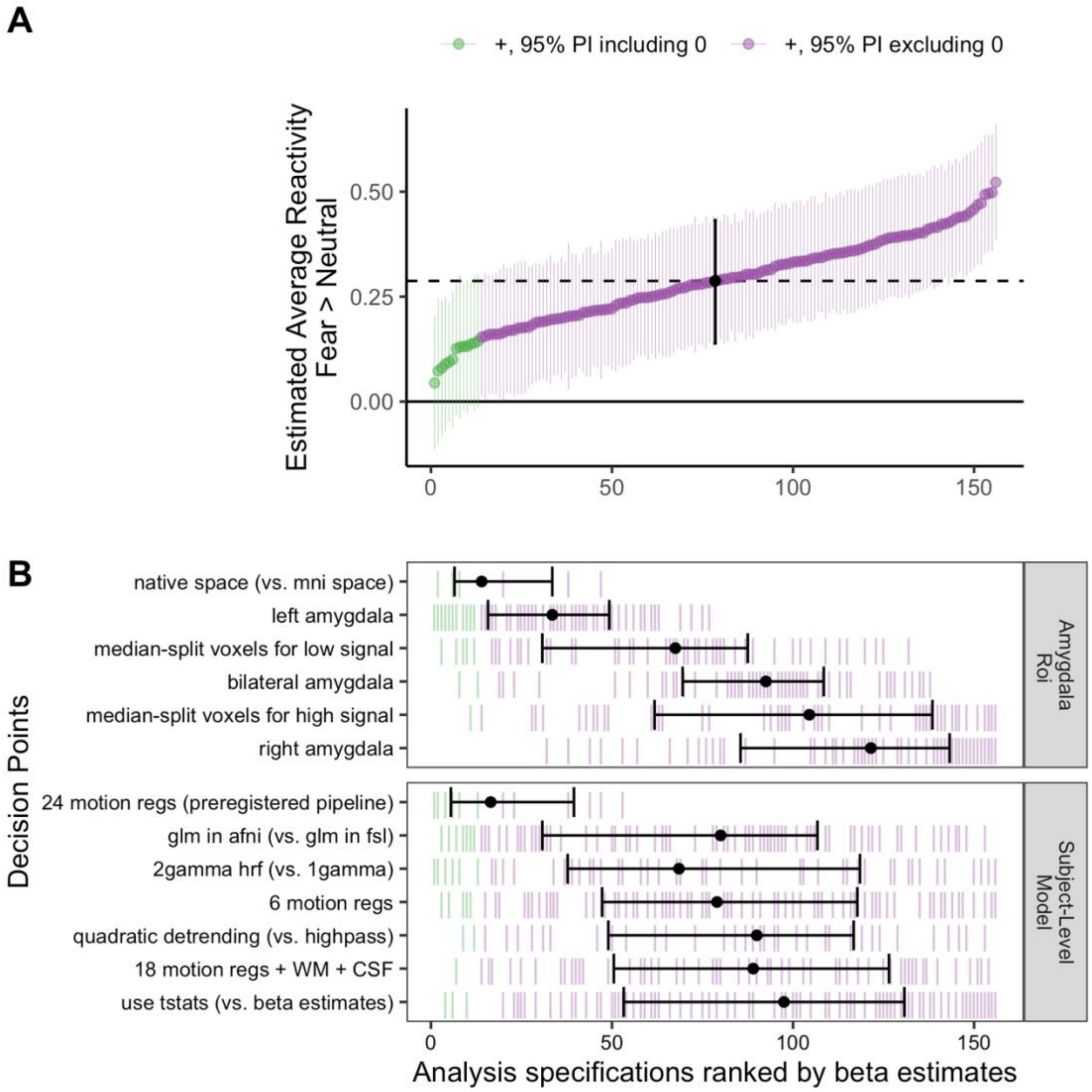


**sFigure 8:** Specification curve for mean fear > neutral amygdala reactivity
**A:** Points represent estimated mean amygdala reactivity for each specification, and lines represent corresponding 95% posterior intervals. **B:** Variables on the y-axis represent analysis choices, corresponding lines indicate that a choice was made, and blank space indicates that the choice was not made in a given analysis.

*Multiverse analyses of age-related changes in amygdala reactivity*

In addition to constructing specification curves for age-related change in amygdala reactivity for the fear > baseline contrast as reported in the main manuscript (see Figure 2), we constructed parallel specification curves for the neutral > baseline and fear > neutral contrasts. Generally, age-related change findings for the neutral > baseline contrast were similar to, but slightly weaker than, fear > baseline: while 98.9% of specifications found negative age-related change, only 42.4% of specifications estimated negative age-related change distinguishable from 0 (see sFigure 9). No pipelines found age-related change in fear > neutral amygdala reactivity distinguishable from 0 (see sFigure 10).

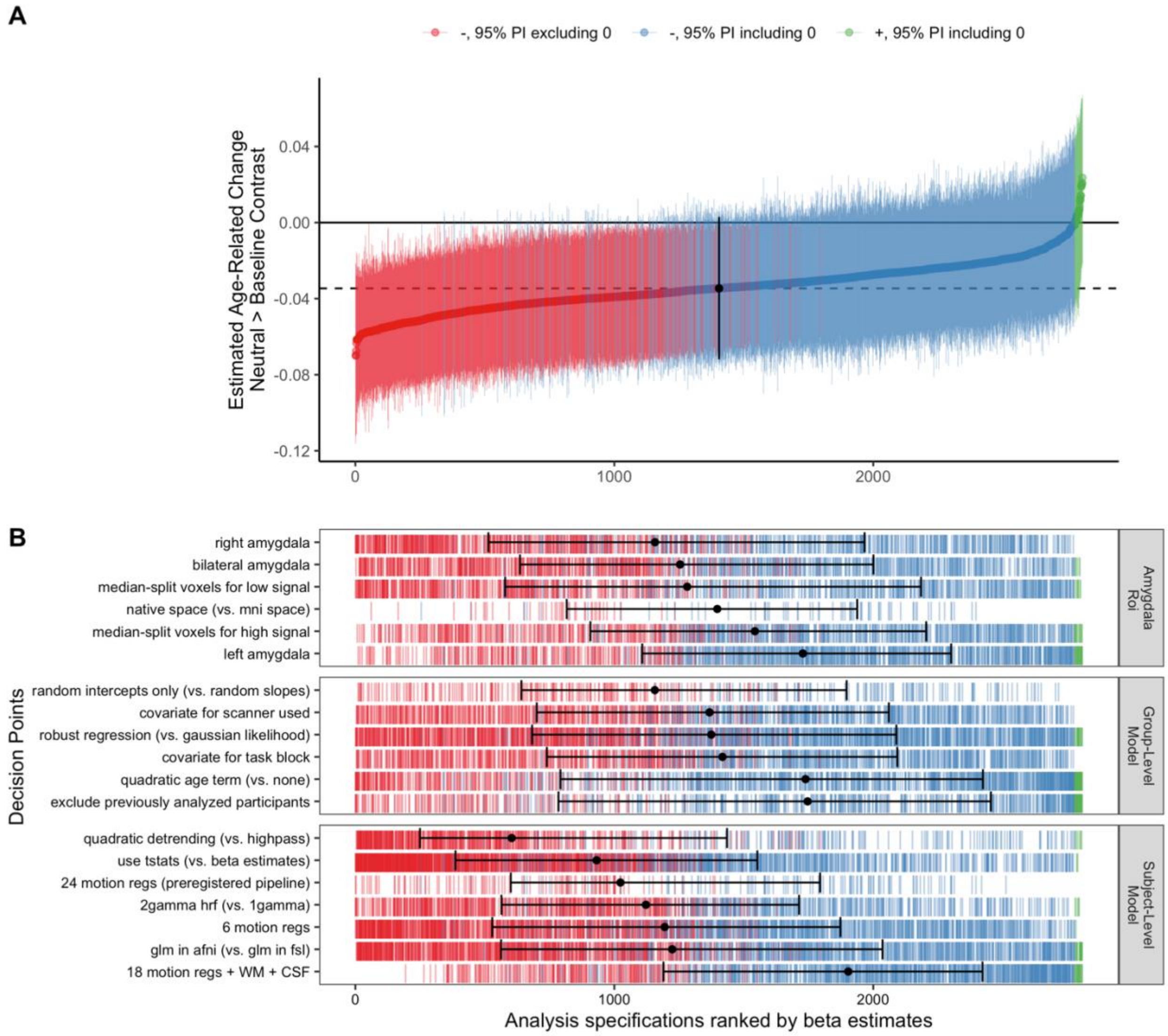

**sFigure 9:** Specification curve for age-related change in neutral > baseline amygdala reactivity **A:** Points represent estimated linear age-related change in amygdala reactivity for each specification, and lines represent corresponding 95% posterior intervals. **B:** Variables on the y-axis represent analysis choices, corresponding lines indicate that a choice was made, and blank space indicates that the choice was not made in a given analysis.

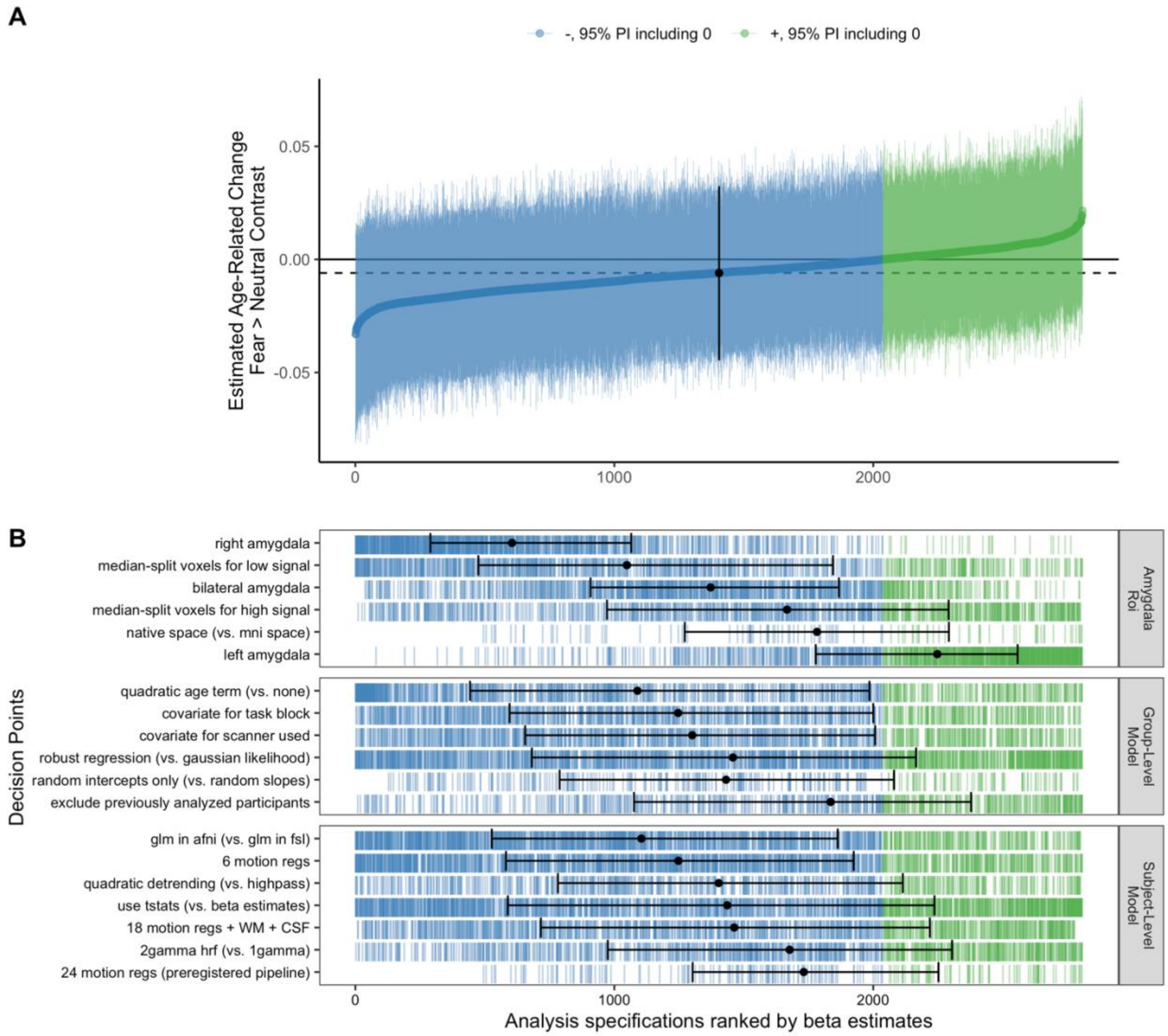


**sFigure 10:** Specification curve for age-related change in fear > neutral amygdala reactivity **A:** Points represent estimated linear age-related change in amygdala reactivity for each specification, and lines represent corresponding 95% posterior intervals. **B:** Variables on the y-axis represent analysis choices, corresponding lines indicate that a choice was made, and blank space indicates that the choice was not made in a given analysis.

*Impacts of pipeline choices on age-related change estimates for amygdala reactivity*

While the goals of the present study were not to precisely quantify impacts of specific pipeline decision points, we explored impacts of all decision points on estimates of age-related changes in amygdala reactivity for each contrast. Specifically for the fear > baseline contrast (see sFigure 11), age-related change was somewhat stronger (more negative) for specifications using a right amygdala region compared to a bilateral region, and weaker for specifications using a left amygdala region. Most notably, specifications excluding the 42 participants previously studied (Gee et al., 2013) found weaker age-related change.

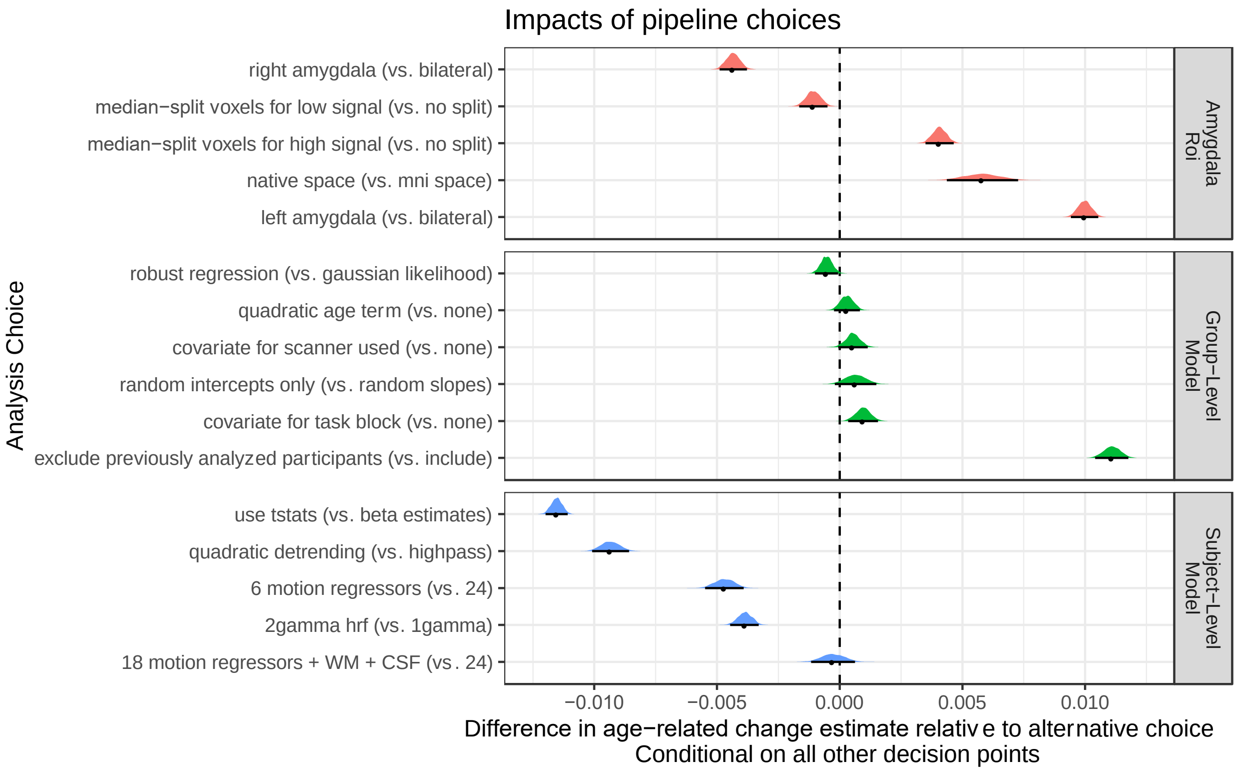


**sFigure 11:** Fork impacts on age-related change for fear > baseline amygdala reactivity Posterior distributions and 95% posterior intervals are shown, representing the average difference in linear age-related change estimates relative to the alternative choice. As most specifications find negative age-related change, negative values indicate more strongly negative change, and positive values indicate weaker change.

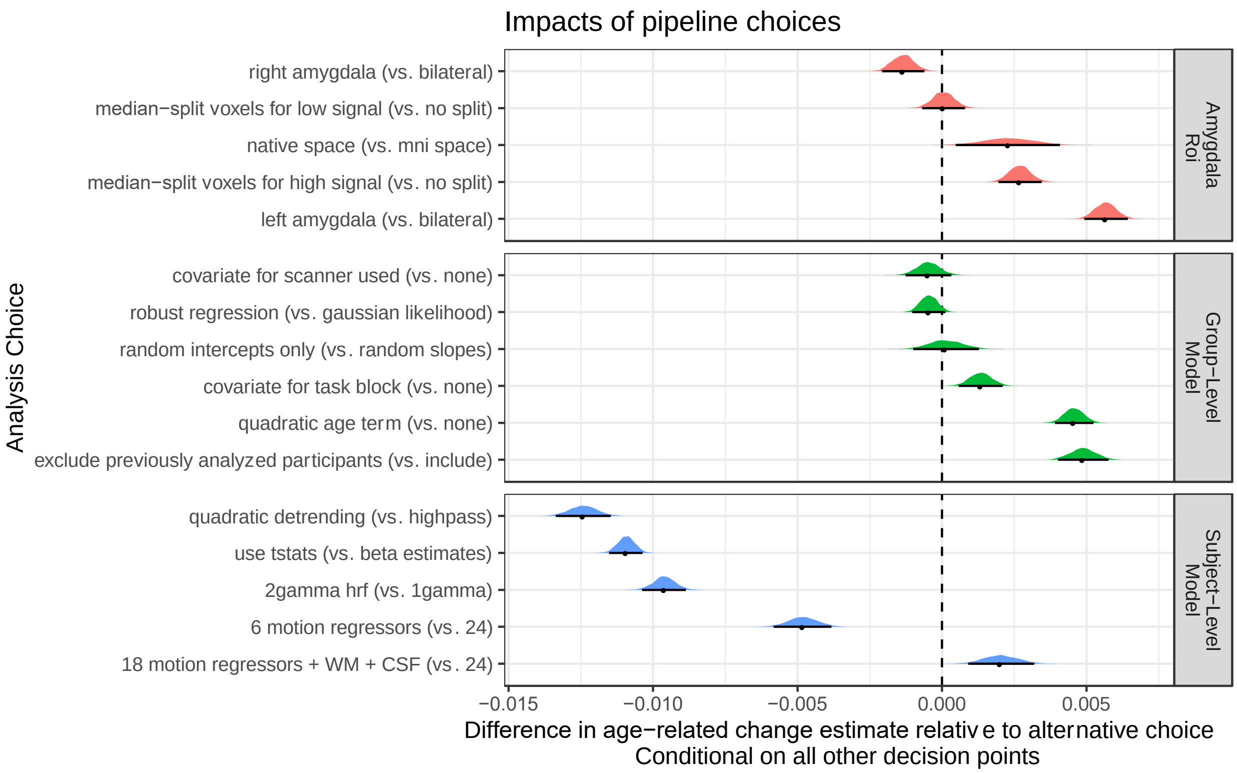


**sFigure 12:** Fork impacts on age-related change for neutral > baseline amygdala reactivity Posterior distributions and 95% posterior intervals are shown, representing the average difference in linear age-related change estimates relative to the alternative choice. As most specifications find negative age-related change, negative values indicate more strongly negative change, and positive values indicate weaker change.

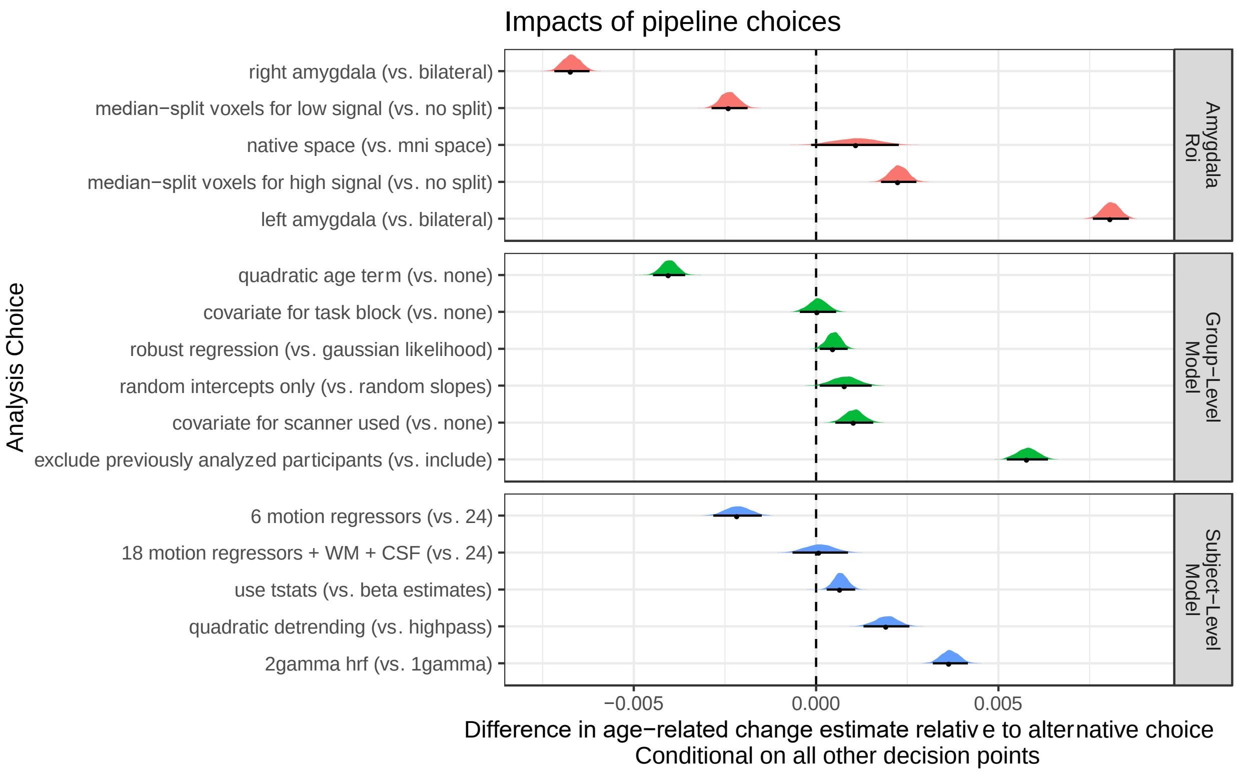


**sFigure 13:** Fork impacts on age-related change for fear > neutral amygdala reactivity Posterior distributions and 95% posterior intervals are shown, representing the average difference in linear age-related change estimates relative to the alternative choice. Negative values indicate more negative change, and positive values indicate more positive change.

*Nonlinear Changes in Amygdala Reactivity*

In addition to linear age-related change, we also included quadratic age terms in some model specifications for somewhat more flexible modeling of longitudinal trajectories of amygdala reactivity. We constructed specification curves for quadratic age-related change parameters across all models including quadratic terms for the fear > baseline (sFigure 14), neutral > baseline (sFigure 15), and fear > neutral (sFigure 16) contrasts. Across all contrasts, very few specifications estimated quadratic terms distinguishable from 0 under a 95% posterior interval. Further, quadratic fits were varied in sign for each contrast, such that some quadratic models estimated developmental 'peaks' while others estimated 'troughs' in amygdala reactivity. When model predictions from separate specifications were plotted as individual 'spaghetti', there was not clear consensus in quadratic trajectories for any contrast (as there was for linear change for the fear > baseline and neutral > baseline contrasts, see sFigure 17). Thus, while the current study may not have been adequately powered to estimate quadratic age-related change, we did not find consistent evidence for either peaks or troughs in amygdala reactivity between ages 4-22.

Because inverse age models may be particularly useful in characterizing rapid change early in development, we also fit such models such that amygdala reactivity was estimated as a function of 1/age. Models were fit using maximum likelihood, and fits indicated age-related decreases in the fear > baseline and neutral > baseline contrasts (sFigure 18). However, such inverse age models used here can characterize rapid change earlier in development but not later in development by design (i.e. they are formulated to capture deceleration in adolescence and stabilization in young adulthood; Luna et al., 2021).

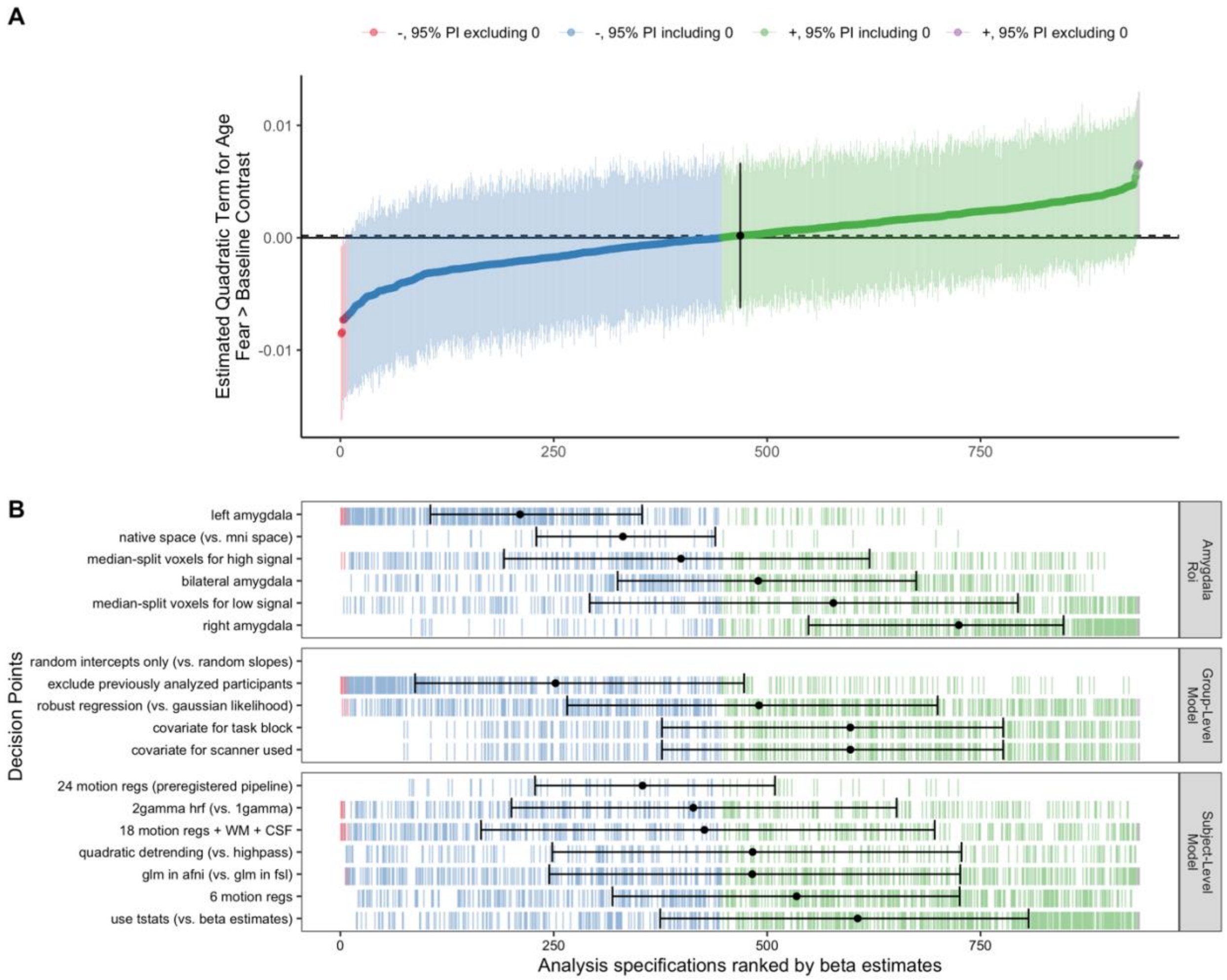


**sFigure 14:** Spec. curve for quadratic age-related change in fear > baseline amygdala reactivity **A:** Points represent estimated quadratic age-related change in amygdala reactivity for each specification, and lines represent corresponding 95% posterior intervals. **B:** Variables on the y-axis represent analysis choices, corresponding lines indicate that a choice was made, and blank space indicates that the choice was not made in a given analysis.

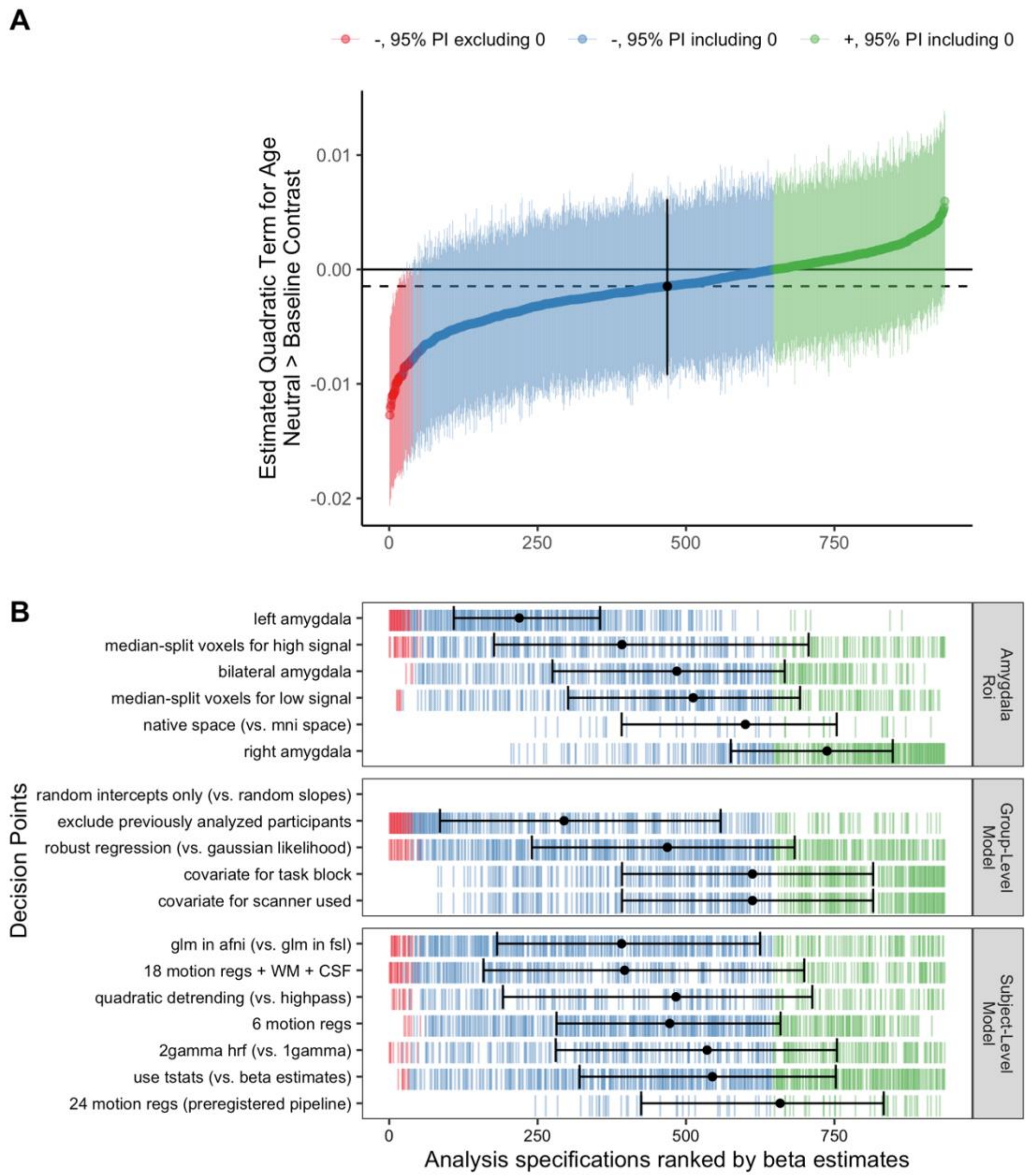


**sFigure 15:** Spec. curve for quadratic age-related change in neutral > baseline amygdala reactivity **A:** Points represent estimated quadratic age-related change in amygdala reactivity for each specification, and lines represent corresponding 95% posterior intervals. **B:** Variables on the y-axis represent analysis choices, corresponding lines indicate that a choice was made, and blank space indicates that the choice was not made in a given analysis.

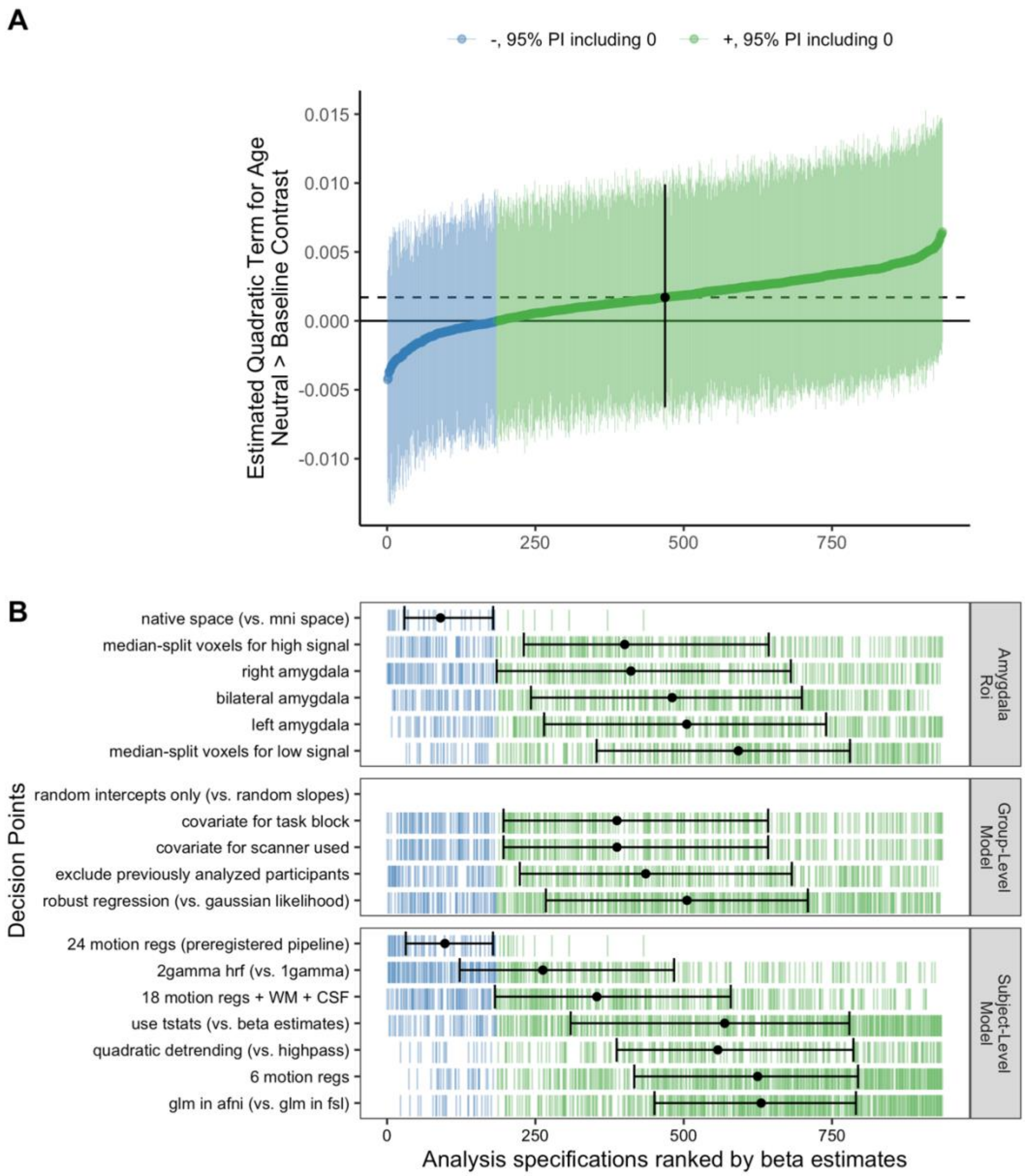


**sFigure 16:** Spec. curve for quadratic age-related change in fear > neutral amygdala reactivity **A:** Points represent estimated quadratic age-related change in amygdala reactivity for each specification, and lines represent corresponding 95% posterior intervals. **B:** Variables on the y-axis represent analysis choices, corresponding lines indicate that a choice was made, and blank space indicates that the choice was not made in a given analysis.

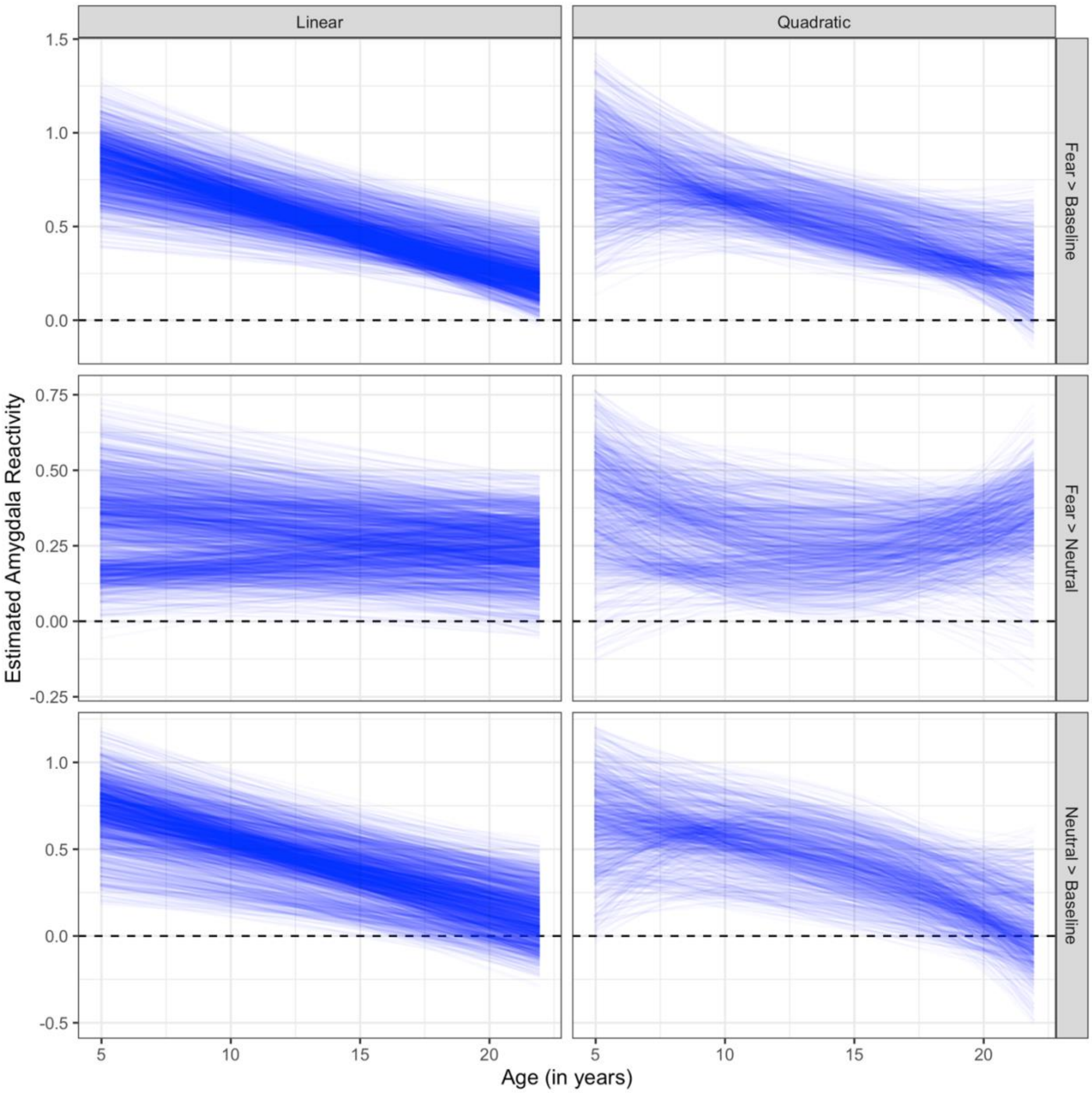


**sFigure 17:** Predictions of age-related change across linear and quadratic model specifications. Each blue line represents 1 specification, with age on the x-axis and estimated amygdala reactivity on the y-axis. For quadratic models (right panel), some specifications found convex change while other indicated concave change.

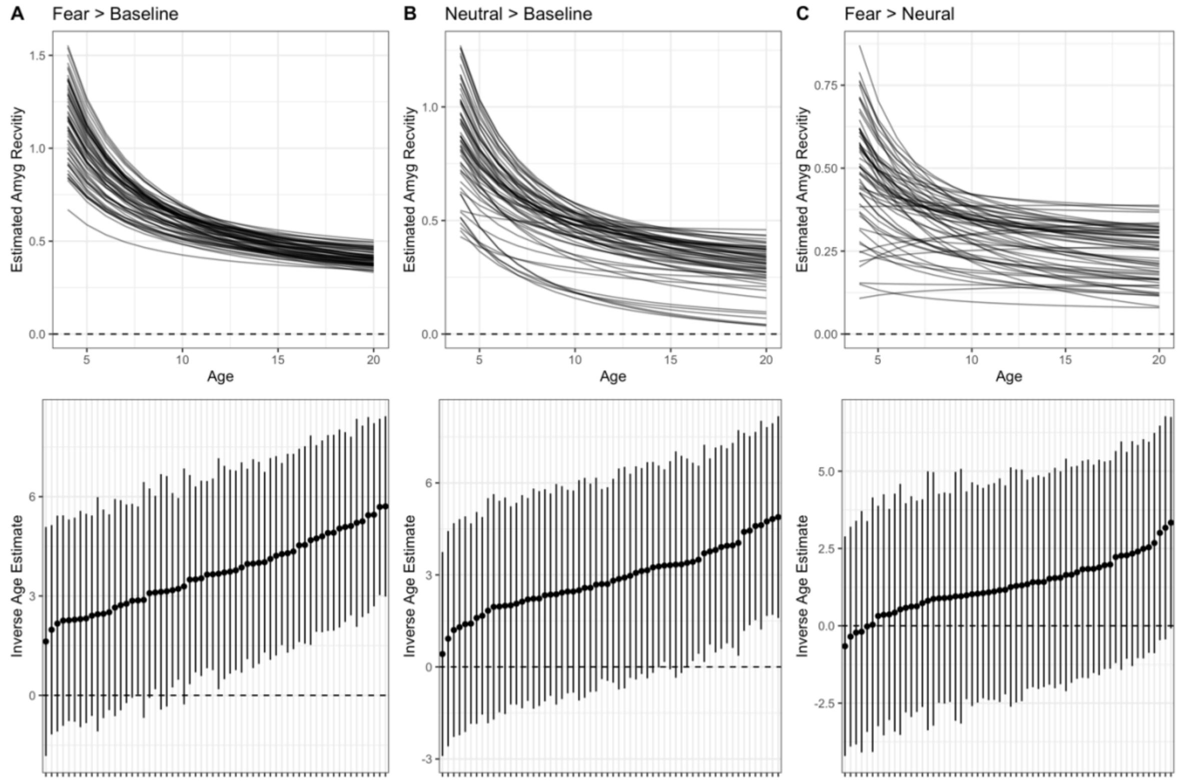


**sFigure 18**: Inverse age models for amygdala reactivity
Spaghetti plots for fitted model predictions (top) and specification curves (bottom) for inverse age estimates for models of amygdala reactivity. Positive parameter fits for inverse age indicate decrease in amygdala reactivity as a function of age. Panels represent the fear > baseline contrast (**A, left**), neutral > baseline contrast (**B, center**), and the fear > neutral contrast (**C, right**)

*Between-participant age associations versus within-participant age-related change*

We constructed a smaller specification curve of models parametrized to differentiate between-participant age associations from within-participant age-related changes in amygdala reactivity. While only between-participant terms indicated consistent associations with age, we also examined, for each specification, whether between-participant and within-participant age terms *differed* from one another. We estimated the differences between such terms through calculating the distribution of paired differences in posterior draws between the two terms, and summarizing using the median and 95% quantiles (to construct a 95% posterior interval). Overall, we found that despite the higher estimation precision for between-participant age associations reported in the main text (see Figure 2), within-participant and between-participant estimates did not consistently differ within most models for fear > baseline or neutral > baseline amygdala reactivity (sFigure 19A). In addition, $R^2$ metrics calculated with approximate leave-one-out cross-validation were close to 0 for all models, indicating predictive performance close no better than chance (sFigure 19B).

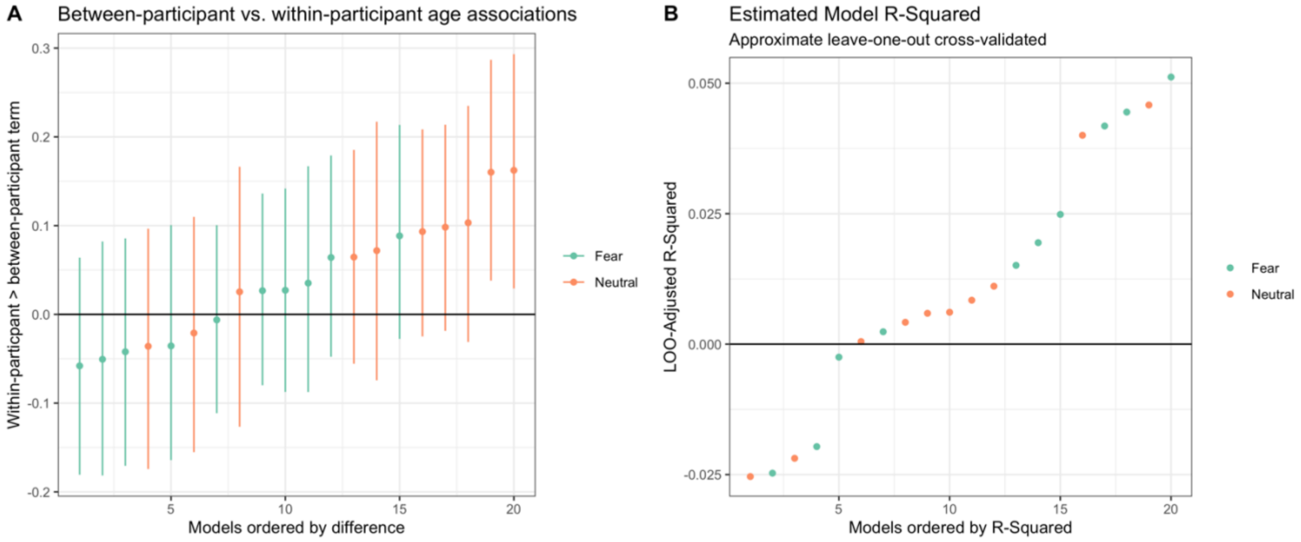


**sFigure 19:** Differences between within-participant and between-participant terms for age-related associations with amygdala reactivity
**A.** Estimated differences between within-participant and between-participant terms for age-related associations with amygdala reactivity. **B.** Approximated leave-one-out cross-validated $R^2$ scores for each specification separately parametrizing within-participant and between-participant terms. $R^2$ values below 0 indicate cross-validated performance poorer than that expected under the “null” model.

*Within-person similarity of voxel-wise amygdala reactivity statistical maps across forks*

We sought to understand whether different preprocessing pipelines, for each given scan, yielded similar voxel-wise patterns of estimates in a within-scan analysis. To understand which preprocessing steps influenced reactivity estimates most, we computed the voxel-wise similarity of statistical maps (t-statistics) for the *same scans* across preprocessing specifications. Thus, for each pair of pipelines, we computed 1 similarity value for each scan for the whole brain, and

1 similarity value for each scan for the bilateral amygdala. While similarity (product-moment correlation across all brain voxels) was positive for most comparisons between the preregistered pipeline (all FSL) and pipelines using C-PAC preprocessing, similarity did vary across scans such that for any comparison, some scans were highly different across specifications (i.e. near-zero or negative correlation values). In general, statistical maps for both the whole brain and the bilateral amygdala from the preregistered FSL pipeline were more similar to pipelines using C-PAC preprocessing + FSL GLMs compared to pipelines using C-PAC preprocessing + AFNI GLMs (see sFigure 20). In addition, within specifications with C-PAC preprocessing, similarity across pipelines was somewhat higher, especially within pipelines using the same GLM software or nuisance regressors (see sFigure 21). Relatively higher similarity among of specifications with C-PAC preprocessing was likely to do the common registration shared by all such pipelines (as opposed to differing registrations from the preregistered FSL pipeline).

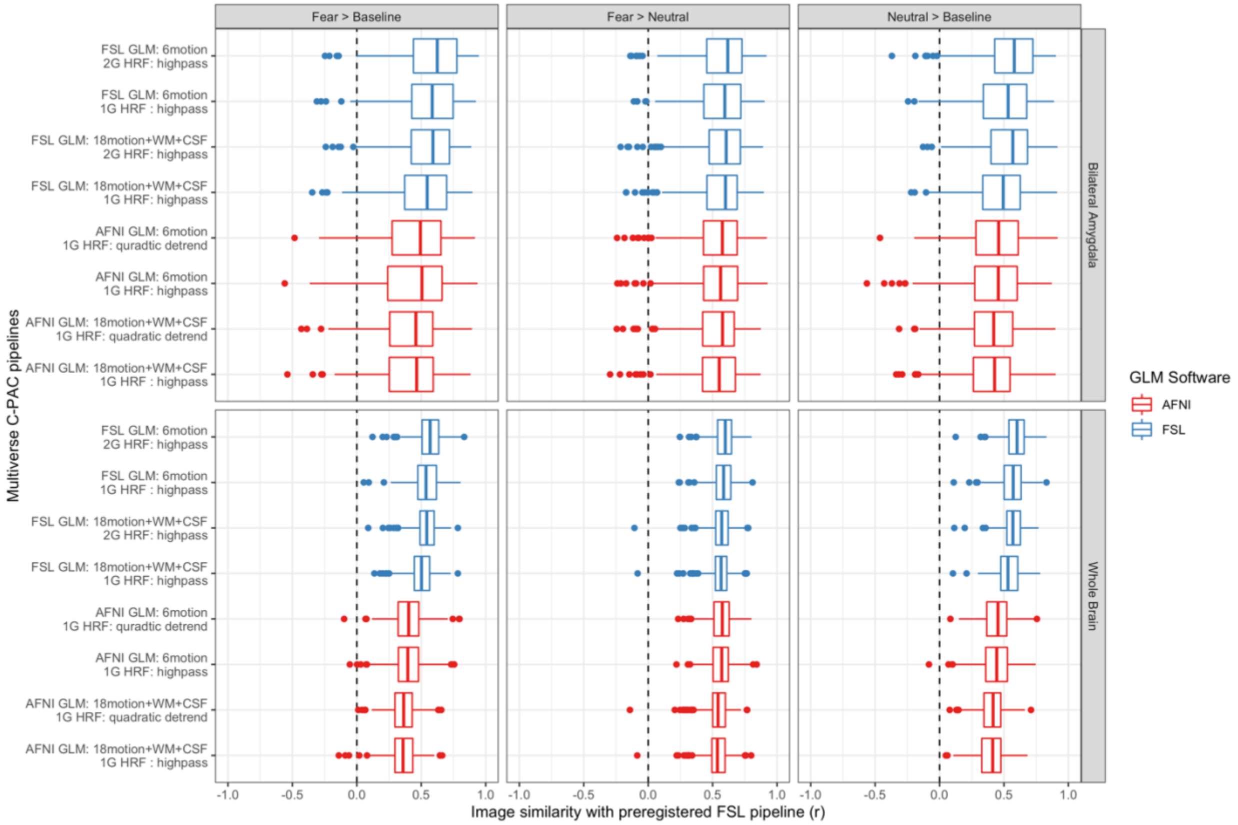


**sFigure 20.** Voxelwise image similarity for amygdala reactivity contrasts between the preregistered FSL pipeline and C-PAC pipelines.
The x-axis indicates product-moment correlations across all brain voxels (bottom panel) and bilateral amygdala voxels (top) for the preregistered FSL pipeline with the pipelines indicated on the y-axis. Similarity is shown for the fear > baseline (left), fear > neutral (middle), and neutral > baseline (right) contrasts.

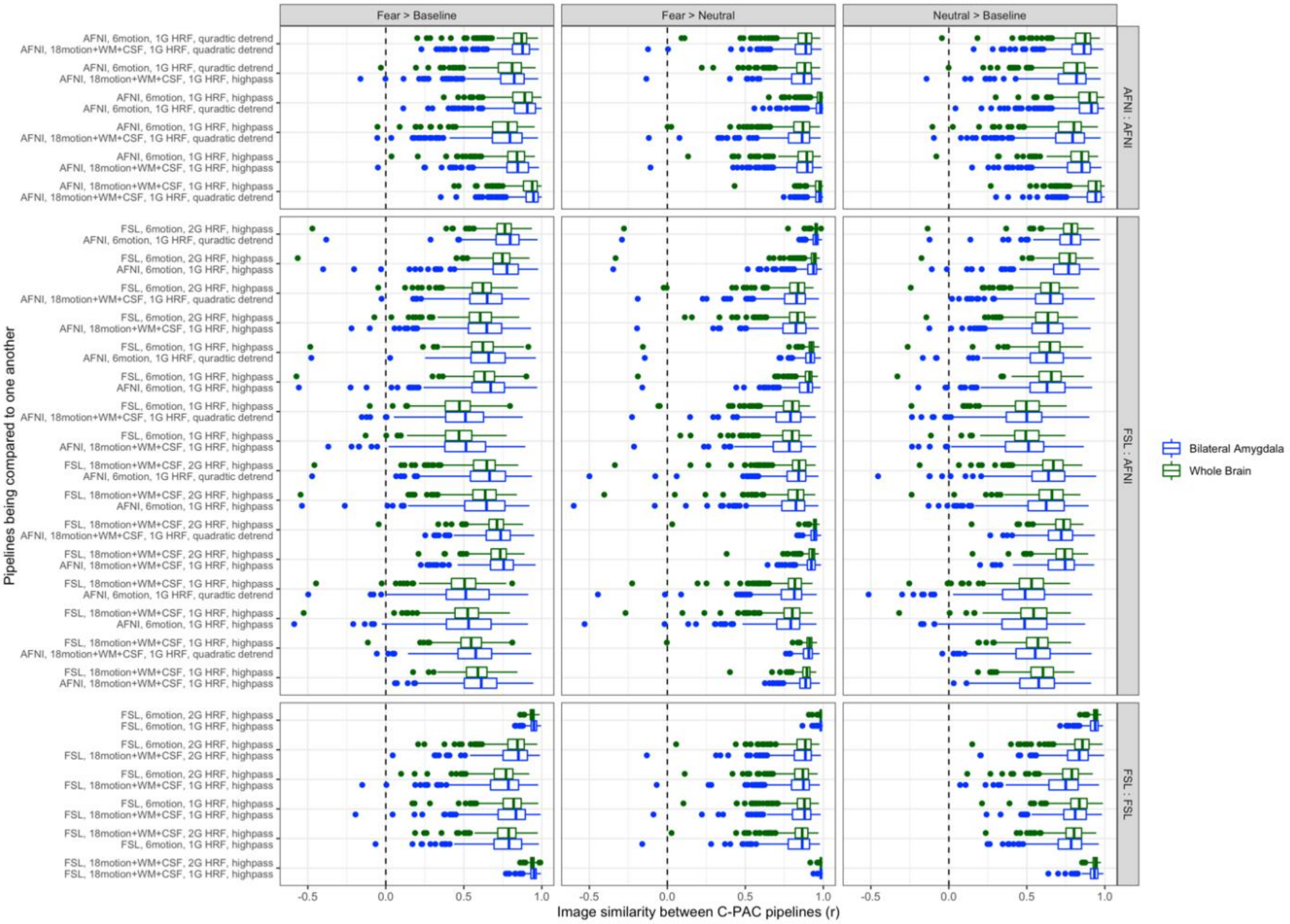


**sFigure 21:** Voxelwise similarity for amygdala reactivity contrasts between all C-PAC pipelines. The x-axis indicates product-moment correlations across all brain voxels (green) and bilateral amygdala voxels (blue) within each scan for all pairwise comparisons of C-PAC pipelines indicated on the y-axis. Similarity is shown for the fear > baseline (left), fear > neutral (middle), and neutral > baseline (right) contrasts. Pipeline comparisons are organized into comparisons between two pipelines with AFNI GLMs (top), one pipeline with an AFNI GLM and the other with FSL (middle), and two pipelines with FSL GLMs (bottom).

*Between-scan correlations of amygdala reactivity estimates across specifications*

To ask whether relative relationships between scans for amygdala reactivity were preserved across preprocessing specifications, we computed correlations between scan-level estimates of mean amygdala reactivity across pairs of pipelines (for bilateral amygdala estimates, t-statistics) for each contrast. For each pair of preprocessing pipelines, we computed the rank-order correlation between vectors of bilateral amygdala reactivity estimates (1 datapoint per scan per pipeline). While correlations were all positive and mostly strong (r >= .7) for most pairs of pipelines for the fear > baseline (sFigure 22) and neutral > baseline (sFigure 23) contrasts, correlations between the most disparate preprocessing pipelines were often much weaker (e.g. from 0.4-0.6). Thus, the between-scan relationships between amygdala reactivity estimates were only somewhat weakly preserved across such pipelines. On the other hand, for the fear > neutral contrast, all pairwise comparisons of pipelines yielded higher (r >=.7) between-scan correlation values (sFigure 24).

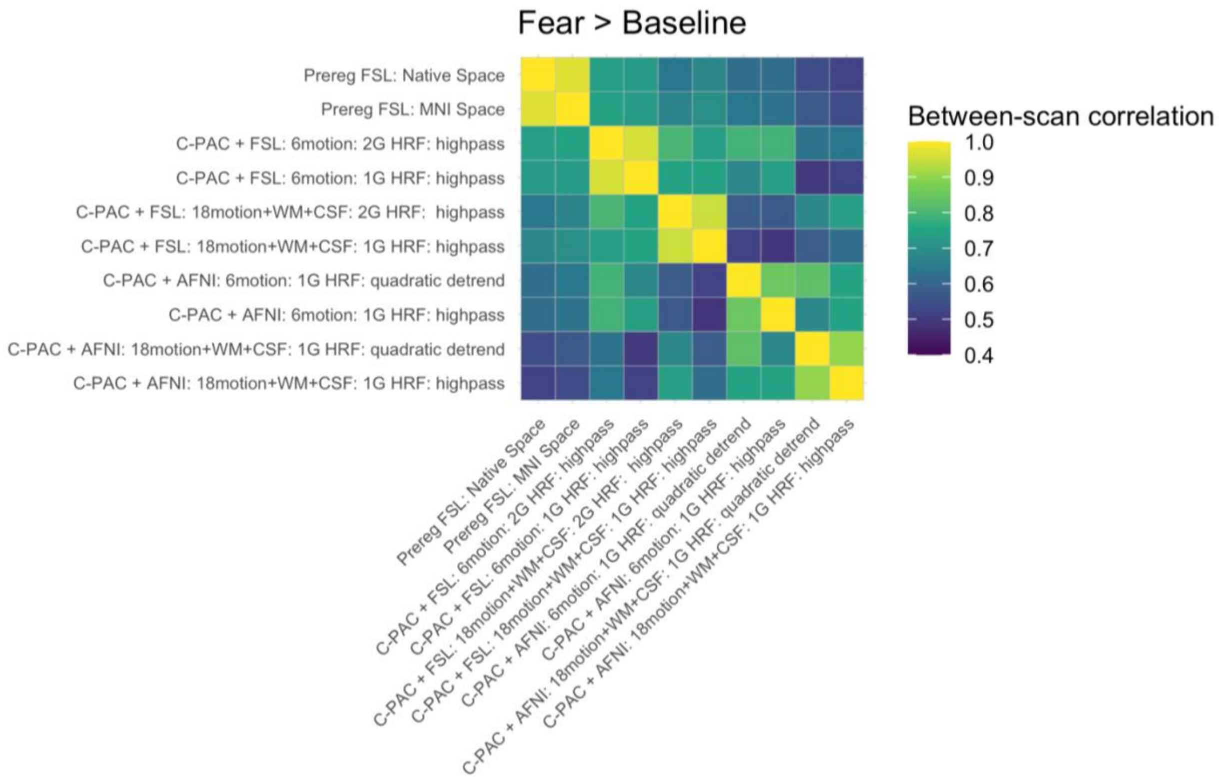


**sFigure 22:** Between scan correlations for amygdala reactivity pipelines for fear > baseline.

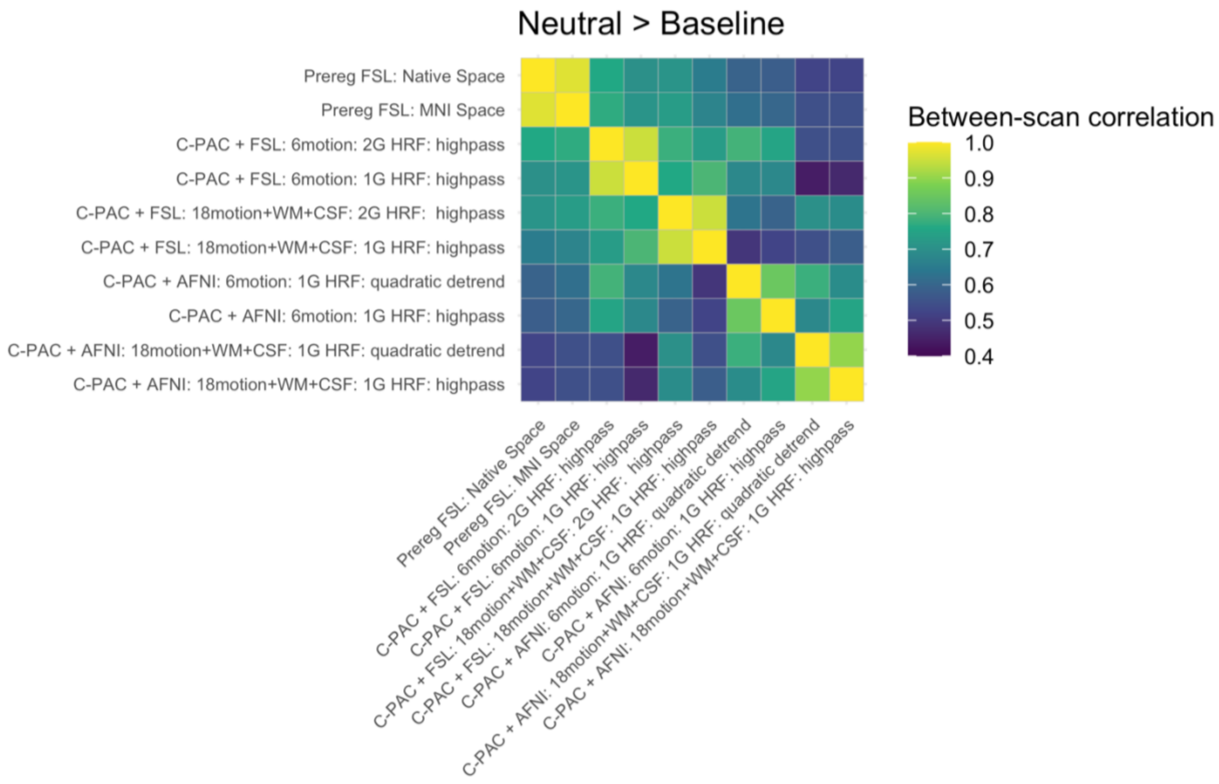


**sFigure 23:** Between-scan correlations for amygdala reactivity pipelines for neutral > baseline.

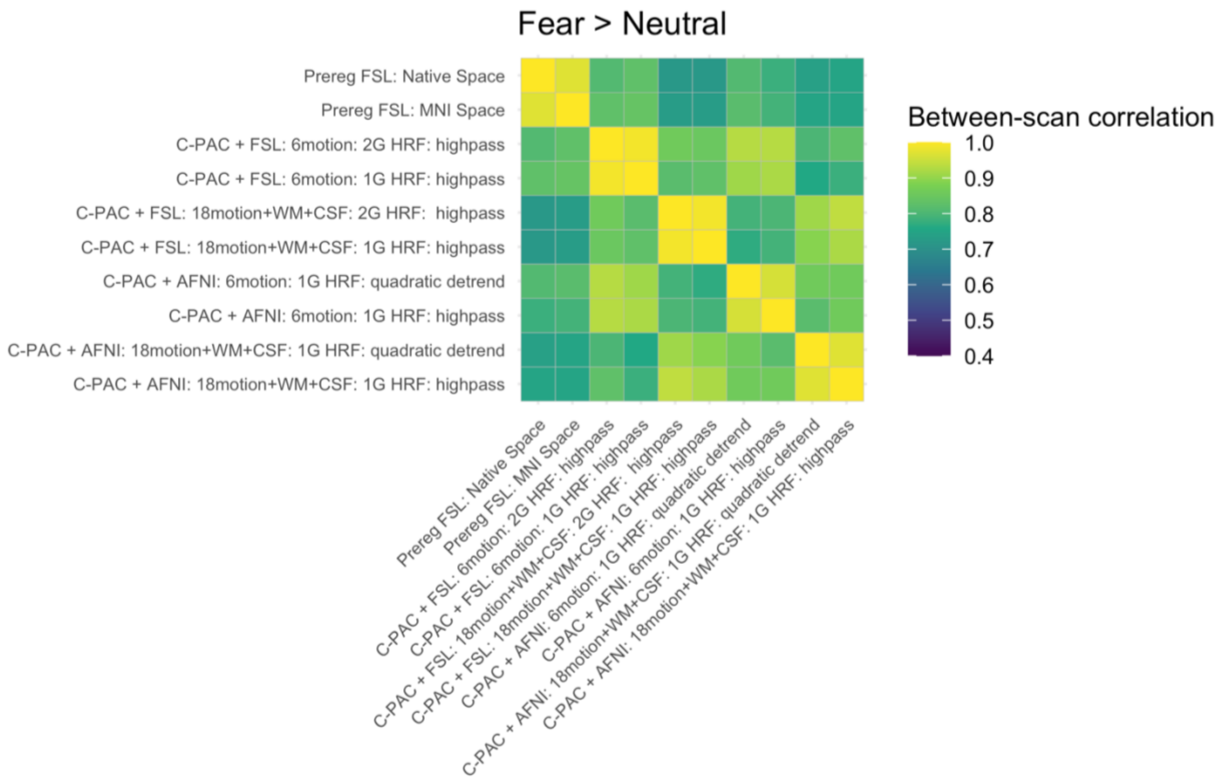


**sFigure 24:** Between scan correlations for amygdala reactivity pipelines for fear > neutral.

*Dependence of amygdala reactivity age-related change findings on previous work*

When we included only scans not previously used as a 'discovery set' to identify voxels changing with age in their reactivity to fear faces (a cluster in the right amygdala; Gee et al., 2013), estimates of age-related change were weaker on average, and the majority of posterior intervals for these estimates included 0. Permutation testing against equally sized samples, where 42 scans not previously studied were excluded from analysis at random (to form a 'null' distribution), indicated that for all pipelines tested, excluding the previously studied scans resulted in numerically weaker (less negative) age-related change (sFigure 25). Specifically, for the right amygdala (where age-related change was found in exploratory whole-brain analyses by Gee et al., 2013), age-related change when excluding these previously studied scans was less strong than the vast majority of permutation iterations excluding other scans at random for most pipelines. This indicates that analyses within the present study including these scans may somewhat overestimate age-related change due to partial dependence on the previous selection of the right amygdala by Gee et al. (2013). However, the magnitude of differences in findings between pipelines including versus excluding these previously-studied scans was small, and the vast majority of age-related change estimates are of the same sign regardless of exclusion of these scans. In addition, participants were younger on average by 1.5 years (95% PI [0.02, 3.00]) when studied at timepoint 1 by Gee et al. (2013) compared to other scans analyzed here due to the longitudinal study design. Thus, bias introduced by partially circular analyses here may not substantially alter conclusions across all specifications. Further, we also note that even analyses excluding these 42 participants cannot be considered entirely independent of the previous work, as the present work still examines follow-up scans from many of the same participants and uses the same stimuli.

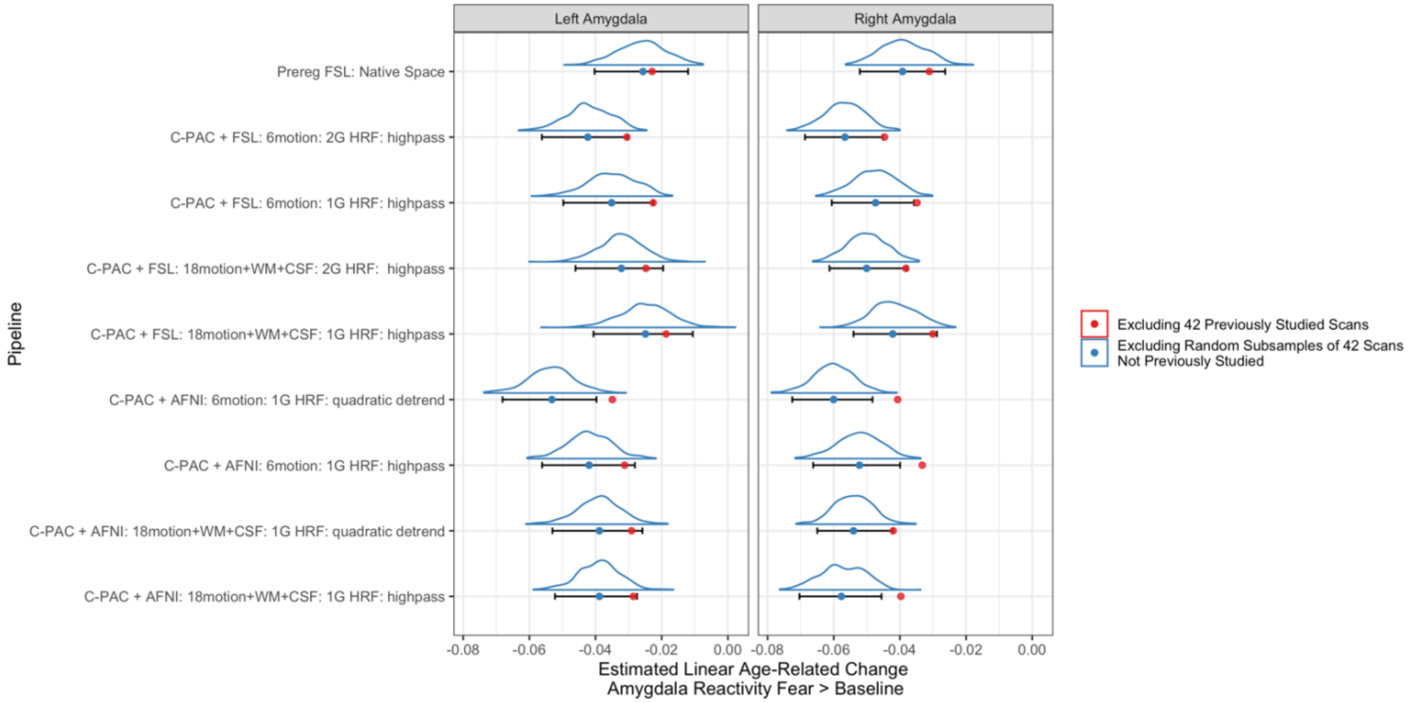

**sFigure 25:** Permutation tests for age-related change excluding previously studied scans
Blue distributions are ‘null’ distributions for age-related change for models of datasets excluding 42 randomly selected scans that had not been previously studied. Error bars indicate 95% confidence intervals based on these distributions, and blue points are the median value. Red points indicate estimate age-related change when excluding the 42 previously studied scans. Red points are always more positive than the blue points (especially for the right amygdala), indicating stronger median negative age-related change when these 42 scans are included than if excluded.

*Head Motion & Amygdala Reactivity*

We computed product-moment correlations between in-scanner head motion (mean FD) and amygdala reactivity for specifications across preprocessing pipelines and contrasts (see sFigure 26). Overall, few specifications resulted in amygdala reactivity estimates that were strongly correlated with head motion, although some estimates were significantly associated with motion (95% confidence interval excluding 0 for each contrast).

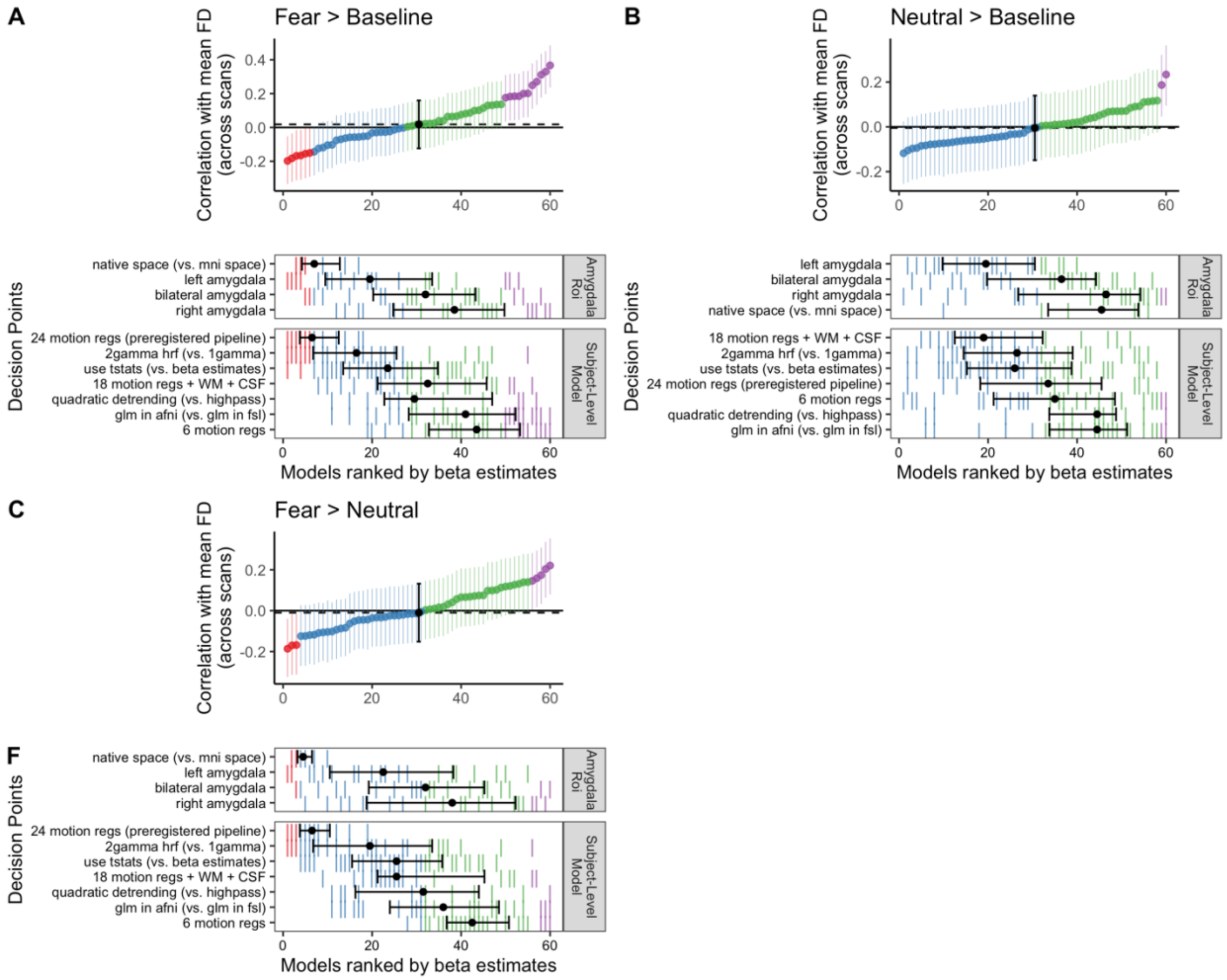


**sFigure 26:** Correlations between head motion and amygdala reactivity for each contrast
Plots show specification curves of correlations ranked by their value for the fear > baseline (A), neutral > baseline (B), and fear > neutral (C) contrasts. Color indicates sign of correlation estimates and whether respective 95% confidence intervals include 0 (red = negative excluding 0; blue = negative including 0, green = positive including 0, purple = positive excluding 0, black = median across all specifications).

Our preregistered exclusion criteria (<= 40 TRs with FD >= .9mm) based on in-scanner head motion was relatively lenient, especially compared to recent recommendations for resting-state fMRI preprocessing (Power et al., 2014). To examine whether results were driven by the inclusion of high-motion scans, we systematically varied an inclusion threshold for analysis from mean framewise displacement during the scan of 0.2-1.0mm in increments of 0.1mm. For each dataset based on the different inclusion thresholds, we modeled age-related change (longitudinal model #1) in bilateral amygdala (both Freesurfer-defined in native space and MNI space) reactivity using both t-statistics and beta estimates from the preregistered preprocessing pipeline. We did not observe meaningful differences in estimated age-related change as a function of head motion exclusion thresholds (see sFigure 27), indicating that age-related change findings in amygdala reactivity reported here are not likely driven purely by high-motion scans.

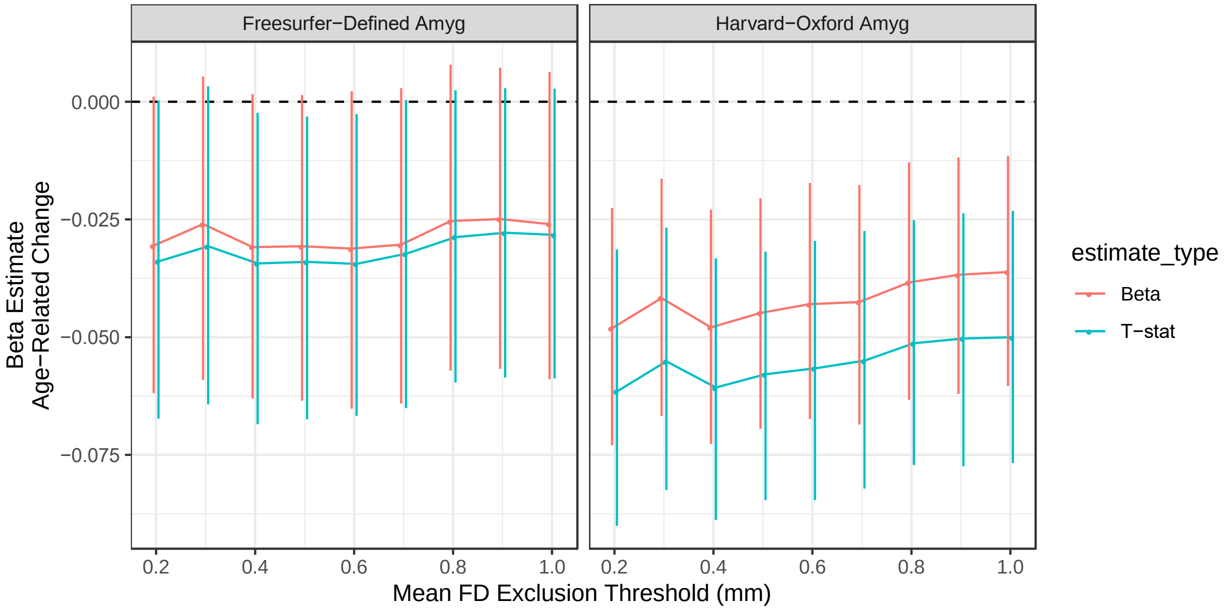


**sFigure 27:** Estimated age-related change as a function of mean FD exclusion threshold
The mean FD threshold for exclusion of a scan is shown on the x-axis, and the point estimate and 95% posterior intervals for corresponding age-related change estimates are on the y-axis.

*mPFC reactivity: supplemental results*

In addition to the amygdala, we also inspected reactivity in each of the 4 mPFC regions for the fear > baseline contrast with separate specification curves. To conserve computational resources, we fit these models using maximum likelihood with the lme4 R package (Bates et al., 2019), rather than fully Bayesian inference. Approximate 95% confidence intervals were constructed from these models by computing the interval ± 2 standard errors from the maximum likelihood estimate. Average reactivity differed by region, such that reactivity to fearful faces in the large vmPFC region was negative (i.e. signal lower than baseline), somewhat negative for mPFC region 3, and somewhat positive for mPFC regions 1-2 (sFigure 28A). Only in the large vmPFC region was average reactivity for fear > baseline reliably distinguishable from 0 across specifications. While age-related change estimates were rarely distinguishable from 0 for any region, estimates were all negative in sign for mPFC regions 1-3 (sFigure 28B-C).

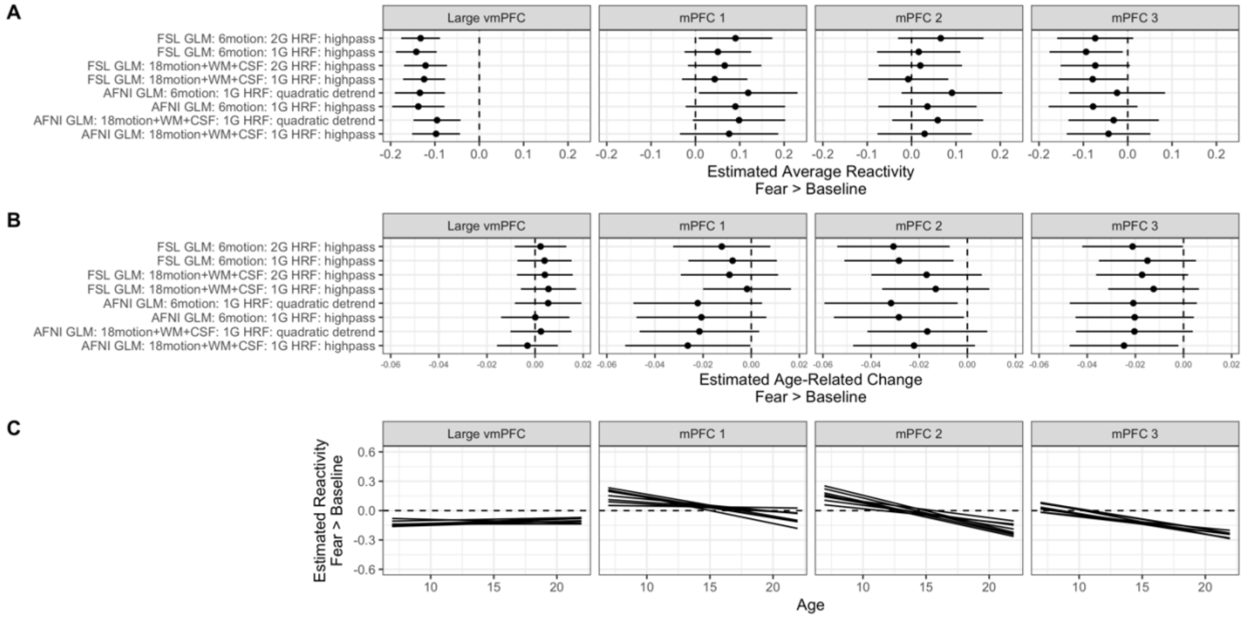


**sFigure 28:** Group average mPFC reactivity and age-related change for fear faces > baseline (A) estimated group mean mPFC reactivity for each preprocessing pipeline and ROI. (B) estimated age-related change in mPFC reactivity for each preprocessing pipeline and ROI. For A-B, error bars are approximate 95% confidence intervals. (C) model predictions for estimated fear > baseline mPFC reactivity as a function of age (in years), with different preprocessing pipelines plotted as individual spaghettis for each ROI.

Because Gee et al. (2013) reported that between-participant associations between amygdala and mPFC connectivity were positive among younger children and negative among older youth in a sample from the first timepoint studied here, we examined associations between amygdala and mPFC reactivity similarly in the current longitudinal sample. For these analyses, we also used multilevel linear regression models with the lme4 R package (without random slopes, but with random intercepts and a covariate for head motion). Across preprocessing pipelines, both the left and right amygdala, and all four mPFC regions, fear > baseline amygdala reactivity was positively associated with fear > baseline mPFC reactivity (see sFigure 29A-B). However, we did not find consistent evidence for age-related differences in associations between amygdala and mPFC reactivity for any mPFC region (sFigure 29C).

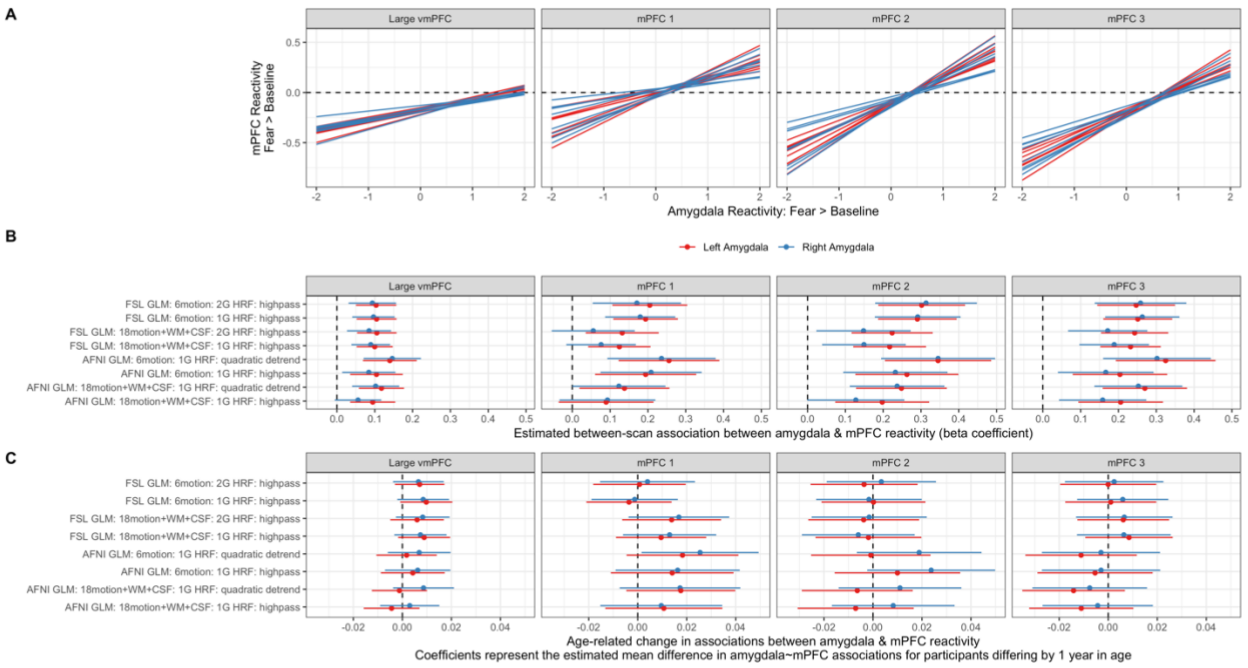


**sFigure 29:** Associations between amygdala & mPFC reactivity
(A) Model predictions for fear > baseline mPFC reactivity (y-axis) as a function of fear > baseline amygdala reactivity (x-axis). Individual spaghettis represent models for different preprocessing pipelines for both the left (red) and right (blue) amygdala. (B) Beta coefficients and 95% confidence intervals for associations between amygdala and mPFC reactivity. (C) Beta coefficients and 95% confidence intervals for age-related change in associations between amygdala and mPFC reactivity. Positive terms would represent stronger (more positive) amygdala—mPFC reactivity associations with increasing age.

### Within-scan changes in amygdala reactivity: supplemental results

*Group average within-scan changes in amygdala reactivity*

As with amygdala and mPFC reactivity, we modeled the group average within-scan change in amygdala reactivity using lme4. We estimated both the average slope of amygdala reactivity across trials (such that negative slope indicates linear decreases in amygdala reactivity across trials), and the mean difference between reactivity in trials 1-12 > 13-24 (first half > second half). We computed these group average estimates across bilateral, right, and left amygdala regions, with and without global signal subtraction, and for both the fear > baseline and neutral > baseline contrasts. Average slopes across trials were negative for both fear and neutral faces, although slopes were on average steeper for fear faces (sFigure 30). While on average, amygdala reactivity was higher for the first half of trials for fear faces across specifications, there were no consistent average differences between trial halves for amygdala reactivity to neutral faces (sFigure 31).

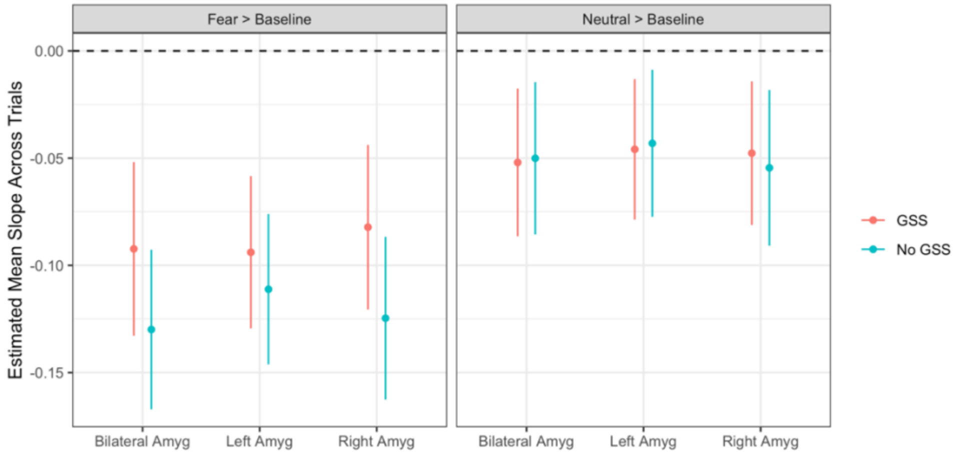


**sFigure 30:** Group average slopes in amygdala reactivity across trials
Negative slopes indicate linear decreases in amygdala reactivity across trials on average. Points display maximum likelihood estimates, and error bars are 95% confidence intervals.

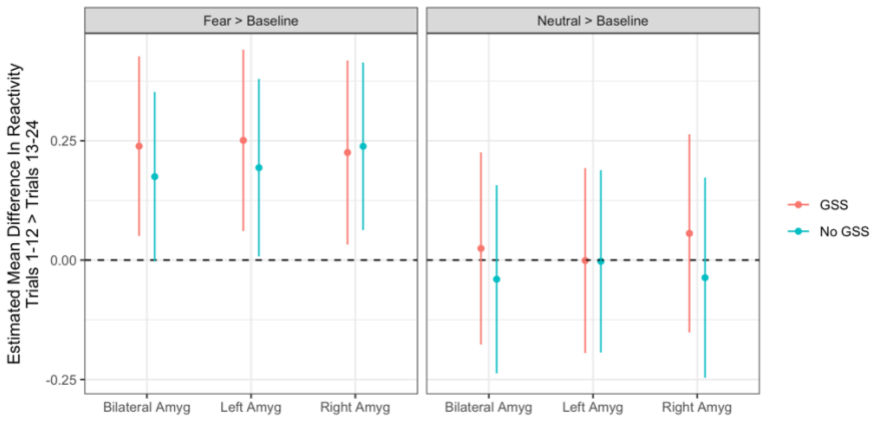


**sFigure 31:** Group average differences in amygdala reactivity across first > second half of trials Positive values indicate higher average amygdala reactivity in the first half of trials. Points display maximum likelihood estimates, and error bars are 95% confidence intervals.

*Age-related differences in within-scan amygdala reactivity change*

In addition to specification curves for amygdala reactivity slopes across trials (main manuscript Figure 3), we constructed parallel specification curves for age-related change in the difference in amygdala reactivity between trial halves. For the fear > baseline contrast, most specifications found evidence for an interaction between trial half and age (100% in the same direction, 66.7% of posterior intervals excluded 0), such that differences between amygdala reactivity in the first half > second half of trials were more negative (i.e. smaller positive differences, see sFigure 31) at older ages on average (sFigure 32A). Posterior intervals for this estimated interaction never excluded 0 for the left amygdala, but always did for the right or bilateral amygdala. For the neutral > baseline contrast, while the majority of specifications (83.3%) found numerically negative change, none of the 95% posterior intervals excluded 0 (sFigure 32B).

Single-trial models also indicated that for both fear and neutral faces, amygdala responses were larger for early trials for younger children, and more similar across age (though still positive) in later trials (see main manuscript Figure 3C, sFigure 33). Specification curves for single-trial models indicated that this pattern was somewhat stronger and more robust to analysis choices for fear faces (100% of 95% posterior intervals excluding 0), than for neutral faces (60% of posterior intervals excluding 0, see sFigure 34). Thus, analyses of slopes across trials, differences between trial halves, and single trial analyses indicated more consistent evidence of age-related change for fear faces than for neutral.

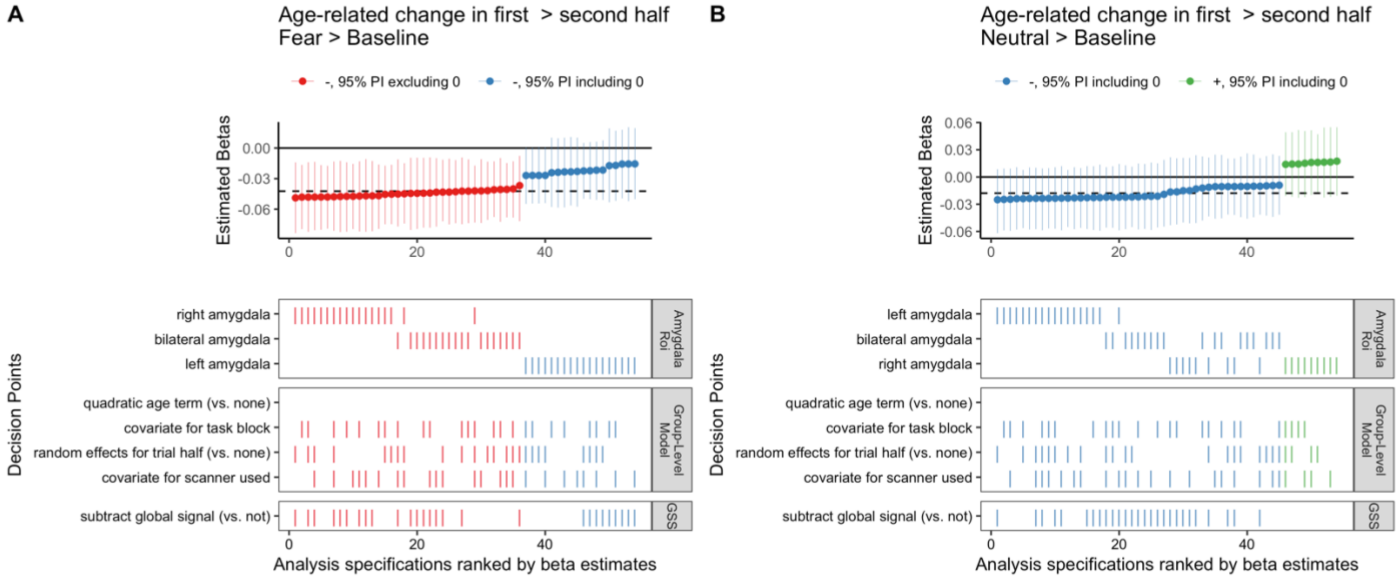


**sFigure 32:** Spec. curve for differences across task half in amygdala reactivity age-related change Specification curves showing parameter estimates and 95% posterior intervals for the estimated interaction term between age*trial half for amygdala reactivity, for the fear > baseline **(A)** and neutral > baseline **(B)** contrasts. Negative terms indicate that age-related change is more negative (stronger) during the first 12 trials compared to the last 12 (see main manuscript Figure 3B).

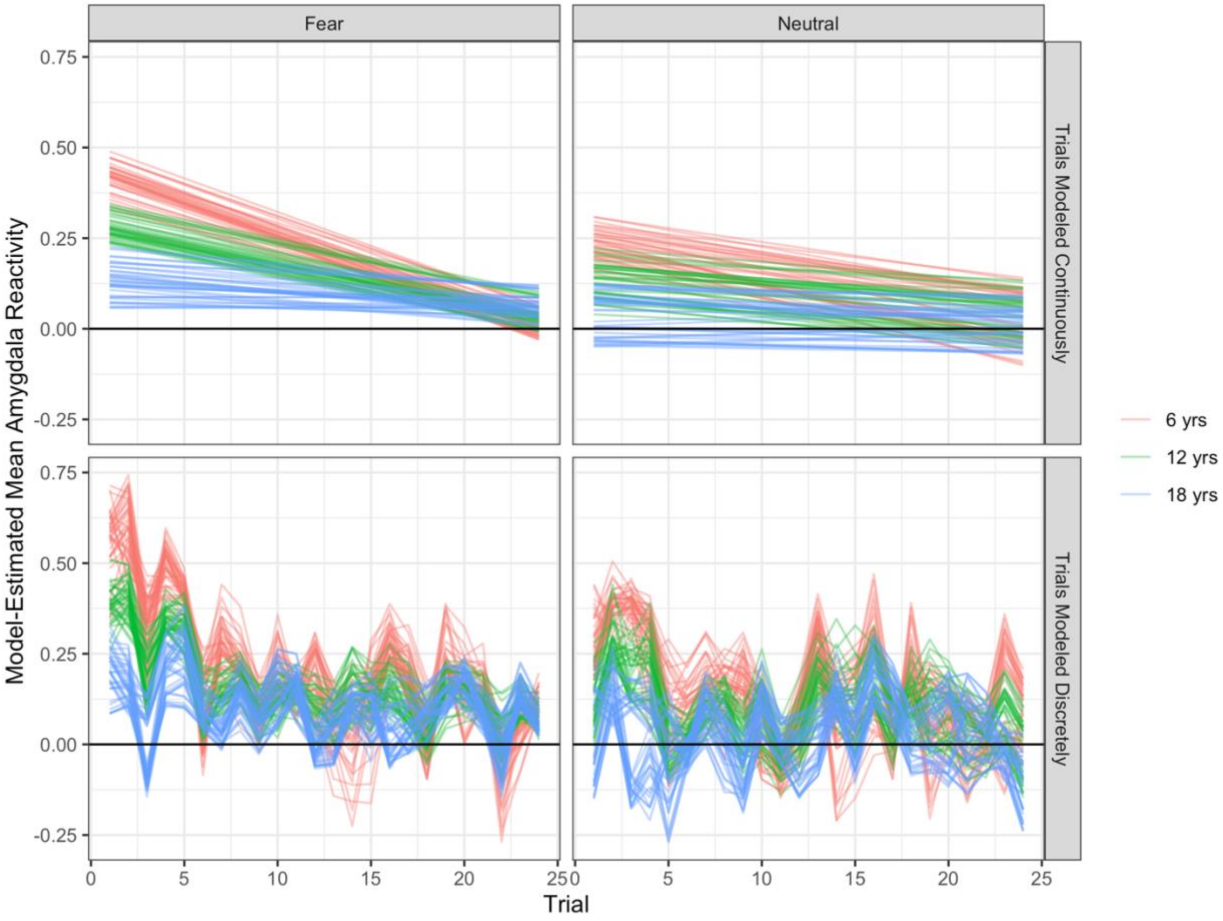


**sFigure 33:** Multiverse single-trial model predictions as a function of trial and age
Predictions for each single-trial model are plotted as individual 'spaghettis' for both fear trials (left) and neutral trials (right). For illustrative purposes, we plot predictions for an average person at age 6, 12, and 18 years of age. In the top panel, models include terms for linear trial associations, while in the bottom panel, trials are modeled discretely.

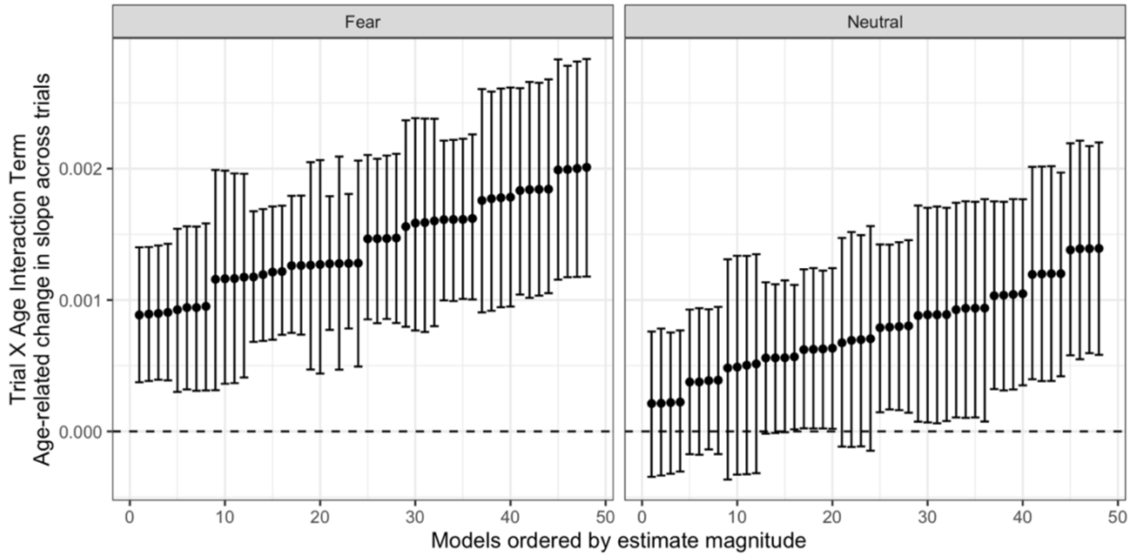


**sFigure 34:** Spec. curve of age*trial amygdala reactivity interactions from single-trial models Positive trial*age interaction terms indicate that slopes for within-scan linear changes in amygdala reactivity (as modeled through a single-trial multilevel model) were more positive (i.e. less negative, because slopes were negative on average) at older ages.

### gPPI Functional Connectivity Results

*Impacts of a deconvolution step on gPPI regressors and estimates*

gPPI regressors are interactions formed from the multiplication of the seed timeseries with the stimulus (task) regressors. In order for gPPI to measure 'task-dependent' connectivity, there must be adequate time when stimuli are 'on' versus 'off' such that such an interaction regressor can represent the difference in connectivity between two regions when the stimulus is present versus absent (McLaren et al., 2012). Unfortunately, within rapid event-related design, there were few periods of baseline 350ms presentations of either fear or neutral faces (ITI was jittered between 3-9s), other than the initial 20-second fixation period at the beginning of the scan. If it is then unclear which TRs represent connectivity when the stimuli are 'on' versus 'off', estimation of 'task-dependent' connectivity is difficult. Thus, other than the initial 20-second fixation period in our paradigm, resolution between stimulus presentations is low and there are very few moments in which the gPPI regressors are at baseline (i.e. flat, see sFigure 35 for an example set of regressors). This may especially be a problem for the gPPI regressor without deconvolution, where the stimulus regressor has already been convolved with the HRF before multiplication with the seed timeseries. Because of the slow temporal dynamics of the HRF, the stimulus regressor rarely returns to baseline between events (this would take 15-20s), and the gPPI regressor is then correlated with the seed timeseries.

Indeed, regressors both with and without deconvolution were collinear with the amygdala seed on average, although multicollinearity was more of a problem without deconvolution (sFigure 36). Thus, gPPI estimates without deconvolution might especially represent associations between brain regions that include “task-independent” signal in addition to connectivity associated particularly with the face stimuli. Within pipelines including a deconvolution step, however, tweaks to regularization methods (adding a lasso penalty) in the 3dTfitter algorithm or up-sampling of the seed timeseries to 0.1s resolution before deconvolution resulted in substantially differing gPPI regressors. As shown in sFigure 36 (top panel), gPPI regressors for the same scan using different deconvolution methods were only often only weakly associated ($r < .5$). Such differences among regressors all ostensibly using deconvolution indicates that our solutions for the ‘underlying neuronal timecourse’ (and the resulting gPPI estimates) may be unreliable due to their sensitivity to such changes in the deconvolution pipeline.

We then asked how similar amygdala gPPI results would be at a voxelwise level when using a pipeline with versus without deconvolution. We compared voxelwise patterns between t-statistic maps for fear > baseline, fear > neutral, and neutral > baseline gPPI with versus without deconvolution for the same scans. Patterns were overall positively correlated across for all mPFC regions (as well as whole-brain patterns), but varied significantly such that the median correlation between patterns with versus without deconvolution was never above $r = .5$ for the fear > baseline or neutral > baseline contrasts (sFigure 37). While patterns were slightly more similar for the fear > neutral contrast, that such correlations were only moderate indicated substantial differences in patterns of results across voxels between pipelines with versus without deconvolution. The higher variability in similarity values across mPFC regions 1-3 is likely because these ROIs were much smaller, and thus correlations were calculated across far fewer voxels. These substantial differences when comparing amygdala gPPI results with versus without deconvolution for the same scan likely play a major role in the discrepancies in findings for age-related change in amygdala—mPFC gPPI. Although we did not test other task paradigms here, we speculate that gPPI analyses with block designs or event-related designs with longer ITIs (20+ seconds) may be more successful at estimating task-dependent connectivity.

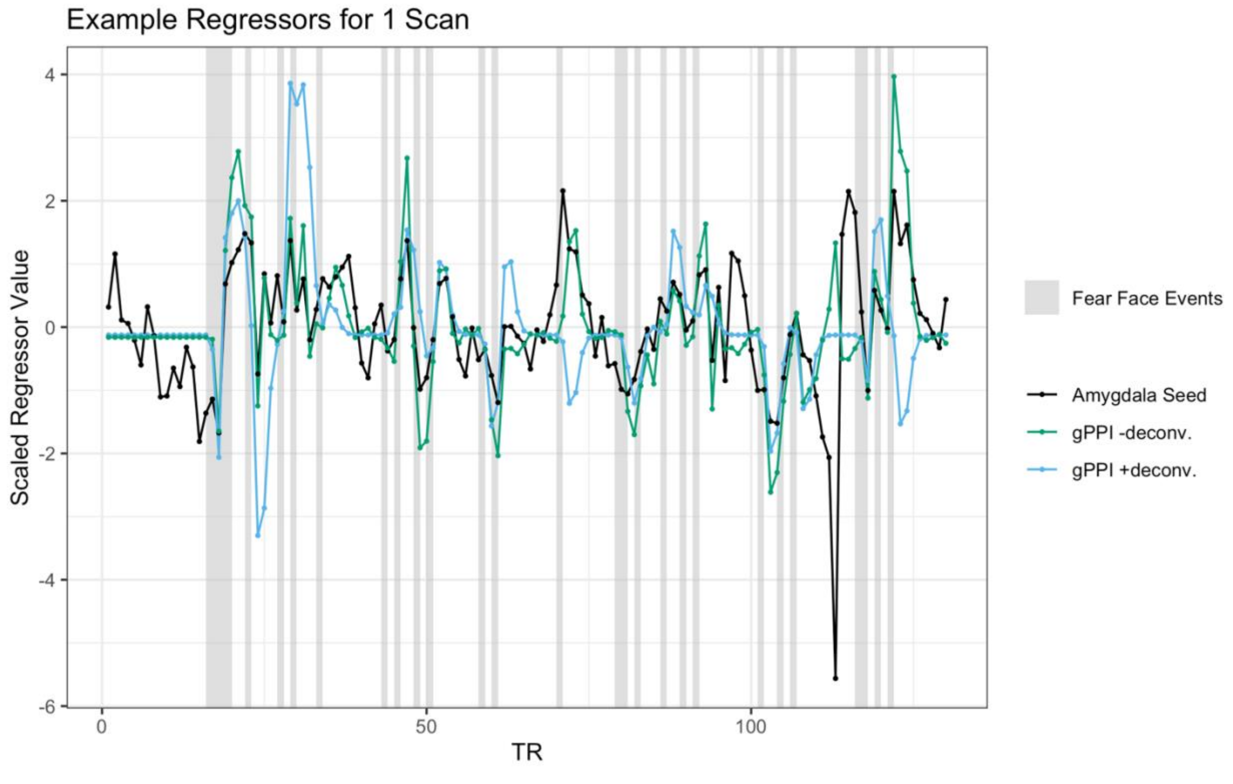


**sFigure 35:** Example gPPI and amygdala seed regressors for one scan
Timecourses for the average timeseries of the amygdala seed (black), gPPI regressor without deconvolution (green), and gPPI regressor with deconvolution (blue) for an example participant. TRs where fear face stimuli are presented are highlighted in grey. gPPI regressors, especially

the regressor without deconvolution, are rarely flat other than at the beginning of the task, because there are few temporal gaps between fear face stimuli of more than 10-15s. Thus, both gPPI regressors, but especially the one without deconvolution, tend to be correlated with the seed timecourse.

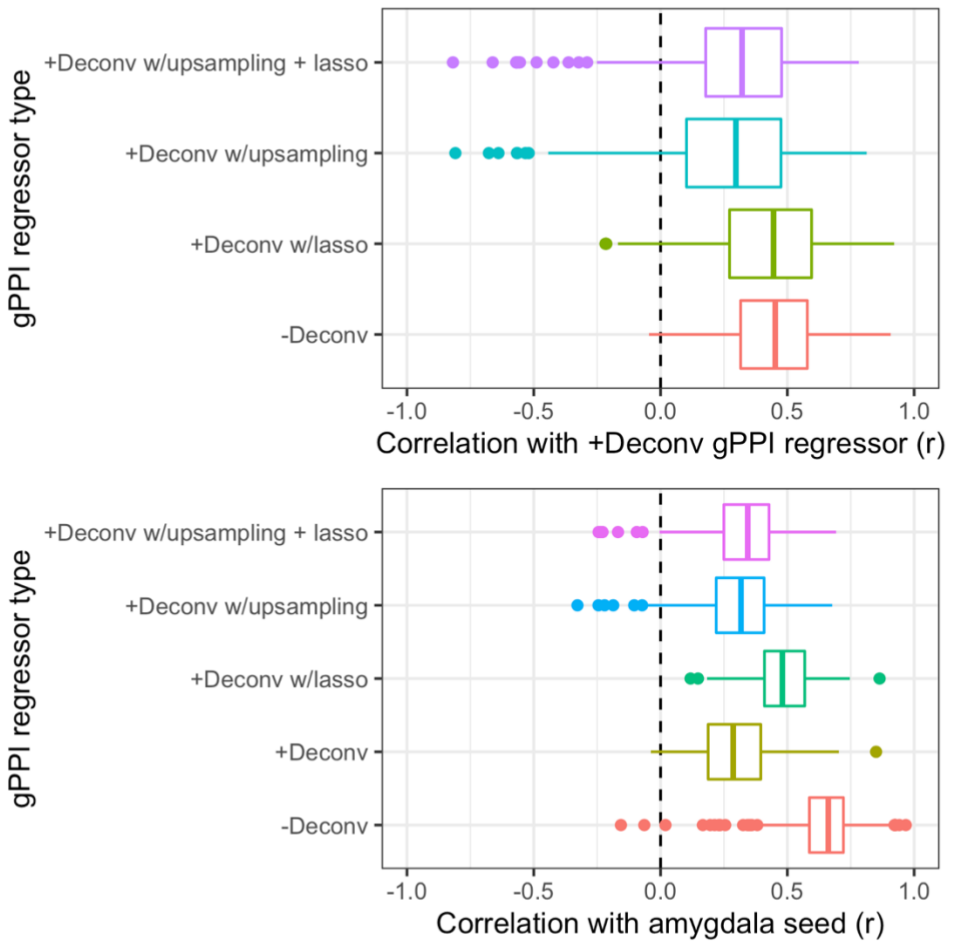


**sFigure 36:** Correlations between gPPI regressors, and between gPPI regressors and the seed **Top:** boxplots show distributions of correlations across between different versions of gPPI regressors for the same scan for all scans. +Deconv represents the deconvolved regressor without up-sampling or lasso regularization, which was used for pipelines in the main manuscript. These +Deconv regressors were about equally similar to -Deconv regressors (regressors used in the main manuscript without deconvolution) as they were to +Deconv regressors with lasso regularization. +Deconv regressors were even less similar to +Deconv regressors with up-sampling, or up-sampling and lasso regularization. **Bottom:** boxplots show distributions of correlations between different versions of gPPI regressors with the amygdala seed timeseries across all scans.

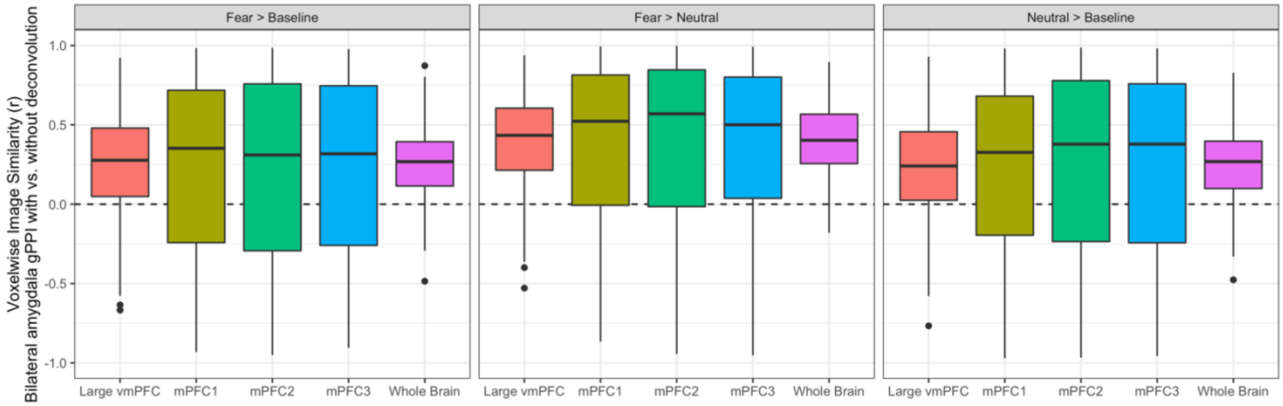


**sFigure 37:** Voxelwise image similarities of gPPI estimates with vs. without deconvolution Values on the y-axis represent product-moment correlations between t-statistic maps for each ROI with versus without deconvolution.

*Group mean amydala-mPFC gPPI estimates*

To preserve computational resources, all models for mean amygdala—mPFC gPPI were fit using the lme4 R package with intercepts allowed to vary by participant. Overall, we did not see consistent evidence for group mean task-dependent amygdala—mPFC connectivity distinct from 0 for any contrast or ROI (see sFigure 38). As discussed previously, however, the lack of detected group average task-dependent connectivity may owe more to the fact that the task paradigm was ill-suited for estimation of gPPI than absence of true connectivity.

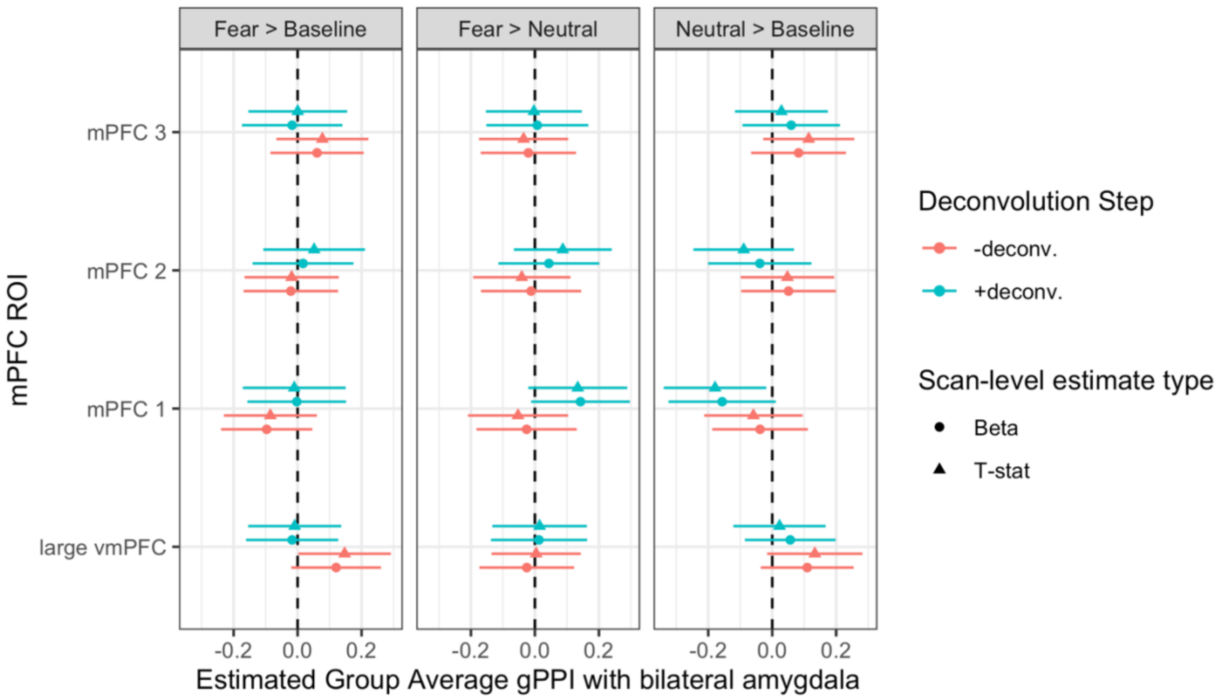


**sFigure 38:** Group average amygdala—mPFC gPPI estimates for each contrast and ROI The x-axis represents estimated average task-dependent amygdala—mPFC connectivity for each ROI (on the y-axis) and contrast (left = fear > baseline, middle = fear > neutral, right = neutral > baseline). Pipelines with deconvolution are represented in blue, and without deconvolution in red.

*Multiverse analyses of age-related change in amygdala—mPFC gPPI*

In addition to the constructing specification curves for age-related change in amygdala—mPFC gPPI for the fear > baseline contrast as reported in the main manuscript (see Figure 4), we constructed parallel specification curves for the neutral > baseline and fear > neutral contrasts. For the neutral > baseline contrast the vast majority of pipelines (98.2%) found positive age-related change, though only 29.5% of pipelines estimated such change with a posterior interval excluding 0 (sFigure 39). Similar to the fear > baseline contrast, pipelines with a deconvolution step tended to find less positive age-related change. For the fear > neutral contrast, 86.1% of pipelines estimated numerically negative age-related change, though only 2.1% of pipelines did so with a posterior interval excluding 0 (sFigure 40). Thus, across all contrasts there was no consistent evidence of age-related change in amygdala—mPFC gPPI.

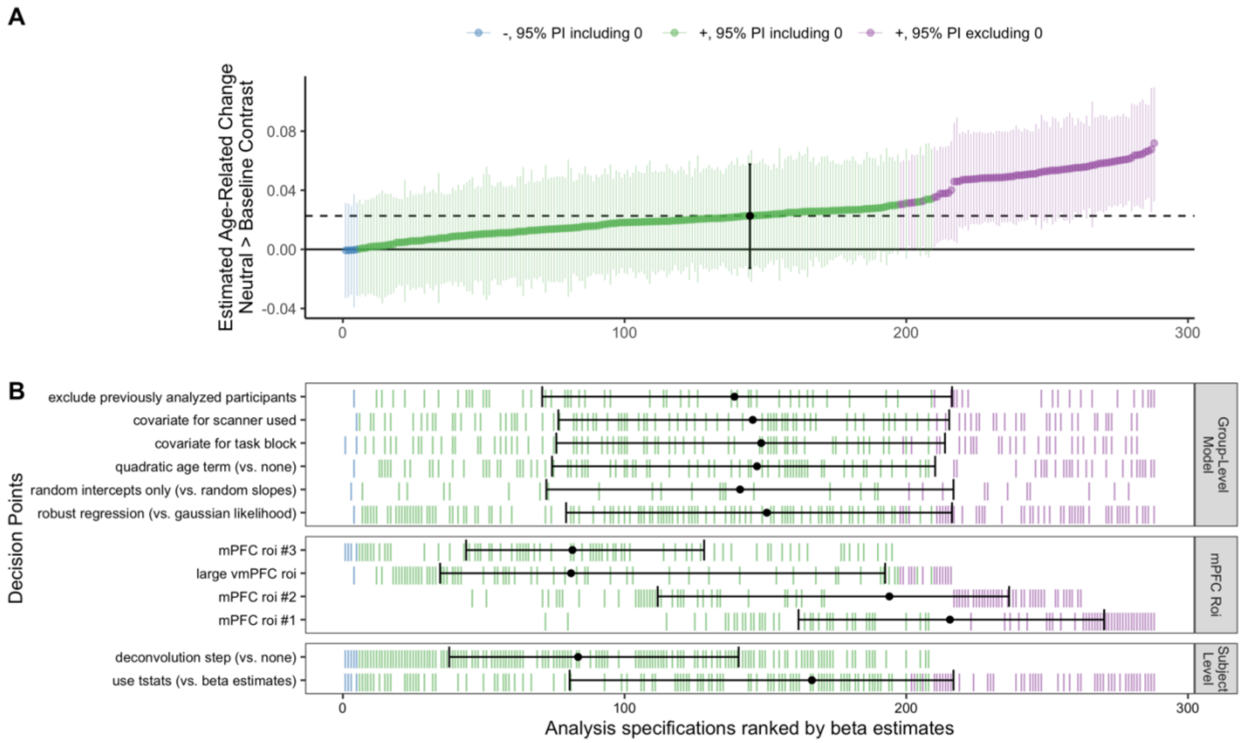


**sFigure 39:** Spec. curve for age related change in amygdala—mPFC gPPI for neutral > baseline
**A:** Points represent estimated linear age-related change in amygdala—mPFC gPPI for each specification, and lines represent corresponding 95% posterior intervals. **B:** Variables on the y-axis represent analysis choices, corresponding lines indicate that a choice was made, and blank space indicates that the choice was not made in a given analysis.

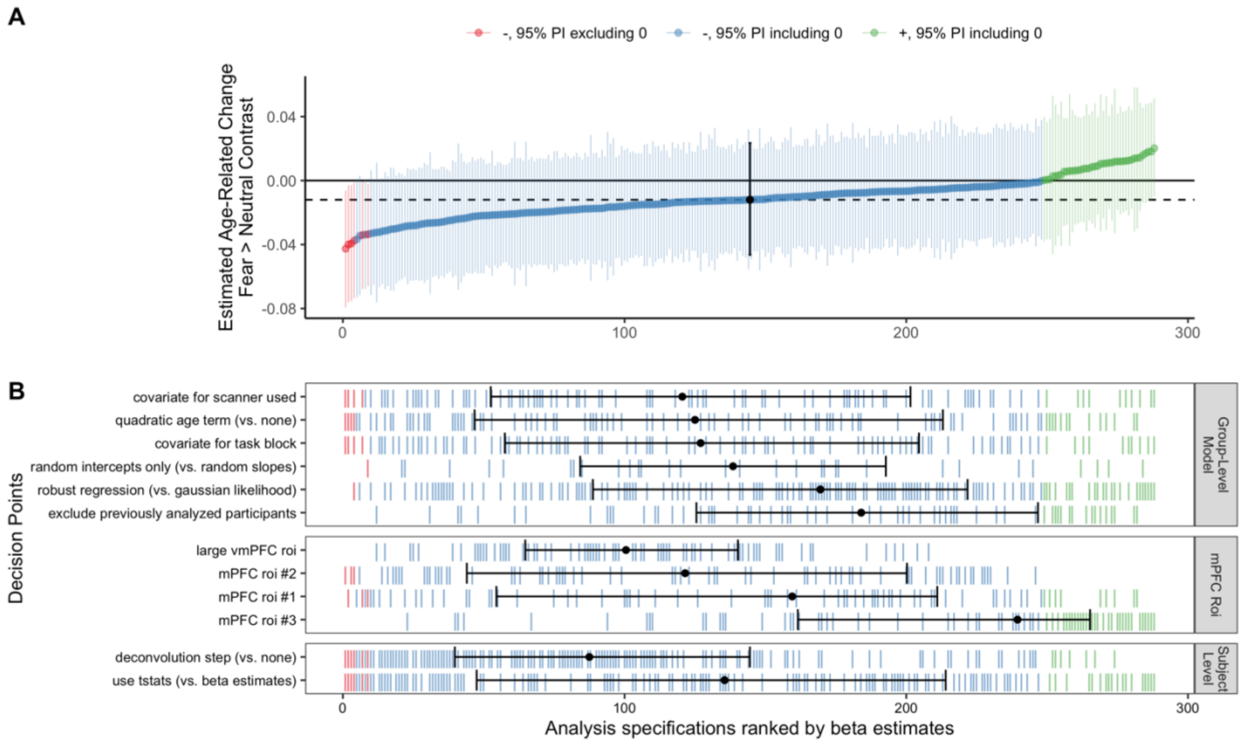


**sFigure 40:** Spec. curve for age related change in amygdala—mPFC gPPI for fear > neutral
**A:** Points represent estimated linear age-related change in amygdala—mPFC gPPI for each specification, and lines represent corresponding 95% posterior intervals. **B:** Variables on the y-axis represent analysis choices, corresponding lines indicate that a choice was made, and blank space indicates that the choice was not made in a given analysis.

*Impacts of analysis choices on age-related change estimates for amygdala—mPFC gPPI*

As with amygdala reactivity, we explored the impacts of gPPI analysis choices on estimates of linear age-related change, conditional on all other decision points. For the fear > baseline contrast, whether to use a deconvolution step or not made by far the biggest impact on estimated age-related change (see sFigure 41). Deconvolution also impacted estimates for the neutral > baseline contrast (sFigure 42) and fear > neutral contrast (sFigure 43).  The chosen mPFC ROI also had a large impact on estimated age-related change.

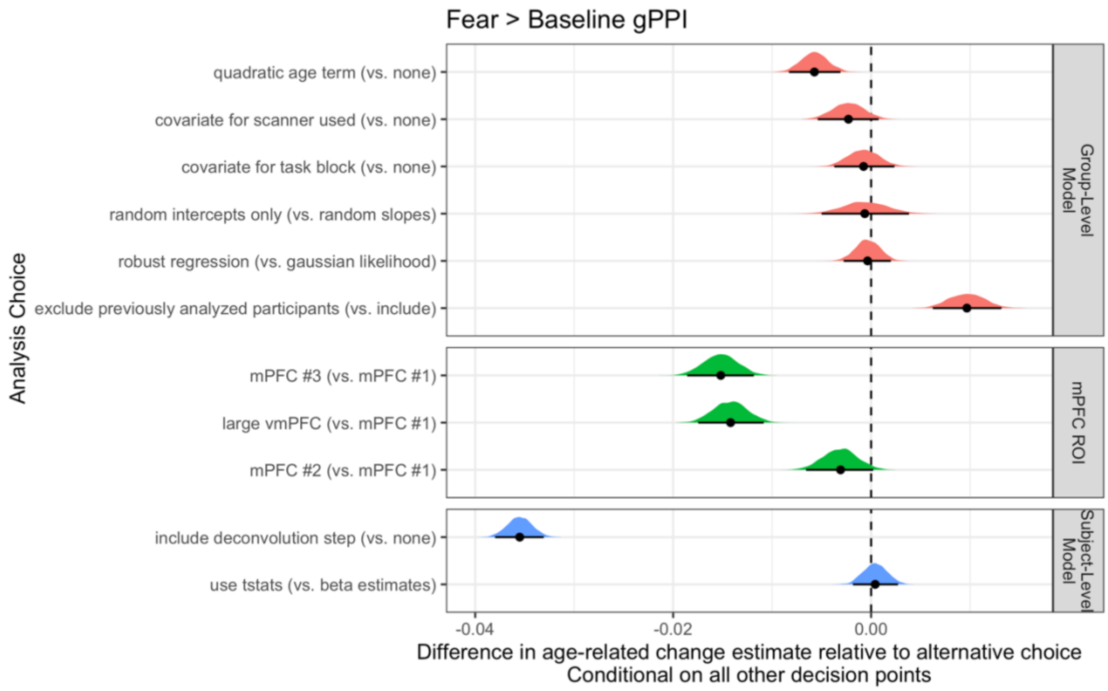


**sFigure 41.** Fork impacts on age-related change for fear > baseline amygdala—mPFC gPPI Posterior distributions and 95% posterior intervals are shown, representing the average difference in linear age-related change estimates relative to the alternative choice.

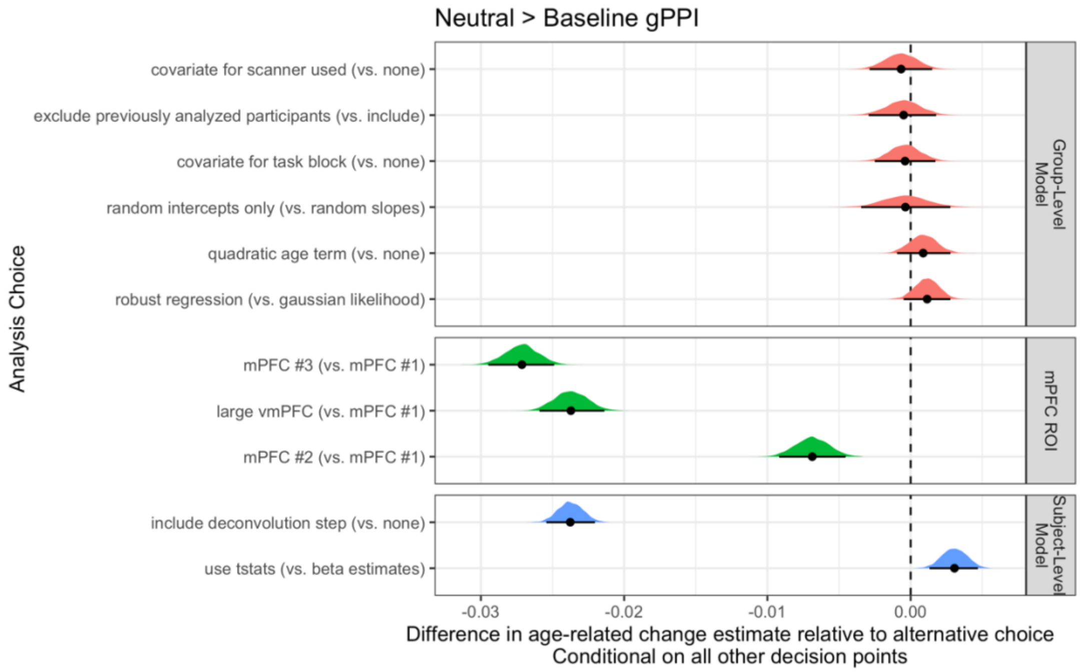

**sFigure 42:** Fork impacts on age-related change for neutral > baseline amygdala—mPFC gPPI Posterior distributions and 95% posterior intervals are shown, representing the average difference in linear age-related change estimates relative to the alternative choice.

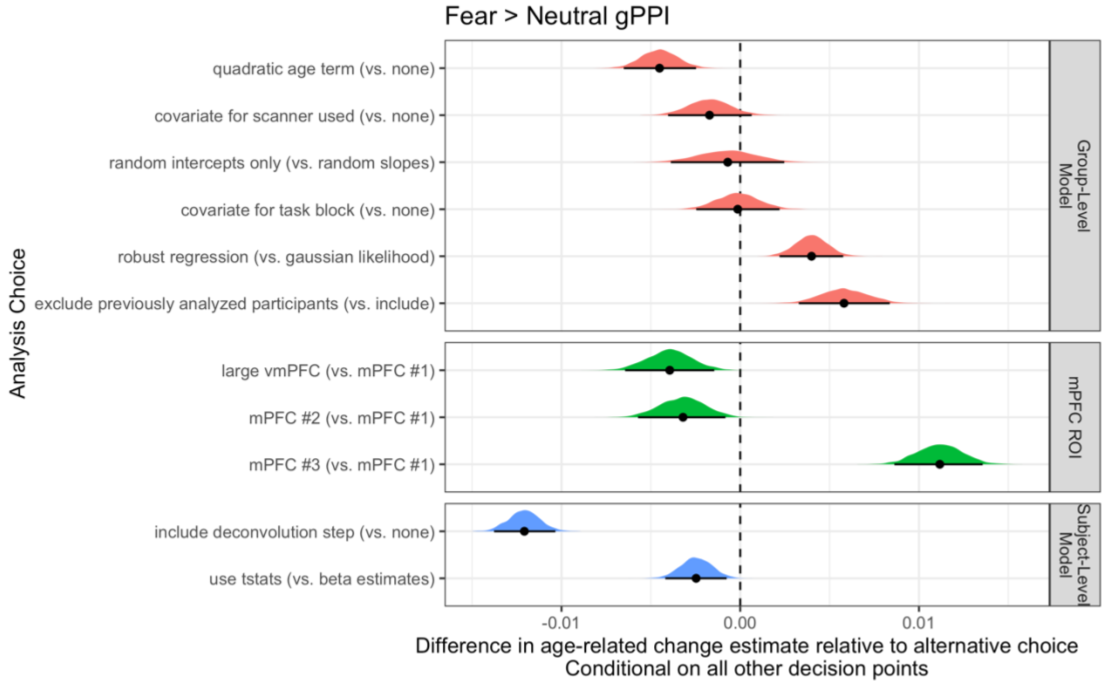


**sFigure 43:** Fork impacts on age-related change for fear > neutral amygdala—mPFC gPPI Posterior distributions and 95% posterior intervals are shown, representing the average difference in linear age-related change estimates relative to the alternative choice.

*Impacts of regressor centering on age-related change in amygdala—mPFC gPPI*

As previous work has recommended centering the task regressor in gPPI models using deconvolution (Di et al., 2017), we investigated whether age-related change gPPI results with deconvolution differed as a function of this centering. Overall, regressor centering had little impact on linear age-related change estimates, with deconvolution and choice of mPFC ROI having relatively more influence on regression estimates (sFigure 44A). Further, scan-level estimates for the fear > baseline contrast were highly similar between pipelines where the task regressor was centered before creating the gPPI regressor and pipelines where no such centering was done (sFigure 44B). The similarity of scan-level estimates indicated that this centering step had little impact on individual gPPI estimates in this instance.

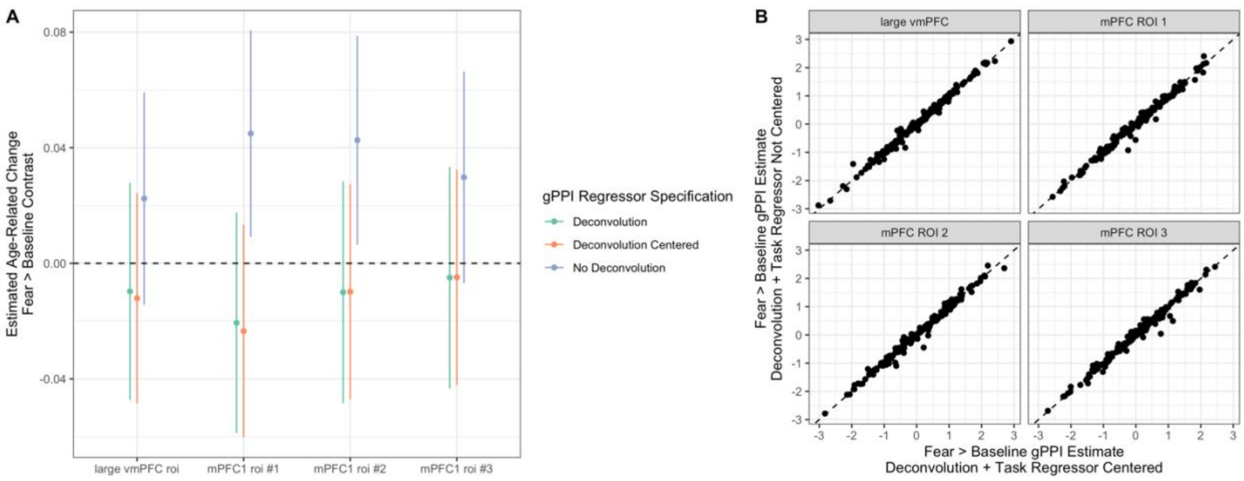


**sFigure 44:** Impacts of centering the task regressor in gPPI models with deconvolution
**A:** Posterior distributions and 95% posterior intervals are shown for age-related change estimates in amygdala—mPFC gPPI functional connectivity as a function of the gPPI regressor specification. Models with deconvolution and without centering of the task regressor (green) demonstrated highly similar estimated age-related change to models with deconvolution and centering the task regressor (orange). Models without deconvolution (purple), by comparison, showed differing estimates of age-related change. Models displayed are for the fear > baseline contrast. **B:** Direct comparison of scan-level estimates for fear > baseline amygdala—mPFC gPPI with deconvolution and with the task regressor centered (x-axis) versus not centered (y-axis). Panels separate gPPI estimates by mPFC ROI.

*Non-linear age-related changes in amygdala—mPFC gPPI*

We constructed specification curves for quadratic age-related change parameters across all models including quadratic terms for the fear > baseline (sFigure 45), neutral > baseline (sFigure 46), and fear > neutral (sFigure 47) amygdala—mPFC gPPI. Across all contrasts, few specifications (20.8% for fear > baseline, 1.0% for neutral > baseline, 8.3% for fear > neutral) estimated quadratic terms distinguishable from 0. Quadratic fits also varied considerably in sign for each contrast, such that there was not consensus on ‘U-shaped’ or ‘inverse U-shaped’ change. Thus, while the current study may not have been adequately powered to estimate quadratic age-related change, we did not find consistent evidence for either peaks or troughs in amygdala—mPFC gPPI.

We also constructed inverse age models for age-related change in amygdala—mPFC gPPI (sFigure 48). As with linear models, such models indicated that a deconvolution step largely influenced age-related change estimates for the fear > baseline contrast, often flipping the sign from positive change (without deconvolution) to negative change (with deconvolution, sFigure 48). However, deconvolution had a relatively smaller impact for the neutral > baseline and fear > neutral contrasts.

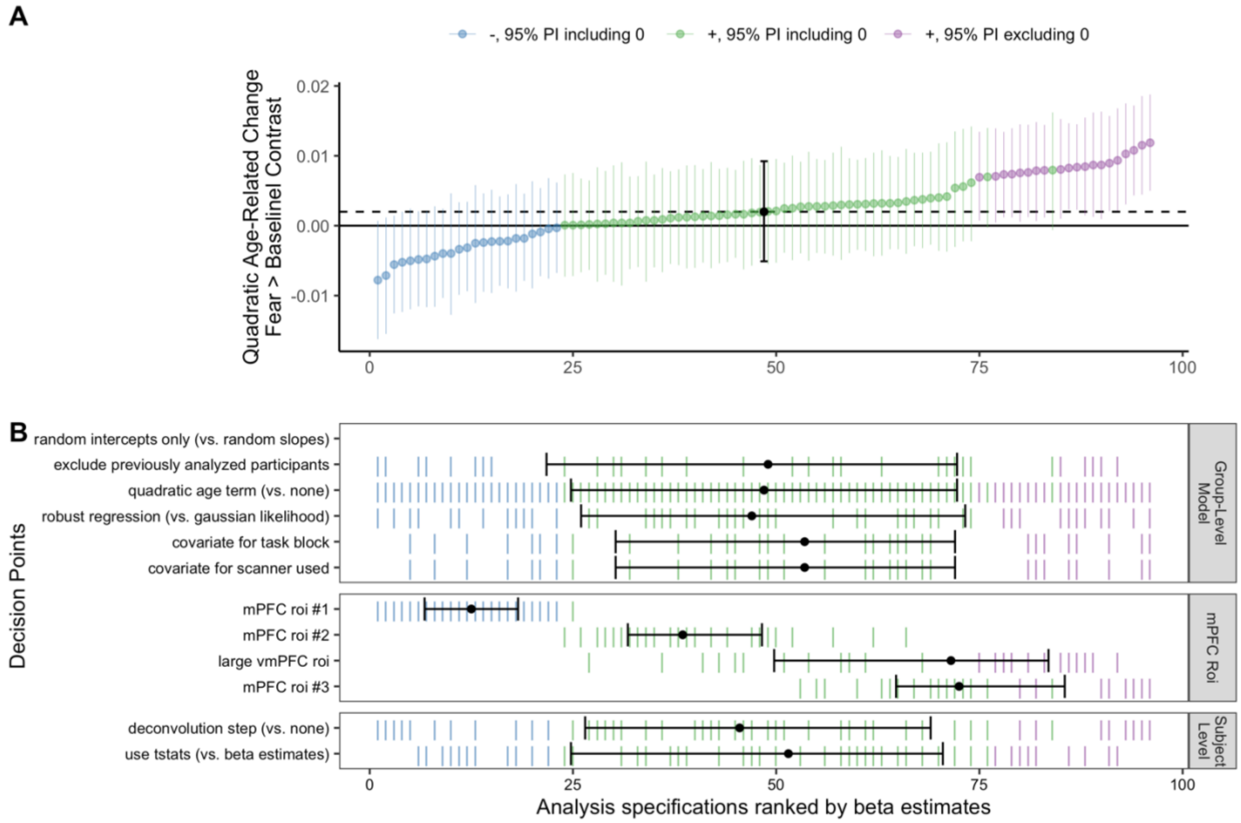

**sFigure 45:** Spec. curve for quadratic age-related changes in fear > baseline amygdala—mPFC gPPI
**A:** Points represent estimated quadratic age-related change in amygdala—mPFC gPPI for each specification, and lines represent corresponding 95% posterior intervals. **B:** Variables on the y-axis represent analysis choices, corresponding lines indicate that a choice was made, and blank space indicates that the choice was not made in a given analysis.

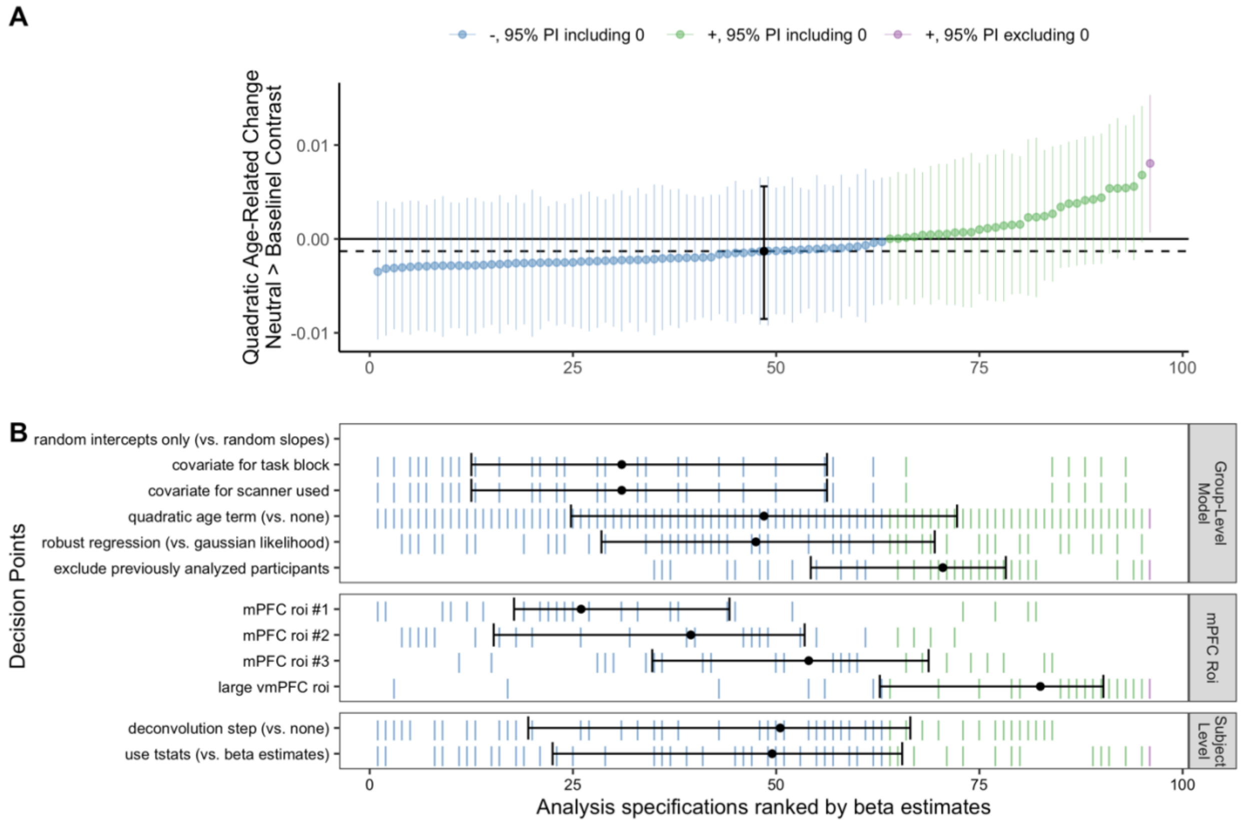


**sFigure 46:** Spec. curve for quadratic age-related changes in neutral > baseline amygdala—mPFC gPPI
**A:** Points represent estimated quadratic age-related change in amygdala—mPFC gPPI for each specification, and lines represent corresponding 95% posterior intervals. **B:** Variables on the y-axis represent analysis choices, corresponding lines indicate that a choice was made, and blank space indicates that the choice was not made in a given analysis.

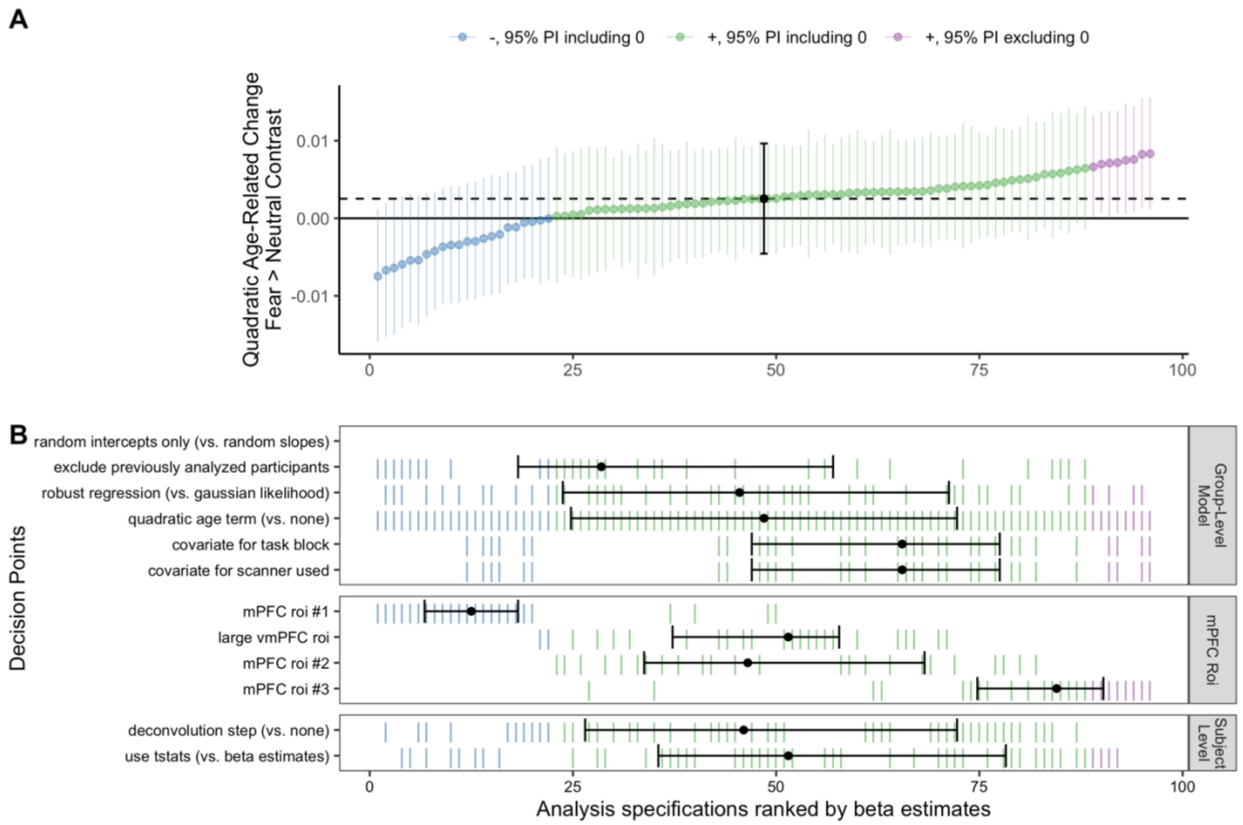


**sFigure 47:** Spec. curve for quadratic age-related changes in fear > neutral amygdala—mPFC gPPI **A:** Points represent estimated quadratic age-related change in amygdala—mPFC gPPI for each specification, and lines represent corresponding 95% posterior intervals. **B:** Variables on the y-axis represent analysis choices, corresponding lines indicate that a choice was made, and blank space indicates that the choice was not made in a given analysis.

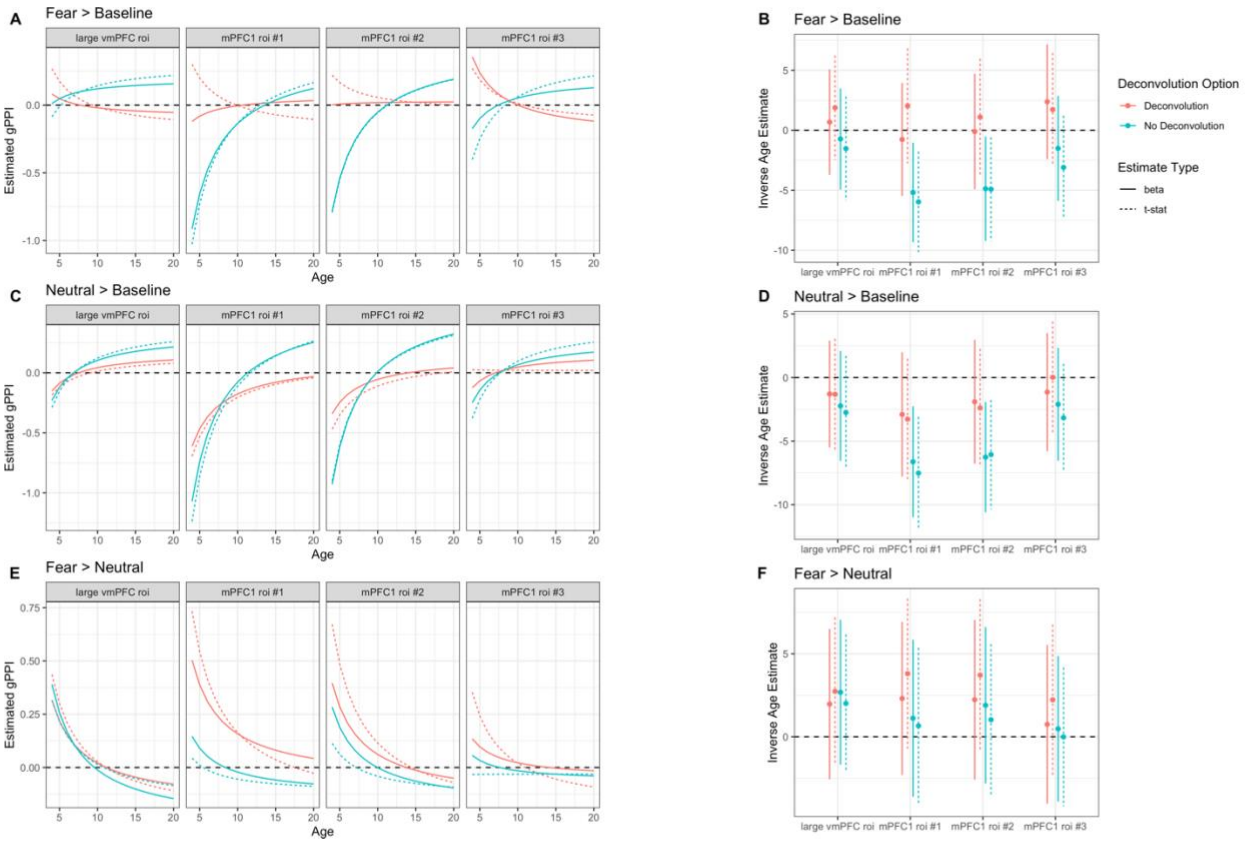


**sFigure 48:** Inverse age models for amygdala—mPFC gPPI
Left panels show fitted model predictions for inverse age models for the fear > baseline (**A, top**), neutral > baseline (**C, middle**), and fear > neutral (**E, bottom**) contrasts. Specifications including a deconvolution option are plotted in red, and without deconvolution in blue. Specifications using beta estimates are plotted with filled lines and with t-stats using dotted lines. Right panels show beta estimates for corresponding models for each contrast. Positive estimates for inverse age indicate decreases in amygdala—mPFC gPPI as a function of age, and vice-versa.

*Correlations between head motion and amygdala—mPFC gPPI estimates*

We computed product-moment correlations between in-scanner head motion (mean FD) and amygdala—mPFC gPPI across pipelines (deconvolution versus none), contrasts, and ROIs (see sFigure 49). Overall, head motion was not strongly correlated with gPPI estimates, such that few 95% confidence intervals for correlations excluded 0.

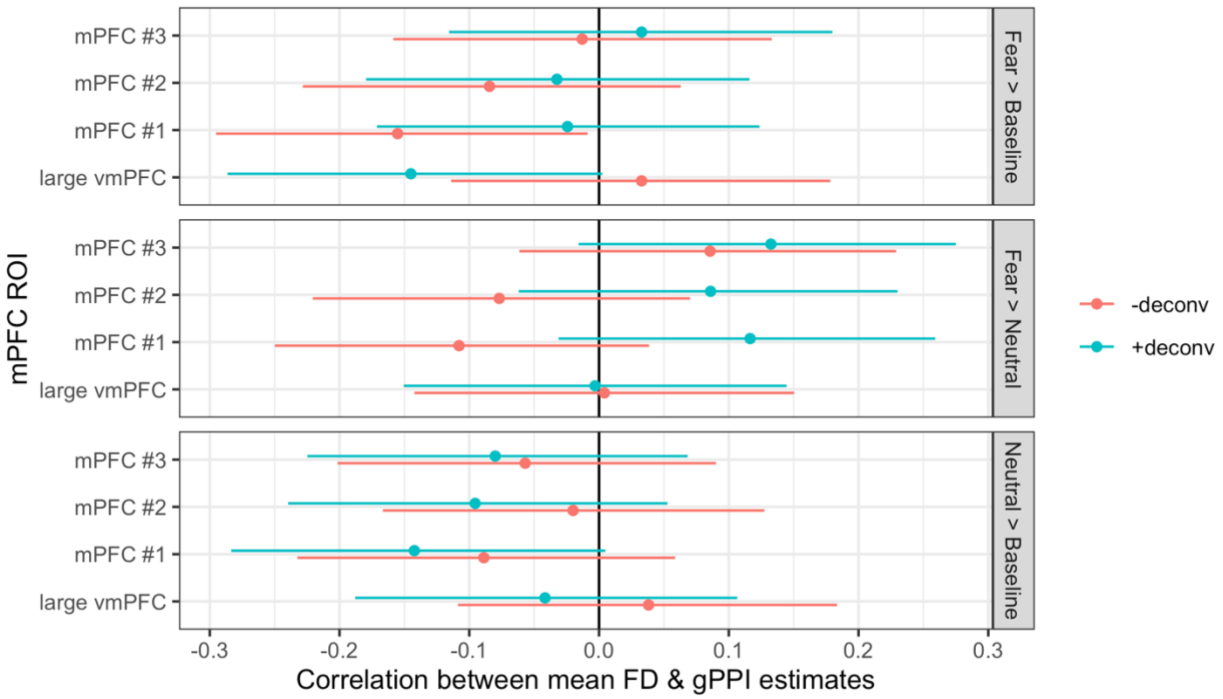


**sFigure 49:** Correlations between mean FD & gPPI estimates across scans
For each contrast, pipeline (+deconv. versus -deconv), and ROI, points show estimated product-moment correlations and error bars represent 95% confidence intervals.

*Task-independent amygdala—mPFC connectivity estimates from gPPI models*

Within the gPPI model, the association between the seed timeseries (or 'physiological' term) and target voxel has been conceptualized as representing 'task-independent' functional connectivity (Greene et al., 2020). Although we cannot be sure that such measurements are truly 'task-independent' without analysis of tasks beyond the current study, we used these estimates to explore amygda-mPFC functional connectivity while controlling for task-induced variance. For all mPFC ROIs, such task-independent connectivity with the amygdala was positive on average across all participants (sFigure 50). The positive task-independent amygdala—mPFC connectivity found here may be an overestimate, however, as gPPI pipelines did not include a global signal correction (Power et al., 2017).

In addition, pipelines with deconvolution found age-related increases in task-independent amygdala—mPFC connectivity, while pipelines without deconvolution found age-related decreases (sFigure 51B). Few 95% confidence intervals excluded 0 for age-related change for either set of pipelines however (sFigure 51A).

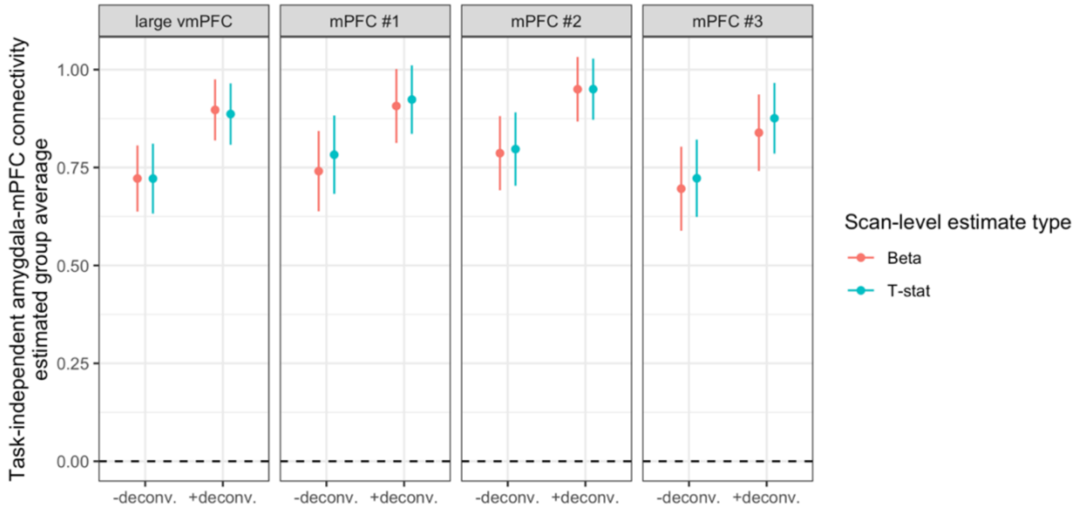


**sFigure 50:** Group average task-independent amygdala—mPFC connectivity
Points show estimates and error bars represent 95% confidence intervals for each pipeline and ROI.

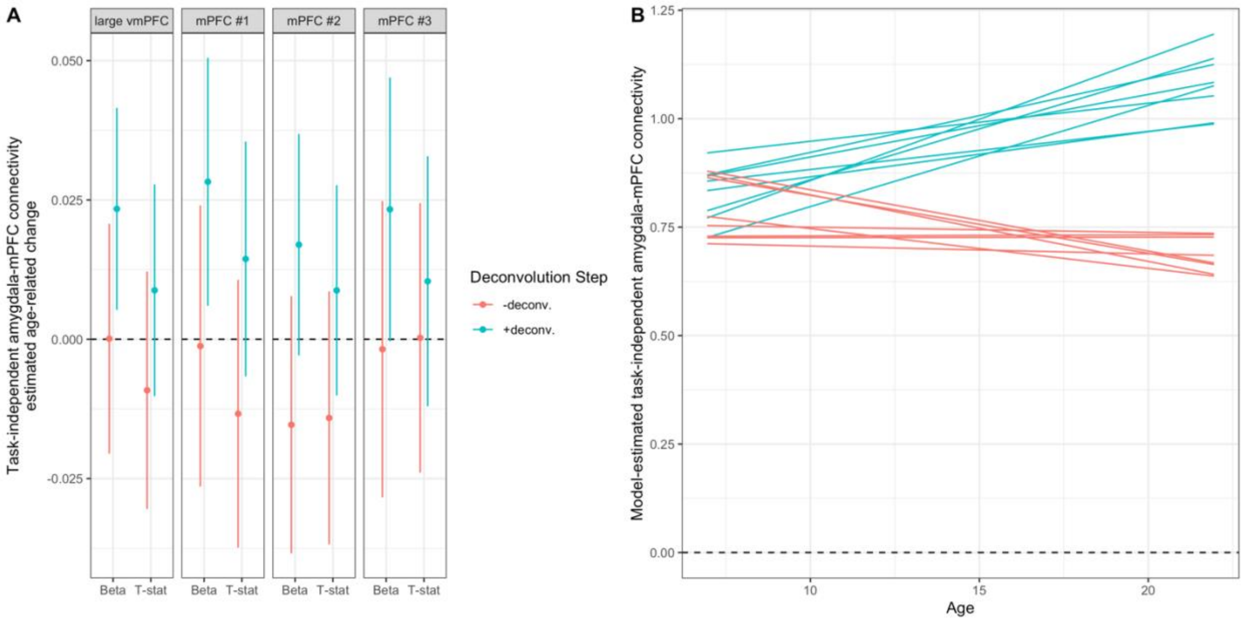


**sFigure 51:** Age-related change in task-independent amygdala—mPFC connectivity
Age-related change coefficients in task-independent amygdala—mPFC connectivity across methods and ROIs (A). The y axis indicates estimated linear age-related change in amygdala—mPFC connectivity. In (B), model predictions as a function of age are plotted as spaghettis in red for pipelines without deconvolution, and blue for pipelines with deconvolution.

### BSC Functional Connectivity Results

*Group mean amygdala—mPFC BSC*

To preserve computational resources, all models for mean amygdala—mPFC BSC were fit using the lme4 R package with intercepts allowed to vary by participant. For both the fear > baseline and neutral > baseline contrasts, we found positive amygdala—mPFC connectivity for all pipelines without a global signal correction, and positive (yet weaker) connectivity for all pipelines with a correction (sFigure 52). We did not find average differences between amygdala—mPFC BSC for the fear > neutral contrast (sFigure 52).

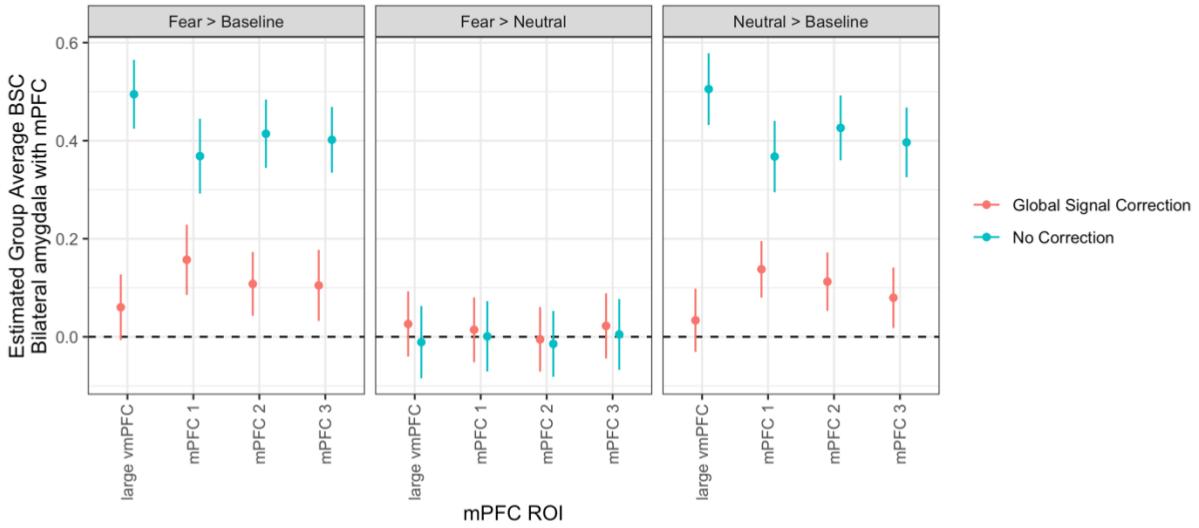


**sFigure 52:** Group mean amygdala—mPFC BSC across contrasts, mPFC ROIs, and pipelines Points represent mean estimates and error bars represent 95% confidence intervals.

*Multiverse analyses of age-related change in amygdala—mPFC BSC*

In addition to the constructing specification curves for age-related change in amygdala—mPFC BSC for the fear > baseline contrast as reported in the main manuscript (see Figure 5), we constructed parallel specification curves for the neutral > baseline and fear > neutral contrasts. For the neutral > baseline contrast, 77.9% of specifications found positive age-related change (most of them for mPFC ROI #1 or the large vmPFC ROI), though only 12.5% of posterior estimates excluded 0 (sFigure 53). For the fear > neutral contrast, only 1 pipeline out of 300 resulted in a posterior estimate excluding 0 for age-related change (sFigure 54).

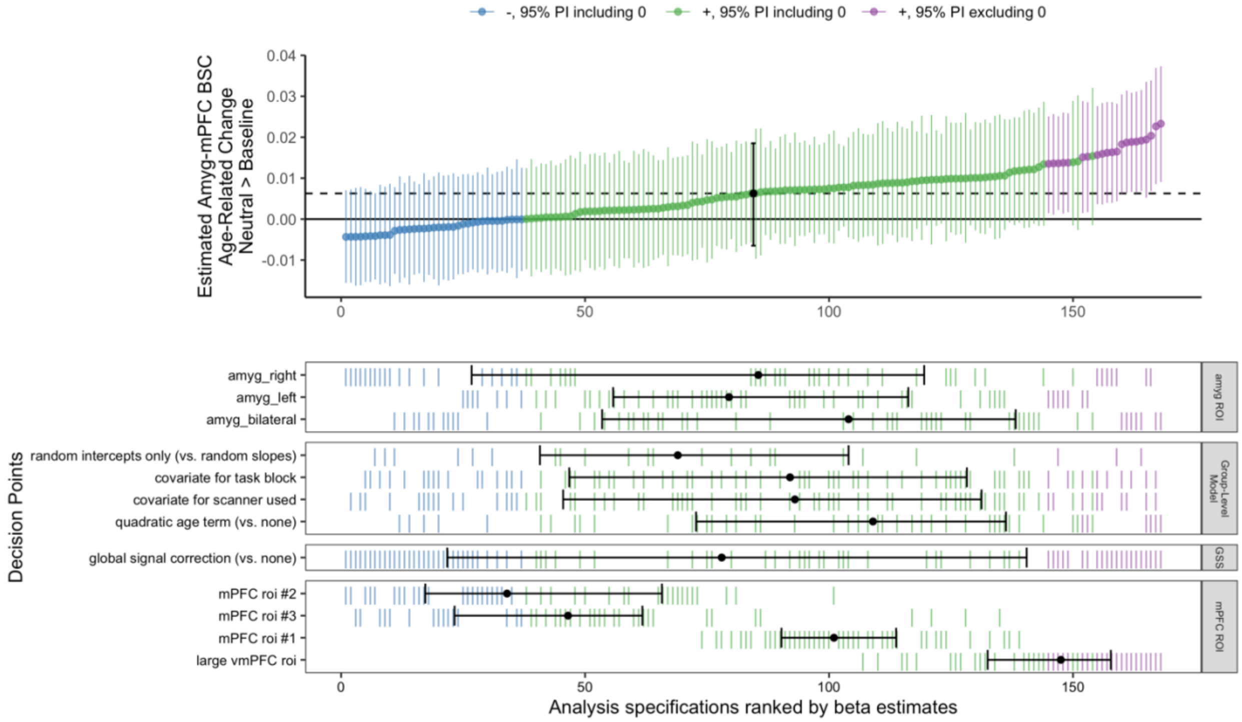


**sFigure 53:** Spec. curve for age-related change in neutral > baseline BSC
**A:** Points represent estimated linear age-related change in amygdala—mPFC BSC for each specification, and lines represent corresponding 95% posterior intervals. **B:** Variables on the y-axis represent analysis choices, corresponding lines indicate that a choice was made, and blank space indicates that the choice was not made in a given analysis.

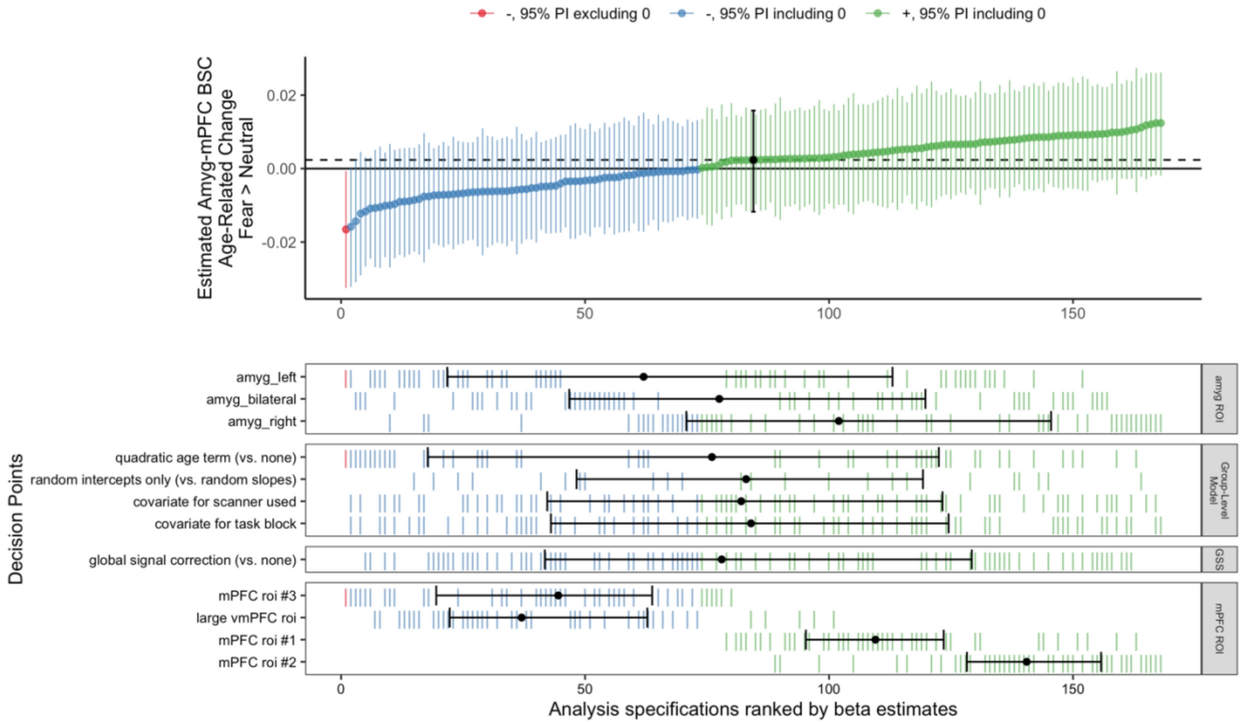


**sFigure 54:** Spec. curve for age-related change in fear > neutral BSC
**A:** Points represent estimated linear age-related change in amygdala—mPFC BSC for each specification, and lines represent corresponding 95% posterior intervals. **B:** Variables on the y-axis represent analysis choices, corresponding lines indicate that a choice was made, and blank space indicates that the choice was not made in a given analysis.

*Impacts of analysis choices on age-related change estimates for amygdala—mPFC BSC*

Unlike gPPI, for all BSC contrasts the choice of mPFC ROI, rather than preprocessing or modeling decisions, made biggest relative impact on estimates of age-related change (sFigures 55-57). For both fear > baseline and neutral > baseline contrasts, pipelines with mPFC ROIs #2-3 showed more negative age-related change compared to mPFC ROI #1 or the large vmPFC ROI. For the fear > neutral contrast, pipelines with mPFC ROI #3 and the large vmPFC ROI showed more negative age-related change compared to mPFC ROIs #1-2 (sFigure 57).

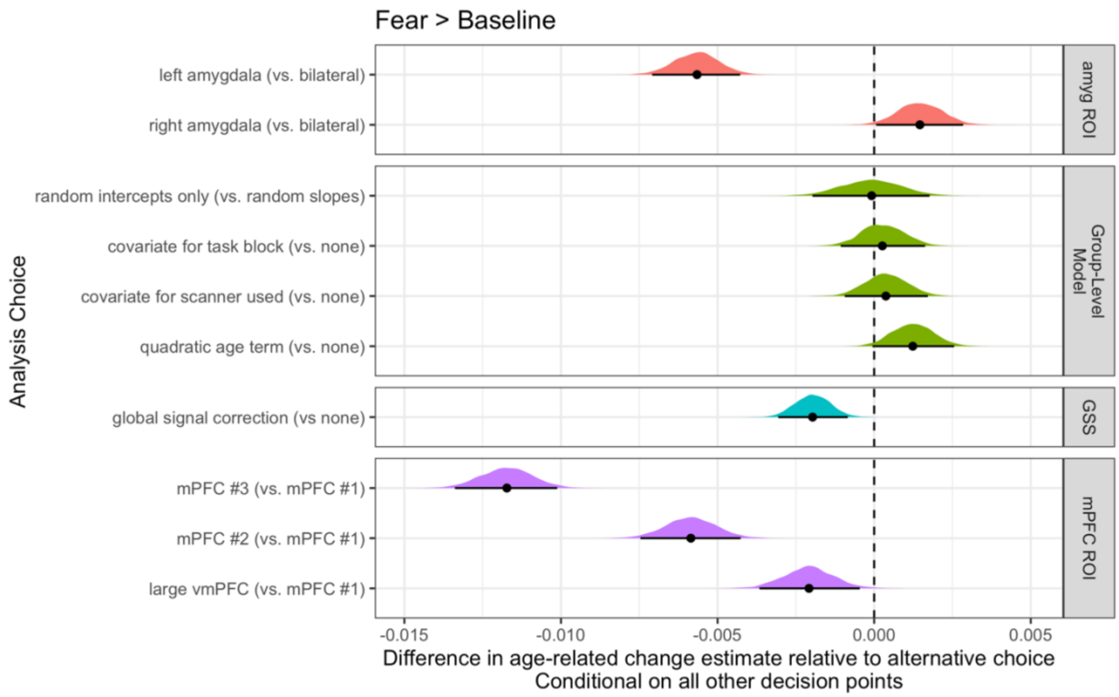


**sFigure 55:** Fork impacts on age-related change for fear > baseline amygdala—mPFC BSC Posterior distributions and 95% posterior intervals are shown, representing the average difference in linear age-related change estimates relative to the alternative choice.

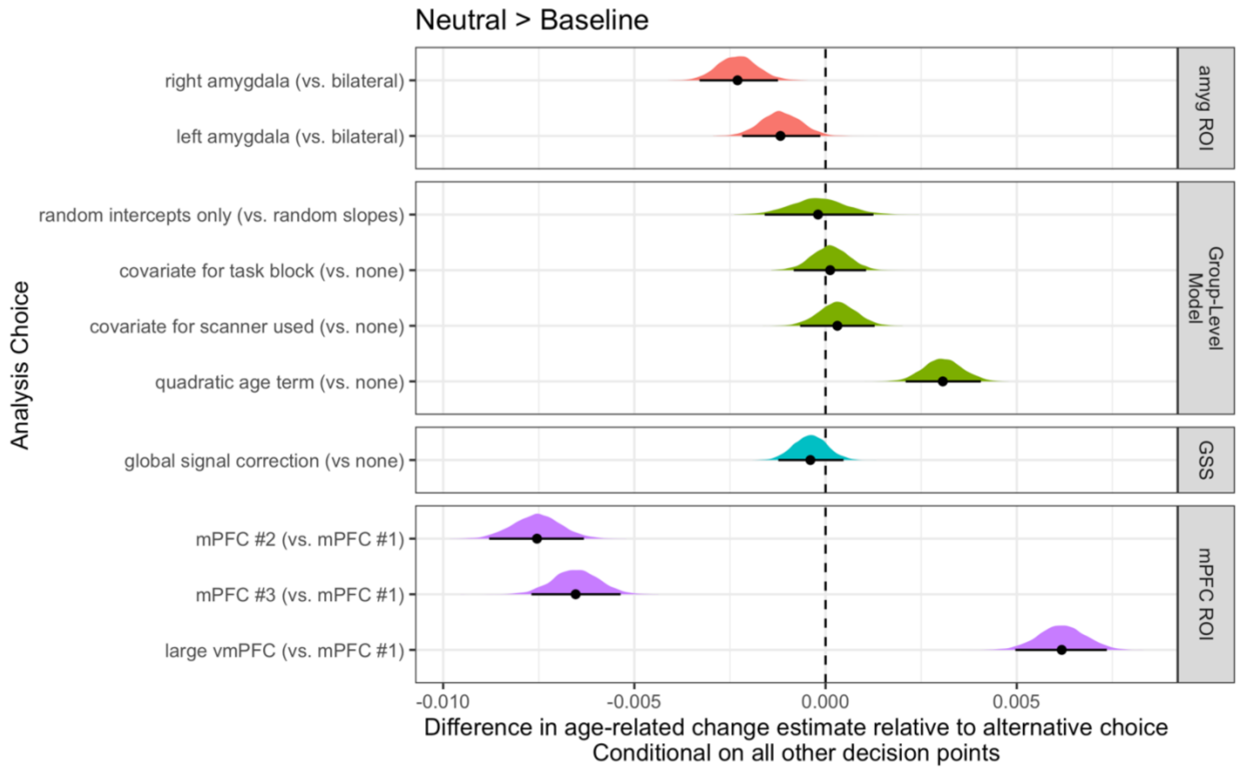

**sFigure 56:** Fork impacts on age-related change for neutral > baseline amygdala—mPFC BSC Posterior distributions and 95% posterior intervals are shown, representing the average difference in linear age-related change estimates relative to the alternative choice.

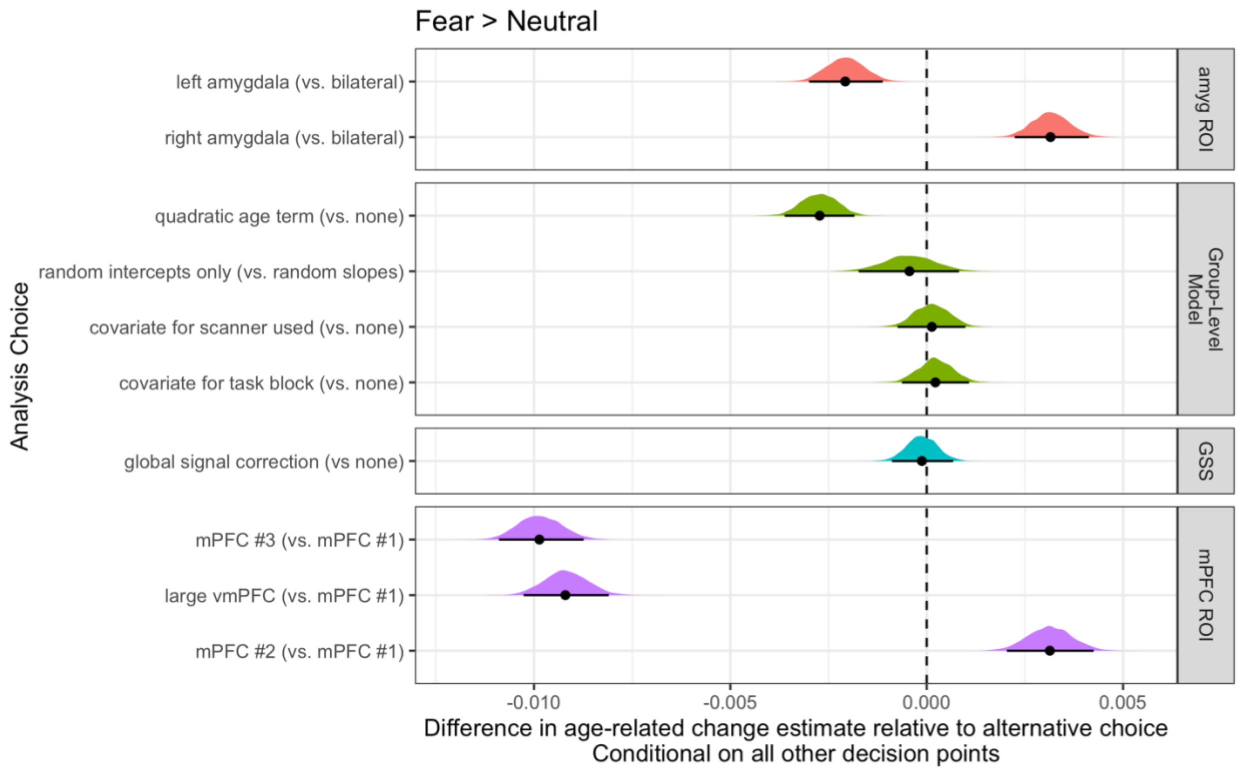


**sFigure 57.** Fork impacts on age-related change for fear > neutral amygdala—mPFC BSC Posterior distributions and 95% posterior intervals are shown, representing the average difference in linear age-related change estimates relative to the alternative choice.

*Nonlinear age-related changes in amygdala—mPFC BSC*

Some model specifications for amygdala—mPFC BSC included quadratic age-related change terms. Overall, however, we found little consistent evidence for quadratic age-related change for any contrast (see sFigures 58-60), as the sign of quadratic terms (‘peaks’ vs ‘troughs’) varied across specifications, and very few terms for each contrast were estimated with the 95% posterior interval excluding 0 (4% for fear > baseline, 8% for neutral > baseline, 18%

for fear > neutral). Inverse age models for age-related change in amygdala—mPFC BSC also indicated little evidence for consistent age-related change (sFigure 61).

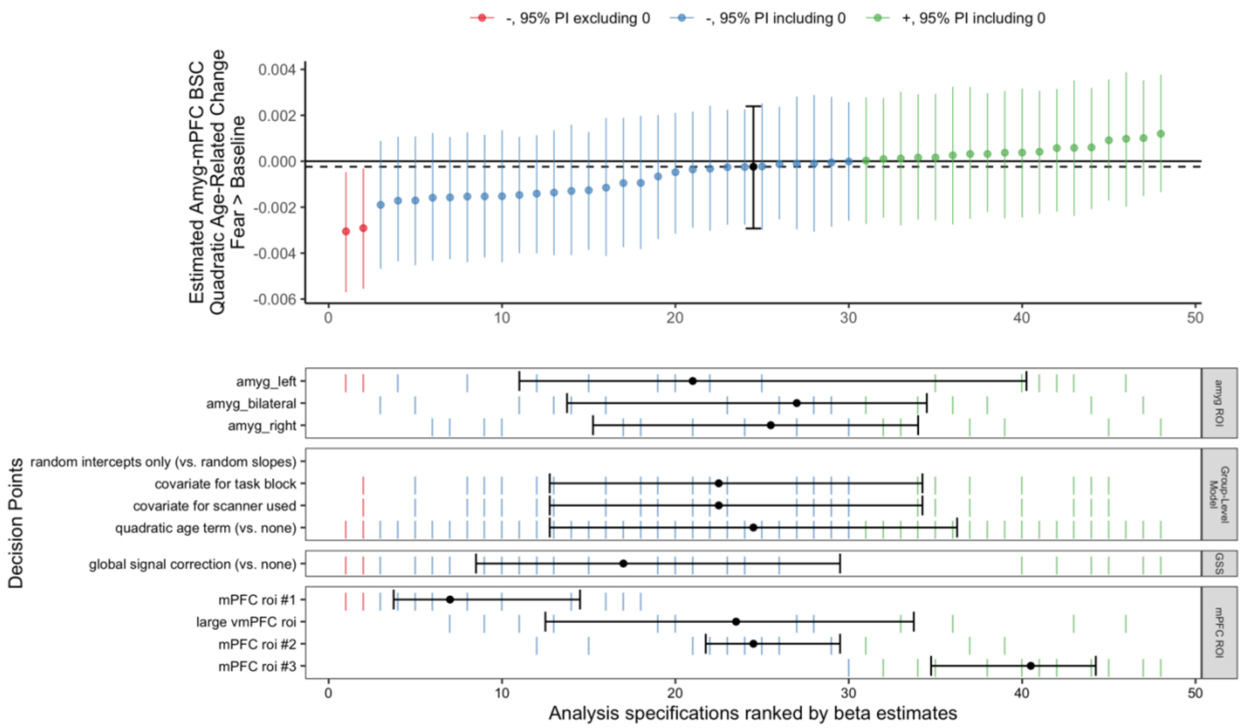


**sFigure 58:** Spec. curve for quadratic age-related changes in fear > baseline amygdala—mPFC BSC

**A:** Points represent estimated quadratic age-related change in amygdala—mPFC BSC for each specification, and lines represent corresponding 95% posterior intervals. **B:** Variables on the y-axis represent analysis choices, corresponding lines indicate that a choice was made, and blank space indicates that the choice was not made in a given analysis.

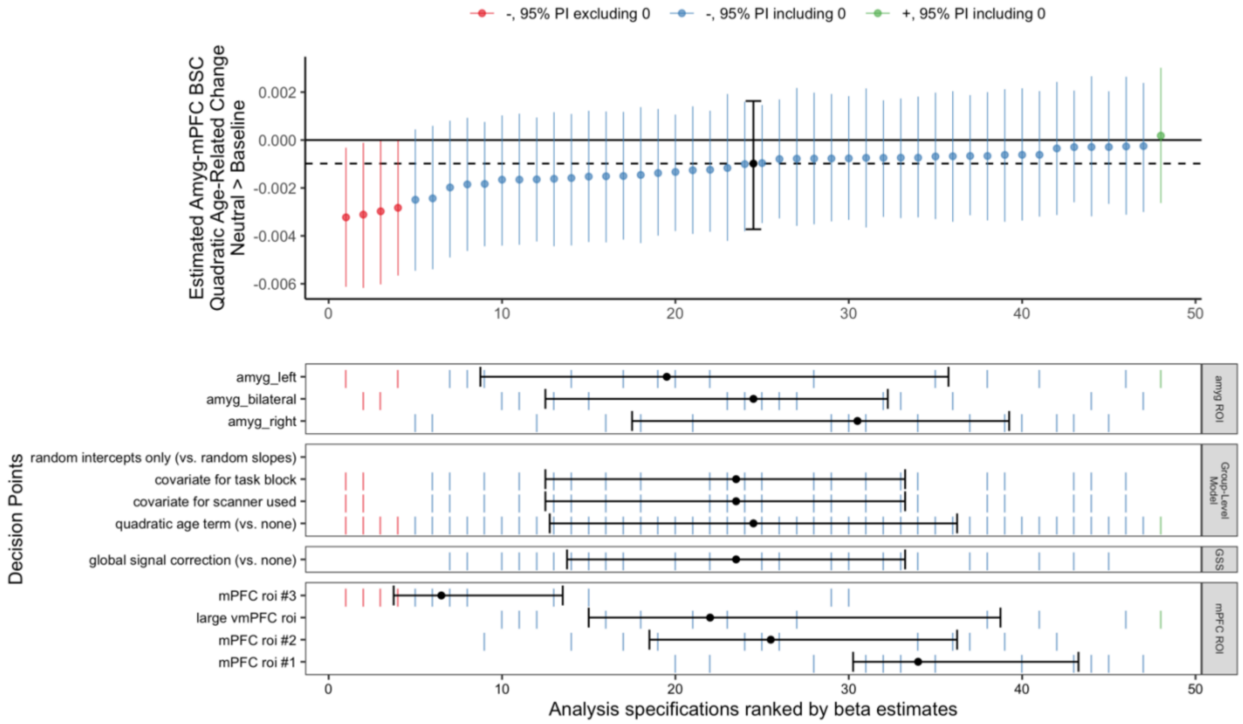


**sFigure 59:** Spec. curve for quadratic age-related changes in neutral>baseline amygdala—mPFC BSC
**A:** Points represent estimated quadratic age-related change in amygdala—mPFC BSC for each specification, and lines represent corresponding 95% posterior intervals. **B:** Variables on the y-axis represent analysis choices, corresponding lines indicate that a choice was made, and blank space indicates that the choice was not made in a given analysis.

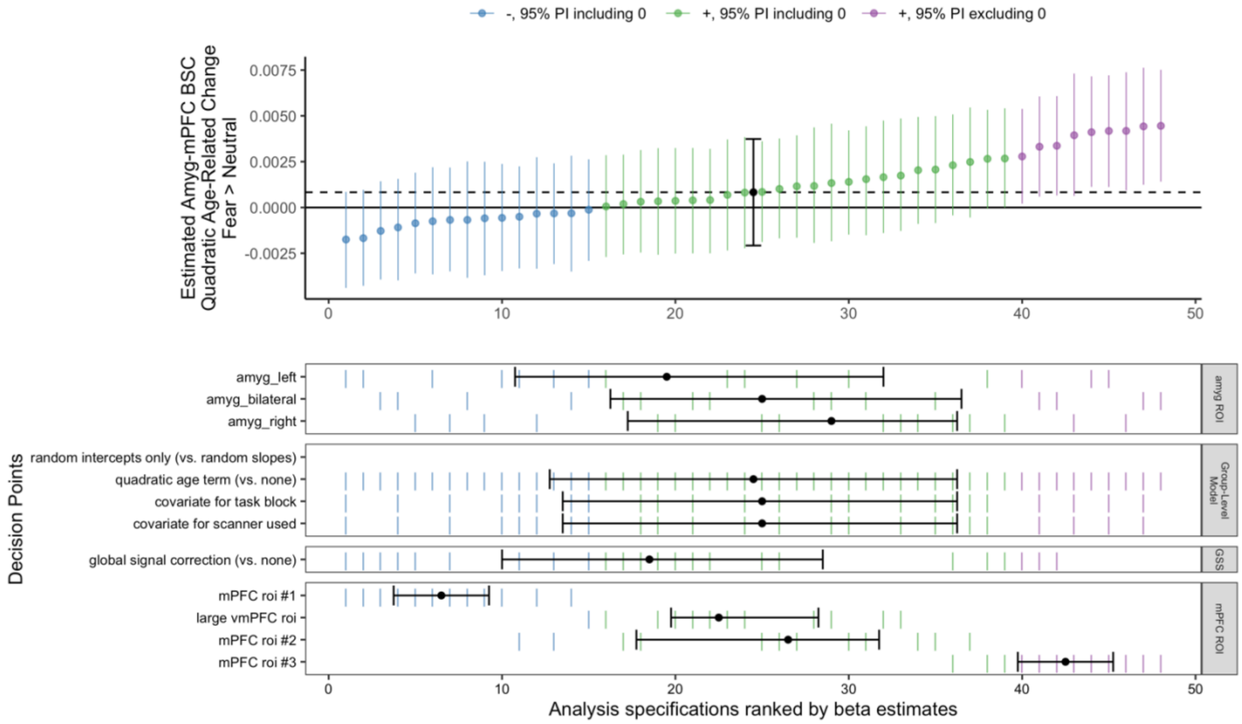


**sFigure 60:** Spec. curve for quadratic age-related changes in fear > neutral amygdala—mPFC BSC **A:** Points represent estimated quadratic age-related change in amygdala—mPFC BSC for each specification, and lines represent corresponding 95% posterior intervals. **B:** Variables on the y-axis represent analysis choices, corresponding lines indicate that a choice was made, and blank space indicates that the choice was not made in a given analysis.

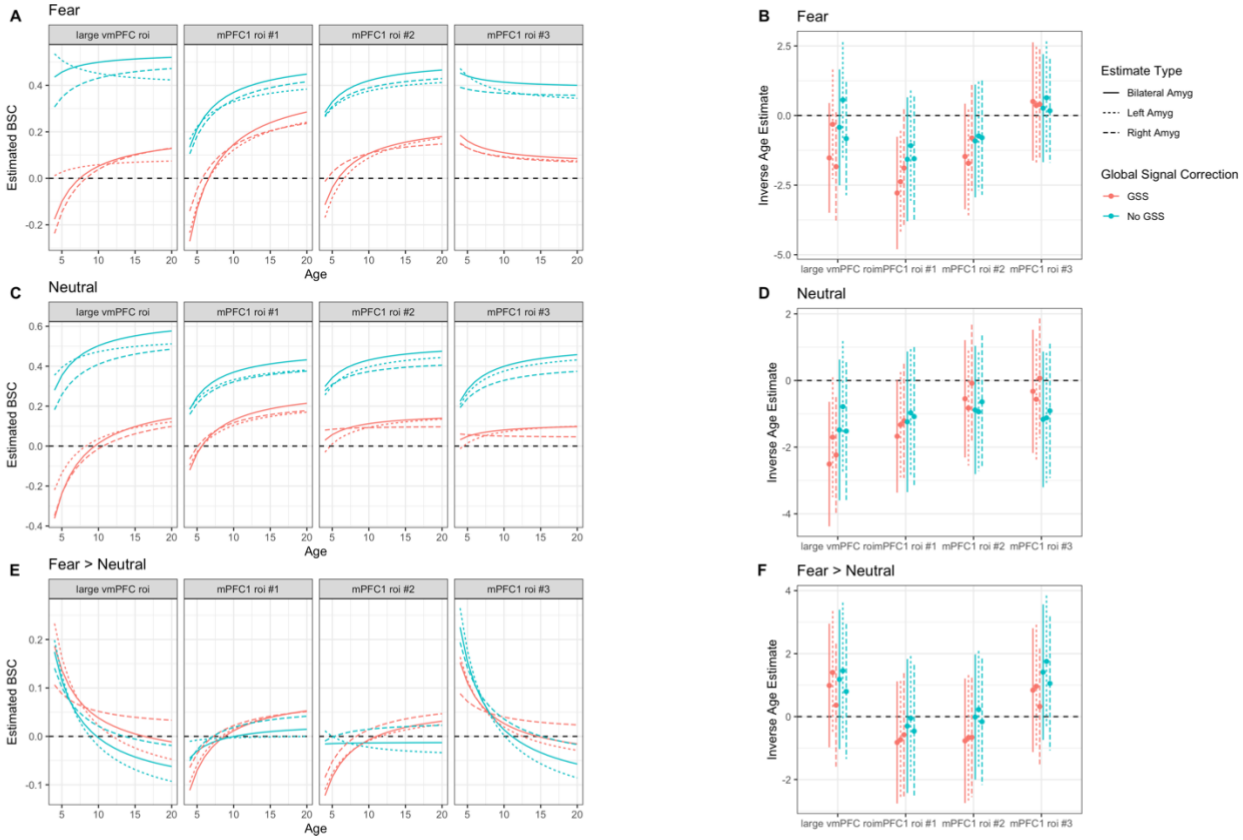


**sFigure 61:** Inverse age models for amygdala—mPFC BSC
Left panels show fitted model predictions for inverse age models for the fear (**A, top**), neutral (**C, middle**), and fear > neutral (**E, bottom**) contrasts. Specifications including a global signal correction are plotted in red, and without it in blue. Line type indicates amygdala ROI (solid = bilateral, dotted = left, dashed = right). Right panels show beta estimates for corresponding models for each contrast. Positive estimates for inverse age indicate decreases in amygdala—mPFC BSC as a function of age, and vice-versa.

*Correlations between head motion & amygdala—mPFC BSC estimates*

We calculated correlations between mean framewise displacement and BSC estimates for each ROI and contrast for pipelines with and without global signal correction. For pipelines without global signal correction fear > baseline and neutral > baseline estimates were positively associated with head motion (see sFigure 62). Such correlations were reduced in pipelines with a global signal correction, consistent with indications that such estimates of BSC for only one condition may also represent some ‘task-independent’ signal that may contain motion and respiratory artifacts. BSC estimates for the fear > neutral contrast were not overall strongly associated with head motion.

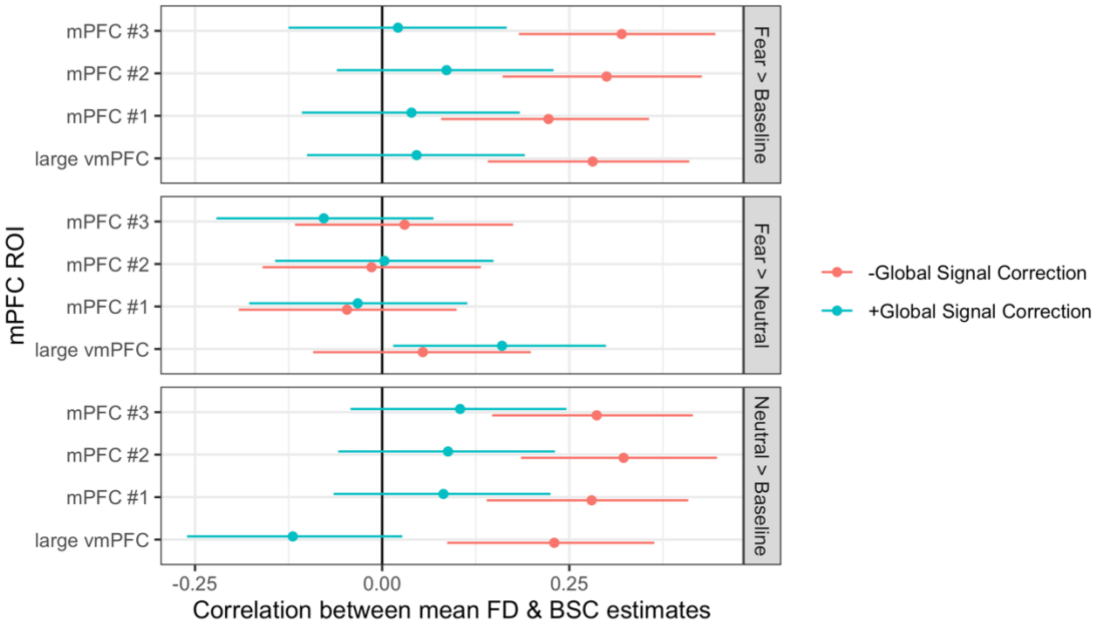


**sFigure 62:** Correlations between mean FD & BSC estimates across scans
For each contrast and ROI, points show estimated product-moment correlations and error bars represent 95% confidence intervals.

*Within-scan similarity between gPPI & BSC amygdala FC*

Previous work from Di et al. (2020) indicated a convergence in BSC and gPPI for contrasts between two task conditions. We addressed this in the present data with a within-scan analysis.To examine whether certain pipelines across BSC and gPPI were representing more similar signals, we computed correlations between vectors of fear > neutral amygdala FC with the rest of the brain between each pair of BSC and gPPI pipelines for the same scan. Overall, patterns of amygdala FC with the rest of the brain were positively associated between all pipelines on average, although not strongly, and associations varied widely across participants (sFigure 63, left panel). gPPI pipelines without deconvolution resulted in somewhat more similar amygdala connectivity patterns with BSC relative to gPPI pipelines with deconvolution.

Amygdala connectivity patterns were the most strongly associated between the two BSC pipelines with versus without global signal correction (sFigure 63, right panel).

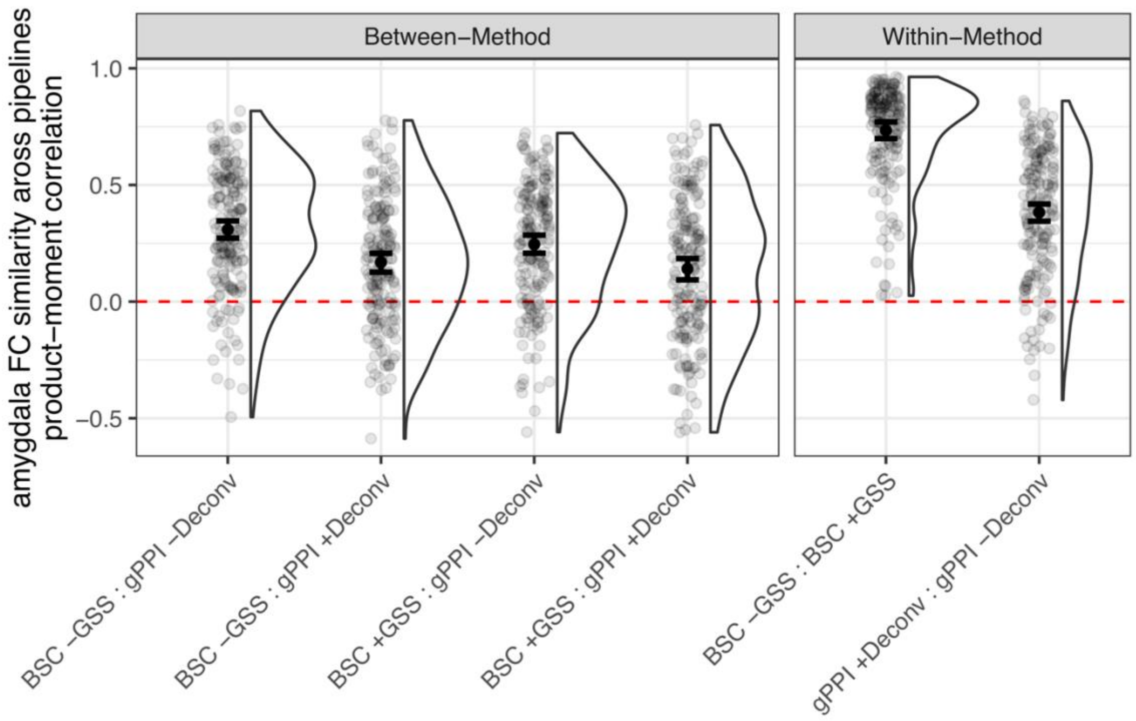


**sFigure 63.** Similarity of amygdala FC with the rest of the brain between gPPI & BSC
Points represent similarity across pipelines for each individual scan for each comparison (x axis marks each comparison of pipelines), with error bars summarizing 95% posterior intervals for estimated mean similarity across scans. **Left:** between-method comparisons of similarity of amygdala FC between BSC and gPPI methods. **Right:** within-method comparisons of similarity of amygdala FC within BSC and gPPI methods, respectively, while altering decisions for GSS (for BSC) and deconvolution (gPPI). GSS = global signal subtraction using post-hoc mean centering.

*Between-scan correlations between gPPI & BSC estimates*

We also sought to examine whether different methods for functional connectivity yielded similar relationships between scans in mean amygdala—mPFC functional connectivity estimates. We computed between-scan rank-order correlations between gPPI and BSC estimates for each contrast to examine whether relative ordering was preserved across different estimates of amygdala—mPFC functional connectivity. For each pair of preprocessing pipelines, we computed the rank-order correlation between vectors of functional connectivity estimates (1 datapoint per scan per pipeline). Most generally, BSC estimates were not strongly associated with gPPI estimates, even for the same contrast and mPFC ROI (sFigures 64-66). However, for the fear > neutral contrast specifically, gPPI estimates without deconvolution were generally positively correlated with BSC estimates for corresponding ROIs, while gPPI estimates with deconvolution were not (sFigure 66). In addition, while BSC estimates for the same ROI with versus without global signal correction were generally highly associated, gPPI estimates for the same ROI with versus without deconvolution were often only weakly associated.

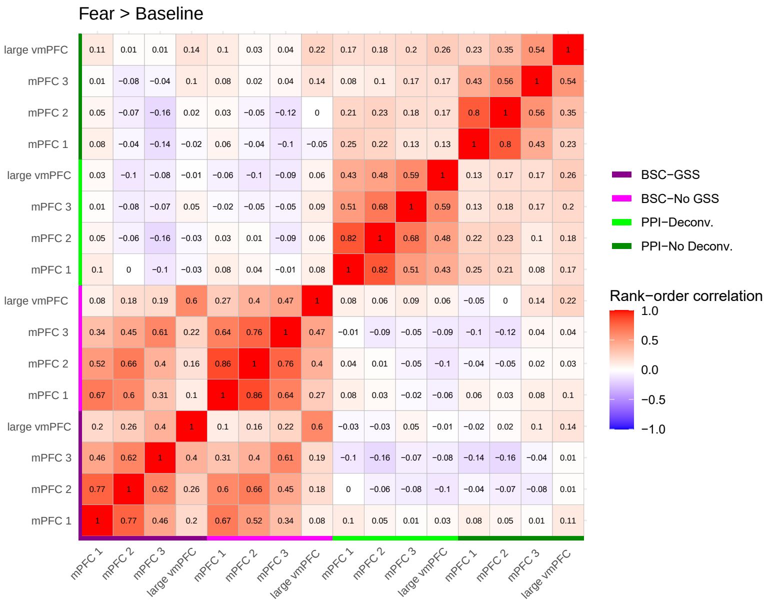

**sFigure 64:** Between-scan correlations between fear > baseline gPPI & BSC estimates

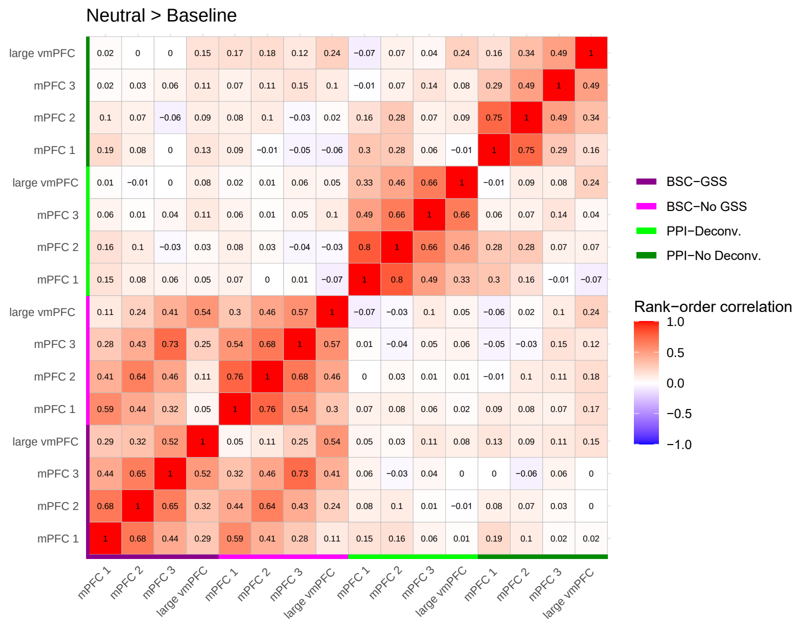

**sFigure 65:** Between-scan correlations between neutral > baseline gPPI & BSC estimates

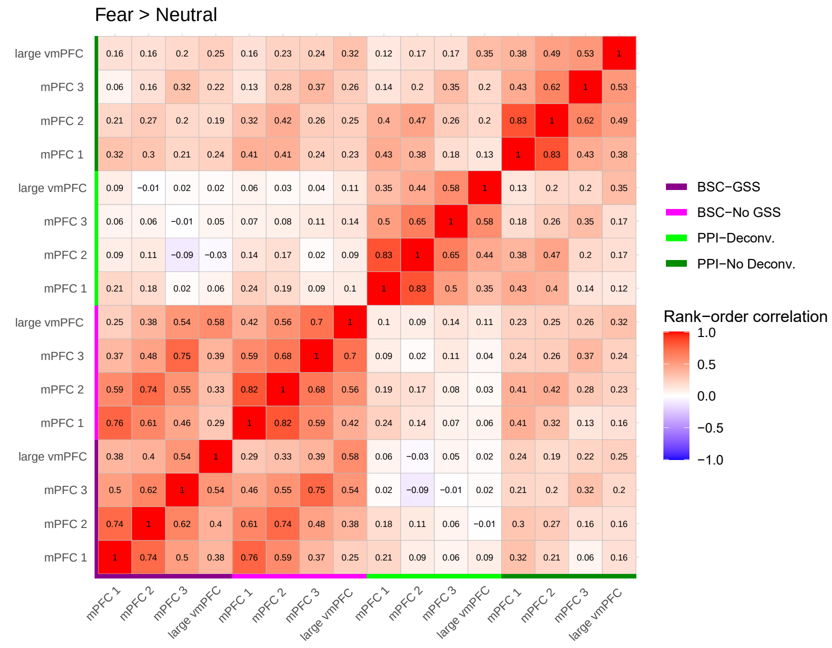


**sFigure 66:** Between-scan correlations between fear > neutral gPPI & BSC estimates

Associations with generalized anxiety and social anxiety behaviors

Here, we primarily focused on associations between amygdala—mPFC responses and separation anxiety in efforts to follow up on previous analyses of the same cohort (Gee et al., 2013), and because separation anxiety behaviors often show pronounced decline among typically developing cohorts of age range (Allen et al., 2010; Francis et al., 1987). However, we also examined associations with parent-reported generalized anxiety and social anxiety behaviors as measured by the SCARED-P and RCADS-P, as well as the Total Anxiety Score from the RCADS-P. Using multilevel linear regression models (covarying for age) with maximum likelihood estimation as described above, we did not find robust evidence for longitudinal associations between amygdala—mPFC measures and any of the anxiety scales examined (sFigure 67).

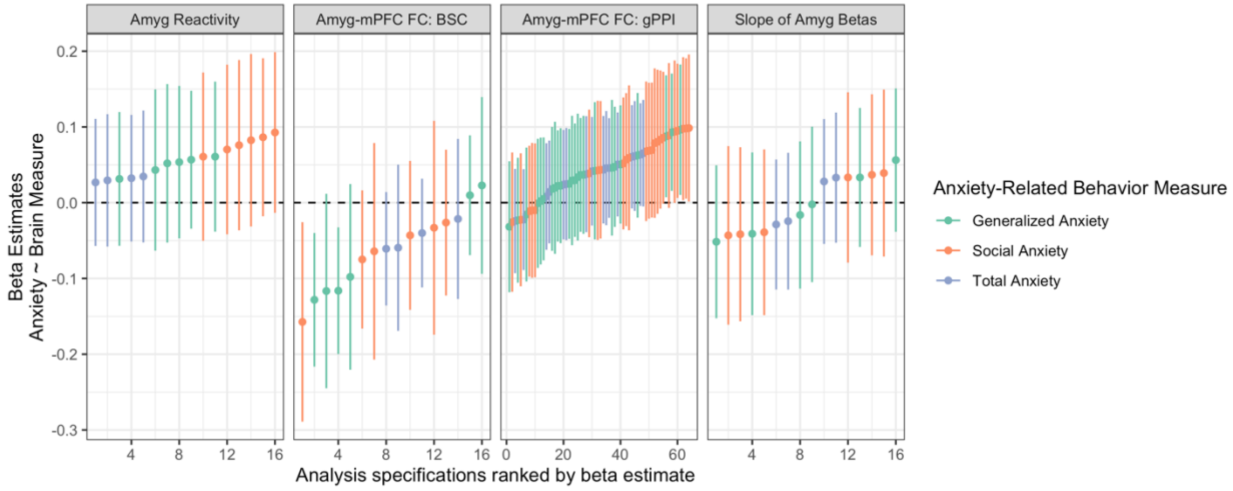


**sFigure 67:** Associations between amygdala—mPFC measures and anxiety-related behaviors Specification curves are shown for longitudinal associations between fear > baseline amygdala reactivity (A), fear > baseline amygdala—mPFC BSC (B), fear > baseline amygdala—mPFC gPPI (C), and slopes for amygdala fear betas (D) and anxiety scales (green = generalized anxiety behaviors, orange = social anxiety behaviors, purple = total anxiety behaviors).
